# MRMhub: one-stop solution for automated processing of large-scale targeted metabolomics data

**DOI:** 10.64898/2025.12.20.695370

**Authors:** Bo Burla, Guoshou Teo, Peter I. Benke, Zinan Lu, Sock Hwee Tan, Shanshan Ji, Jeongah Oh, Pei Yen Lim, Vaitheeswari, Amaury Cazenave Gassiot, Kavita Venkataraman, E Shyong Tai, Federico Torta, Markus R. Wenk, Mark Y.Y. Chan, Hyungwon Choi

**Affiliations:** Singapore Lipidomics Incubator, Life Sciences Institute, National University of Singapore, Singapore; Department of Medicine, Yong Loo Lin School of Medicine, National University of Singapore, Singapore; Department of Biochemistry, Yong Loo Lin School of Medicine, National University of Singapore, Singapore; Cardiovascular Disease National Collaborative Enterprise, Consortium for Clinical Research Innovation, Singapore; Saw Swee Hock School of Public Health, National University of Singapore; Signature Research Programme in Cardiovascular & Metabolic Disorders, Duke-NUS Medical School, Singapore; College of Health and Life Sciences, Hamad Bin Khalifa University, Doha, Qatar; Department of Cardiology, National University Heart Centre, Singapore

## Abstract

Data processing and quality control are essential for complex targeted mass spectrometry (MS) assays in large-scale metabolomics studies. However, existing software solutions have significant gaps in robustness and scalability. We report MRMhub, a one-stop solution for streamlined processing of large-scale targeted MS data. MRMhub consists of a novel peak integration engine with unique algorithmic design to address the scalability challenge and a comprehensive collection of post-acquisition data processing and analytical quality control tools. The ensemble facilitates rapid and consistent processing of complex chromatograms, quantification, drift and batch correction, quality assessment and control, feature filtering, and data/workflow sharing and reporting. MRMhub can process data from highly complex assays in population-scale studies within minutes, with full digital footprints warranting reproducibility and traceability. We demonstrate its performance using two large-scale lipidomics data sets. We distribute the source code and data sets freely for community development.

## Introduction

Liquid chromatography (LC) coupled to tandem mass spectrometry (MS) is the leading platform for quantitative analysis of small molecules. Targeted approaches are typically based on multiple reaction monitoring (LC–MRM), with methods optimized for specific analyte classes such as metabolites and lipids^1, 2^. MS-based assays are increasingly used in large-scale applications involving thousands of biological samples or more^3–11^, and these experiments require weeks to months of LC-MS analysis time. The workflow poses significant challenges from an operational point of view because of the difficulty in executing sample preparation protocols without major sources of variation and maintaining LC-MS system in stable working conditions during the long analysis period (**Fig. 1a**). Therefore, these experiments are carefully planned with safeguards such as randomization of sample analysis sequence, periodic injection of quality control samples^12, 13^ and use of internal standards (ISTD) to allow for post-analysis correction of experimental artifacts and time-dependent variation.

**Fig. 1.**
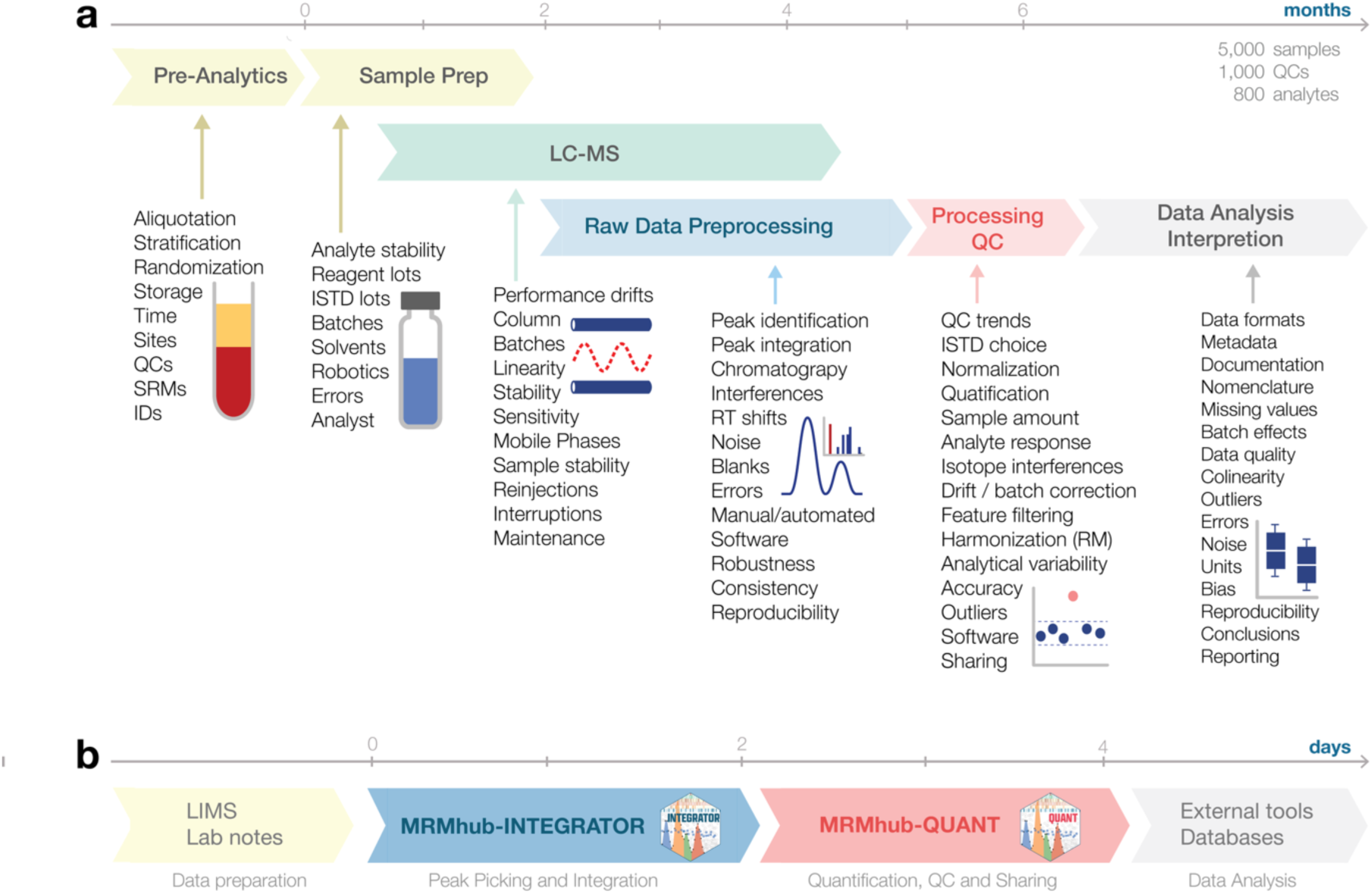
Workflow and challenges of a targeted quantitative metabolomics/lipidomics analysis. **a.** Key aspects that are required, contribute, or may bias the final results. Key aspects that are required, contribute, or may bias the final results. The timeline reflects the approximate throughput achievable with a robotic pipetting system, a single LC-MS instrument with an analysis time of 15 minutes per sample, and data processing using vendor software and spreadsheets. **b.** MRMhub data processing pipeline, its core modules with their key functions, and approximate timeline to process a dataset from above using this pipeline.

In addition to the challenges with laboratory operation, data processing can have an equally significant impact on analysis throughput. In metabolomics and particularly in lipidomics, individual MRM ion traces may comprise multiple peaks from isomeric and isobaric compounds, adducts, isotopologues, and in-source fragmentation products that may be partially or fully overlapping^14, 15^. Processing complex ion chromatograms thus requires expertise of well-trained analysts. In large-scale studies targeting hundreds of analytes in hundreds to thousands of samples, peak integration is a tedious, time-consuming and technically daunting task. Although a few software solutions provide a level of automation, manual adjustments to peak picking and feature annotation are unavoidable in practice. The amount of work required for data inspection increases with the complexity of assay and the scale of experiment, and human interventions may not remain consistent and reproducible throughout the time-consuming process.

Specifically, the existing tools can be improved in regard to two major aspects^16–19^. First, the algorithms implemented in the existing peak integration tools do not borrow information across injections in terms of integration borders and retention time (RT) drift. In any target analyte, if the MRM window features chromatograms with multiple modes and thus the overlapping ion trace needs to be deconvoluted into individual ion chromatograms, this task cannot be easily coordinated across different samples, and the results may thus be inconsistent. Second, with large number of analytes packed in a single LC-MRM method, some tools run into scalability issues in standard desktop hardware as the number of analysis files grows large, limiting their applicability. Our benchmarking exercise using population-scale data, as narrated later, collectively points to software architecture issues as the root cause of this problem, specifically on-memory storage of raw data and many other pre-processing requirements. These limitations undermine software performance, data management and visualization options in large-scale studies.

In addition, downstream data processing steps, including the derivation of concentration and the calculation of quality metrics, are just as important as peak integration in large-scale studies. As we demonstrate later, data normalization approaches using lipid class-specific ISTDs and quality control (QC) samples do not address all time-dependent variations and analytical artifacts visible to the analyst, potentially introducing their own bias and artifacts. Hence specific data processing and quality assurance (QA) or QC guidelines for MS-based metabolomics methods have been proposed^20, 21^. However, we were not able to find a single open-source tool that offers a full suite of functionalities implementing these guidelines^22–27^.

In this paper, we report an integrated solution for processing large-scale MRM data sets, called MRMhub (**Fig. 1b**). MRMhub consists of two interconnected modules: (i) INTEGRATOR for highly automated and fast processing of complex MRM chromatograms, and (ii) QUANT for comprehensive post-acquisition data processing and quality control of datasets. These tools can be used jointly or independently. We demonstrate the performance of our novel software solution using three datasets from published studies (see **Methods**). We use SPERFECT^28^ study as the main example dataset (937 samples, 503 transitions) in this manuscript (**Dataset 1**). We also use data from a failed analysis batch from the DYNAMO study^9^ for RT drift estimation (**Dataset 2**), and lastly, we demonstrate the outstanding scalability of MRMhub using the main dataset (4,591 samples, 829 transitions) of the DYNAMO study (**Dataset 3**).

## Results

### INTEGRATOR module

We first introduce a novel peak integration module, termed INTEGRATOR. The module has evolved from an existing automated engine, originally developed as MRMkit^29^. As we demonstrate below, new functionalities bring significant improvement to peak integration performance, such as automatic retention time (RT) drift correction, synchronized feature integration and enhanced customizability. The module carries out peak integration in four steps (**Figs. 2a–d**): (1) data and metadata import and validation, (2) estimation of transition-specific RT drifts, (3) cross-sample consensus feature detection, and (4) refinement of peak integration borders. Once the borders are set, the algorithm completes one cycle of automated integration and presents the results through visualization (**Fig. 2e–f**). After inspecting the result, the user can refine the integration outcome by changing custom parameters for each analyte and re-perform integration iteratively (**Fig. 2g**) or refine integration bounds case by case using RT table, i.e. for each sample manually (**Fig. 2h**) in the final cycle.

**Fig. 2.**
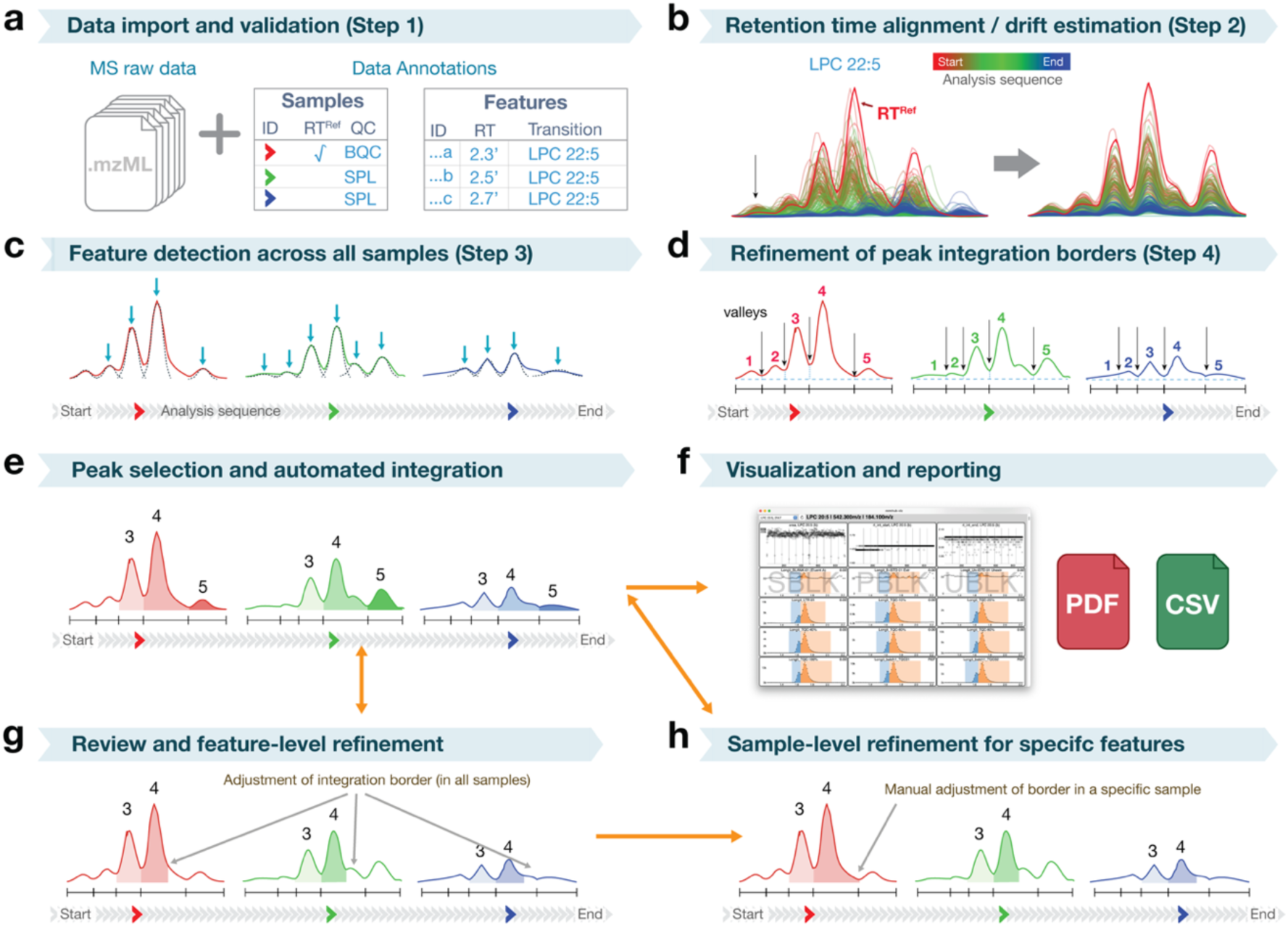
Operations of the INTEGRATOR module. **a**. The algorithm first validates input MS data and meta data files. **b.** It next estimates RT drift for each analyte, aligns ion traces across samples and determines shared patterns. **c.** Subsequently, it surveys the peaks across the samples and determines the number of peaks. **d.** In the next step, the algorithm determines integration borders accounting for the RT drift in each sample. **e.** Lastly, the algorithm integrates the peaks defined in the feature list. **f-g**. The user annotates the peaks of interest, performs automated integration iteratively with the help of ion chromatogram viewer app, and reports the integration results. **h**. The user can also adjust peak integration borders case by case using RT matrix file if such a process is necessary. The example shown in **a-h** is the complex case of lysophosphatidylcholine (LPC) 22:5 comprising three isomer peaks, and two isotopic interference peaks from LPC 22:6.

We describe the INTEGRATOR algorithm in more detail. INTEGRATOR imports platform-independent mzML files as input^30^, supporting all major LC-MRM platforms. It imports the following metadata: (i) a table defining MRM transitions and feature-specific integration settings, and (ii) a sample list specifying analysis order, sample types, and designated reference samples for RT drift estimation. INTEGRATOR validates the integrity of the metadata against the mzML files, flagging mismatches. Preparation of these files encourages sound practice of digitalizing the entire experiment.

Subsequently, the module estimates RT drift of each analyte from the first sample injected to the last, where the drift is calculated using temporal cross-correlation of entire chromatograms relative to designated time reference samples (**Supplementary Fig. 1**). Drift correction ensures accurate peak annotation, especially when using narrow peak detection windows and fixed/cross-sample consensus integration borders. RTs were mostly stable in **Datasets 1** and **3**, with average shifts ≤ 0.01 and ≤ 0.02 min, respectively, except in the last batch of **Dataset 3** where shifts increased up to 0.04 min. **Dataset 2** displayed spurious RT shifts up to 0.5 min, thereby exceeding one peak width. INTEGRATOR correctly estimated RT drifts even in these extreme cases, enabling accurate RT correction (**Figs. 3a and 3b**). Without such correction, peak annotation in the raw ion traces would have led to errors, due to peaks shifting into the initial windows of other features (**Fig. 3b**). INTEGRATOR reports RT drift estimates in visualizations and exported datasets.

**Fig. 3.**
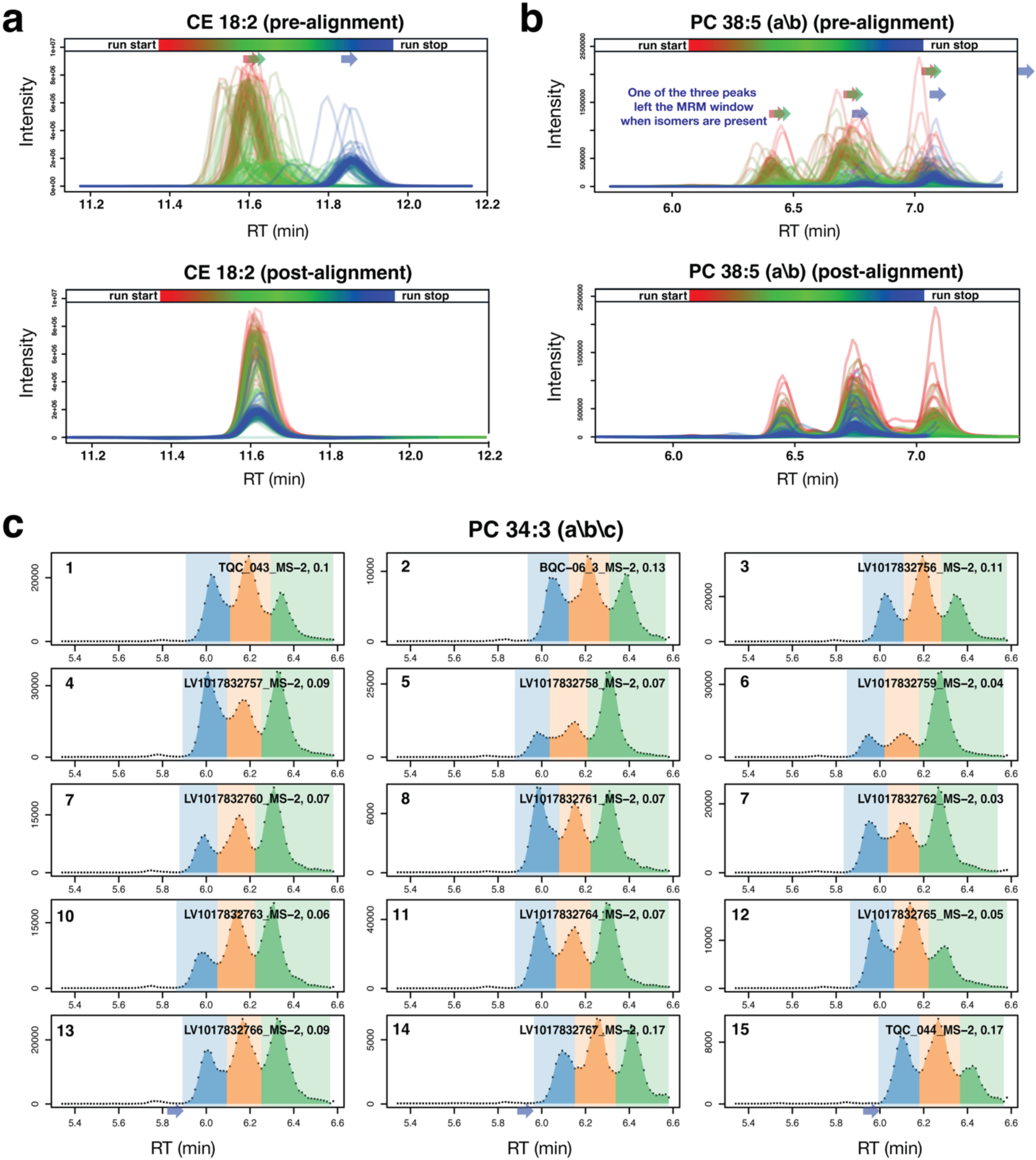
Peak integration of complex chromatograms. **a.** Ion traces of CE 18:2 drifted by approximately 15 seconds during the analysis in **Dataset 2** (failed batch). RT alignment algorithm of INTEGRATOR flawlessly realigns the ion traces. **b.** The same alignment algorithm works for PC 38:5, which showed multiple ion traces with similar drift patterns. **c**. INTEGRATOR identifies the integration borders consistently throughout the analysis sequence (numbered), irrespective of their relative abundance levels or RT drift.

INTEGRATOR controls peak finding by setting the expected RT for a target peak and defining a global detection window (**Fig. 2e**). The module regulates its peak integration behaviour by setting allowable peak width ranges. Overall, peak shapes in **Dataset 1** exhibited gradually increased tailing, therefore we set the global maximum right border width larger than that of **Dataset 3**. INTEGRATOR accurately performed automated integration of single, unimodal and resolved peaks, despite the presence of adjacent peaks of varying intensities. (**Supplementary Figs. 2a, 2b and 2c**).

We observed equivalent performance for the integration of multiple partially resolved peaks with varying intensities from single ion traces, without requiring feature-specific settings beyond the expected RT (**Supplementary Fig. 2d**). Another unique feature of INTEGRATOR, aside RT shift correction, is its sensitivity, robustness and consistency in automatically determining integration borders for partially overlapping peaks across large number of samples, which occurs commonly in lipidomics assays using reversed phase LC (**Supplementary Fig. 2e**). In the challenging cases such as **Dataset 3** with persistent and spurious RT shifts, INTEGRATOR correctly and consistently performed peak detection and integration (**Fig. 3c**).

While automated determination of integration borders is efficient most of the time, manual adjustments for shared borders are still necessary to achieve consistent integration across samples. For example, INTEGRATOR allows the user to intentionally co-integrate peaks by manually setting the left and right borders of a peak group (**Supplementary Fig. 2f**). This is useful for combining unresolved peaks or isomer peaks at the integration stage. INTEGRATOR also adjusts manually set borders during RT shift correction, allowing for their accurate use even in datasets with severe RT drifts. For blank samples, where clear peaks are often absent, integration boundaries are learned from samples and QC materials (**Supplementary Figs. 2g-2h**).

Finally, INTEGRATOR enables specification of peak borders for individual analytes and samples using a RT table at the final stage of peak integration process (**Figs. 2g and 2h**). To facilitate this process, we provide a dedicated viewer app, MRMhub-viz, with an alternative option to export results to PDF files (see **Methods**). Among the 858 features in **Dataset 3**, we applied fixed borders to 39, and alternate peak widths for another 68. No individual sample-level RT modification was performed or deemed necessary.

Together, the flexibility of INTEGRATOR enables robust processing of complex chromatograms with a reasonable balance between full-scale automation and manual, feature-wide intervention. We provide full peak integration results for all three aforementioned datasets (see **Data Availability**).

### QUANT module

QUANT is an R package for end-to-end post-processing of targeted small-molecule MS data, i.e. from peak areas to concentrations **(Supplementary Fig. 3)**. QUANT serves as a library for writing scripts to process individual dataset or to develop custom workflows, while it also offers intuitive functions accessible to lab analysts with basic programming experience. It supports diverse analytical designs from semi-quantitative metabolomics and lipidomics analysis to fully quantitative assays in clinical and environmental applications and is scalable to large-scale studies.

#### Assessment of analysis structure and detection of outliers

The module processes integrated peak areas from INTEGRATOR or external sources. It can import metadata describing samples, features, and other analysis parameters from various data formats. As the first step, QUANT similarly verifies data integrity and completeness, reducing inconsistencies that can be common in complex collaborative analyses. We applied the QUANT workflow to **Datasets 1 and 3**. Unless otherwise stated, **Dataset 1** was used as the main example below. Computational notebooks containing the code, corresponding results, and rendered reports are provided as **Supplementary Information** and are also available as HTML reports (see **Data Availability**).

In the first step, QUANT produces a sample sequence plot to visualize the overall analysis structure, including timeline, batch structure, QC samples, sample stratification and interruptions. The visualization provides a global view of the analysis plan **(Fig. 4a)**. It also produces relative log abundance (RLA) plots, revealing systematic effects on feature abundances as they may indicate changes or artifacts in sample preparation or instrument performance **(Fig. 4b)**. The RLA plots facilitate outlier detection and allow the user to exclude technical outliers from further processing. In **Dataset 1**, one sample type showed approximately two-fold reduction in the signals of endogenous analytes and spiked-in internal standards (ISTD), which was subsequently traced to the use of half the injection volume in this case (**Fig. 4b and Supplementary Figs. 4a and 4b**). The RLA plots also revealed a study sample with very low analyte but normal ISTD signals, suggesting an error in pipetting the sample during extraction (**Supplementary Figs. 4c and 4d**).

**Fig. 4.**
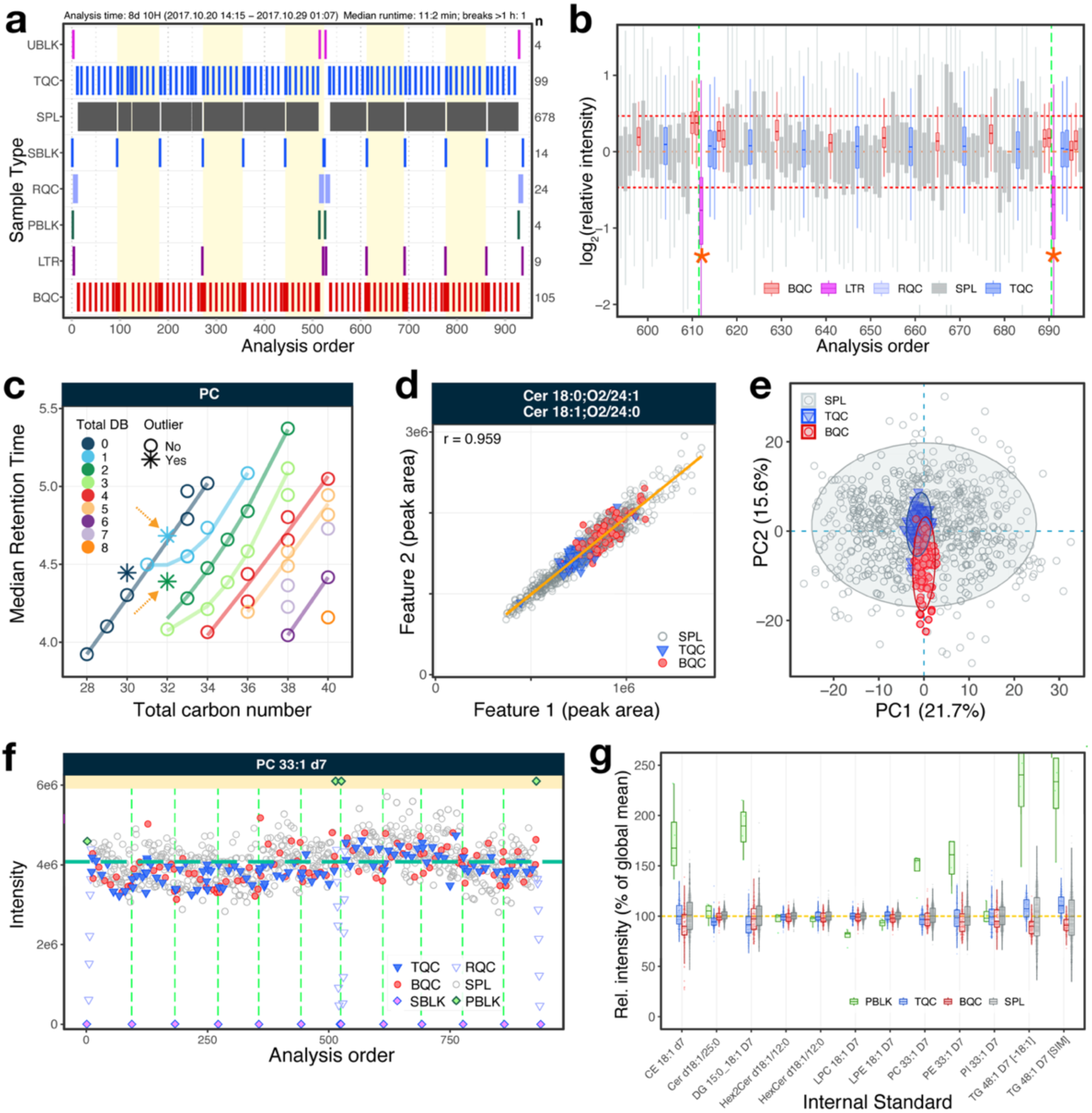
Analysis overview and peak annotation. **a**. Analysis design with frequency and placement of QC samples. See **Methods** for more details on individual plots and the QC sample types. **b.** Relative Log abundance plot showing distribution of all measured features raw intensities per sample. Potential technical outlier measurements are marked with asterisks. Horizontal orange lines denote fold-difference thresholds used for outlier classification. Vertical dotted lines indicate boundaries between analytical batches. **c.** Retention time vs total chain carbon number for different chain desaturations levels to identify potential misannotations (points as stars, highlighted with arrows), **d.** Isobaric features with highly correlated intensities, due to the higher signal of Cer 18:1;O2/24:0 overlapping on Cer 18:0;O2/24:1. **e.** PCA plot of all raw peak intensities from samples and QCs based on all measured features. **f.** Raw peak intensity of the internal standard for PCs spiked into samples and QCs at same amounts. **g.** Distribution of relative internal standard peak intensities in different sample types.

As mentioned, peak annotation can be challenging in complex ion traces. QUANT provides tools to check for potential annotation errors (**Fig. 4c**). In lipidomics applications, for instance, plotting average RTs against lipid chain length and saturation revealed that some sphingomyelin (SM) isotopologues were misannotated as phosphatidylcholines in our initial analysis (**Fig. 4c, Supplementary Fig. 5a**). In addition, plotting highly correlated feature pairs helps the user identify false duplicate annotation, such as those from isobaric features (**Fig. 4d**).

PCA provides another way to identify potential sample outliers and systematic effects (**Fig. 4e and Supplementary Fig. 6a**). In **Dataset 1**, PCA implemented in QUANT confirmed the outliers identified in the RLA plot, and the corresponding loadings plot revealed specific sources of variability. In this example, Batch QC (BQC) samples, representing different aliquots of a pooled sample extracted independently, were dispersed along the second principal component (PC2, vertical axis). A closer inspection of the loading scores onto PC2 revealed that LPCs are the major contributor to this variability (**Supplementary Fig. 6b**).

QUANT provides flexible functions for plotting feature variables, such as peak areas and RTs over the analysis time, which allows a granular view of analytical performance such as intensity drift and batch effect (**Fig. 4f, Supplementary Fig. 7**). Furthermore, comparing ISTD signal distributions between sample types can highlight potential sample preparation and sample-specific matrix effects, indicated by the considerably higher variability in the ISTD signals of triacylglyceride (TG) across study samples compared to QC (**Fig. 4g).**

#### Quantification and drift/batch effect correction

The QUANT module automatically processes data from raw peak areas to concentrations. This requires detailed metadata, including sample type, the ISTD mapping, standard concentrations, and sample amounts, e.g., volume or protein concentration. With appropriate metadata, QUANT seamlessly performs data normalization using analyte-specific or feature class-wide ISTDs and reports concentration data incorporating external calibration curves or spiked-in ISTD concentrations, as demonstrated in **Datasets 1** and **3**.

While the processing steps aim to reduce analytical variation, they can inadvertently introduce variability and bias as well. For example, class-specific ISTDs or inaccuracies in sample amount determination can introduce such unwanted variation. To assess such potential artifacts, QUANT allows for comparison of the coefficient of variation (%CV) before and after a processing step. In the example, the %CV of TG species in the study samples doubled after normalization, highlighting that substantial noise was introduced (**Fig. 5a**). This was likely due to the previously discussed matrix effects (**Fig. 4g**) that differentially impacted endogenous and ISTD TG features.

**Fig. 5.**
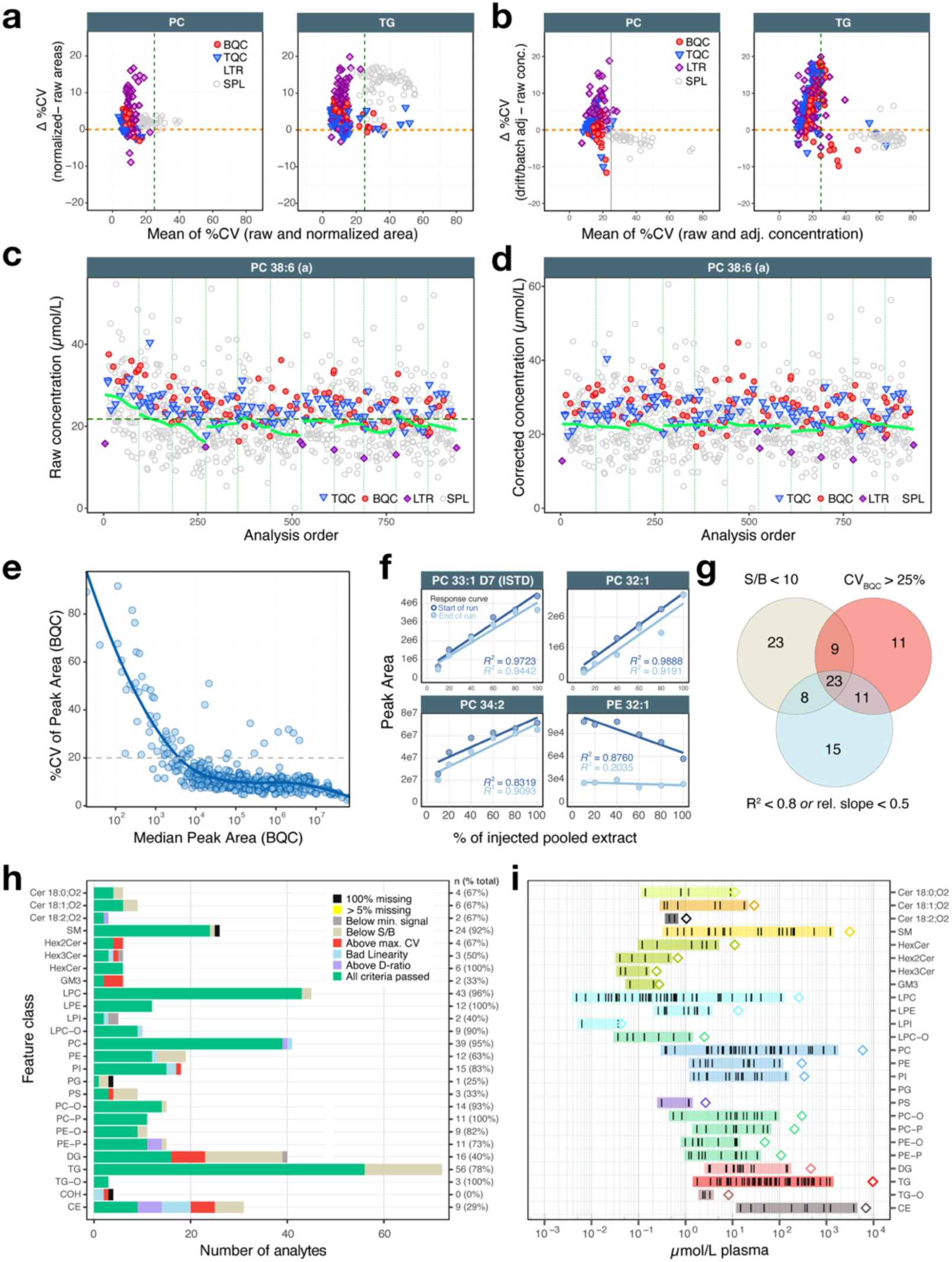
Data postprocessing, QC, and feature filtering. **a.** Effect of internal standard-based normalization on %CVs of features in samples and QCs. Each data points corresponds to one feature of the corresponding class. See **Methods** for more details on individual plots and the QC sample types. **b.** Effect of drift- and batch correction on the %CVs of features in samples and QCs. **c.** Raw peak areas in different sample types of all samples. The dark green lines indicate mean ± 2xSD. **d.** Final, batch and drift corrected concentrations. See panel c. for annotations. **e.** Analytical variability as a function of raw peak area, assessed using BQCs. Each point corresponds to a measured feature. **f.** Response curves of selected features. The x-axis denotes the % of injected extracted plasma volume compared to the volume used with study samples. **g.** Number of features failing one or more of the indicated QC criteria. **h.** Summary of the feature filtering. Features are first evaluated against the lower-level criteria signal-to-blank (S/B) ratios and limit of detection (LOD), followed by the higher-level criteria, coefficient of variation (CV), linear response and D-ratio. **i.** Final concentrations of filtered features grouped by feature class. The diamond symbols indicate total concentration in the corresponding feature class.

Batch and drift effects may still persist after ISTD-based normalization and quantification via calibration, especially in large-scale metabolomics analysis. QUANT provides several drift correction functions, including QC-based locally estimated scatterplot smoothing (LOESS) and cubic spline-based methods commonly used in metabolomics^12, 31^. Batch correction is currently implemented as sample or QC-based median centering. In **Dataset 1**, we applied sample-based Gaussian kernel smoothing and batch centering in QUANT. This reduced the overall %CV of study samples, indicating amelioration of the technical variability from these effects (**Fig. 5d** and **Supplementary Fig. 8b**). The efficacy of these corrections depends on various factors, such as quality of QC samples and adequate sample size. Moreover, these methods are susceptible to issues such as the influence of outliers and the risk of overfitting, which can introduce further bias and noise. This, once again, underscores the importance of supervised corrections^20^.

QUANT enables users to interrogate analysis data using flexible functions for comparing QC parameters, for example by assessing the effect of signal intensity on analytical variability (**Fig. 5e**). Furthermore, QUANT offers assessment of the linearity of feature response relative to the sample amount. This QC step is valuable in metabolomics experiments, considering the wide dynamic range of measured features and that the linear range of an instrument can shift over time (**Fig. 5f**).

Lastly, QUANT provides customizable QC filters to exclude features based on criteria such as high analytical %CV, low signal-to-blank ratio, non-linearity, or high data missingness. It can summarize the reasons for QC failures, the interrelatedness of the criteria, with the number of features passing per analyte class. This QC step can reveal systematic and class-specific analytical issues. For example, the summary of the feature filtering shows that only half of all measured cholesteryl esters species could be measured reliably (**Fig. 5h**). As a final overview plot, the median concentrations of QC-filtered features in the measured samples or reference materials can reveal differences relative to other datasets in which the same materials were analyzed or highlight potential quantification errors when compared with established reference values (**Fig. 5i**).

### Reproducibility and data sharing

By design, MRMhub creates full documentation of peak integration process and facilitates sharing of results. Raw mzML files can also be bundled with the lightweight, standalone INTEGRATOR applications and parameter files, enabling reprocessing and review of the integration step without external software. The tool exports processed data and workflow details can be exported to different formats, including tables, Excel reports, serialized R objects, and community formats such as Bioconductor’s SummarizedExperiment^32^. QUANT also enables reproducible computational notebooks that run and document data processing workflows and can be rendered as HTML, PDF, or Word reports. Together, MRMhub enables sharing of complete metabolomics datasets aligned with FAIR data principles^33^.

### Computational Performance and Benchmarking

To demonstrate the scalability and performance of MRMhub, we applied the tool to a moderately sized data (**Dataset 1**) and a large-scale dataset (**Dataset 3**). We applied MRMhub workflow on an Apple M2 MacBook Pro laptop using multithreading for both datasets (see **Methods**). A single round of peak integration took 8 and 42 seconds for the two datasets respectively, while generating PDFs for the integration results took 0.5 and 4 minutes at most. The core data processing workflow (data import/export and processing) took 0.5 and 1 minute, respectively. Generating all detailed QC plots and reports required additional 3 and 4 minutes. The memory usage for the full workflow did not exceed 1 and 10 GB for the two datasets, respectively.

We remark that these timings reflect purely software run times. For large datasets, it is necessary to perform rigorous review of peak integration results, and subsequently re-optimize an assay-specific, optimized integration parameter set. A few iterations of this process may require one day. Subsequent QC review, outlier investigation, optimization of the normalization and drift/batch correction workflow, and review of feature filtering can typically be accomplished within an additional day, depending on the depth of review and optimization. Addressing major analytical artifacts may require extra time and custom code. Despite the additional requirements, these processes are facilitated by leveraging data structures and interfaces built in MRMhub, which make such modifications efficient and reproducible.

In summary, end-to-end data processing of **Dataset 3** (∼3.7 million features) was completed within two days by a single analyst on a standard laptop computer. The MRMhub workflow is thus substantially faster than conventional approaches that use vendor software for peak integration and spreadsheets or scripts for postprocessing. At the same time, MRMhub ensures a flexible, fully reproducible and documented process with in-depth QC and consistent reporting.

We further benchmarked the performance of MRMhub modules with available open-source tools using **Dataset 1** and **Dataset 3** (see **Supplementary Information**). MRMhub offers the most complete and advanced set of functionalities among available tools to date, providing a robust and scalable framework for end-to-end processing of large-scale metabolomics and lipidomics studies.

## Discussion

We have demonstrated the comprehensive, largely automated functionalities of MRMhub, which provides an ideal software solution for reproducible quantification and data quality control in large-scale targeted MS experiments. Our workflow not only offers full digitization of experimental records and automated data processing steps, but also encourages implementation of rigorous experimental design. MRMhub naturally ensures reproducibility of data processing and facilitates knowledge transfer on an analytical method and datasets from one laboratory to another.

MRMhub has its own limitations nonetheless. While it provides robust, customizable, automated peak integration for complex chromatograms across many samples where other tools fail, it may not resolve or be accurate enough for extreme cases with noise, highly variable convoluted features, or spurious retention time shifts. For such analytes, advanced methods such as Gaussian peak deconvolution or machine learning may be more applicable, but these are currently not available in scalable open-source tools. Ultimately, improving data quality at the source, i.e., at analyte extraction and LC-MS analysis, is more sustainable than attempting to computationally rectify flaws in low-quality data.

At the current stage, the INTEGRATOR module provides an interactive terminal user interface and a dedicated visualizer, while efficient data review is also possible via simple PDF files. The QUANT workflows, on the other hand, is assembled as a computational notebook. Although the workflow lacks a GUI, our current implementation allows for flexible, highly customizable pipelines tailored for different analytical designs and the direct generation of reports that include processing code and corresponding QC visualizations. While the process requires basic familiarity with IDEs such as RStudio, VS Code, or Jupyter, these tools are easily accessible to most lab analysts today. Future work on MRMhub will include developing an integrated GUI.

MRMhub is a comprehensive software solution for targeted MS experiments. The computational framework for automated peak integration and postprocessing can easily be expanded to targeted re-processing of data from other MS acquisition modes such as large-scale untargeted MS experiments for metabolites and proteins. We will continue to develop these extensions of INTEGRATOR module as an open-source solution and provide improved interfaces to connect the data to QUANT module for seamless quantification of analytes and QC.

## Methods

### Datasets

#### Dataset 1: SPERFECT study

The data comes from the SPERFECT study, which stands for “<u>SP</u>hingolipids as s<u>E</u>rial bioma<u>R</u>kers o<u>F</u> disease progression in acut<u>E</u> myo<u>C</u>ardial infarc<u>T</u>ion”^28^. We performed targeted lipidomics analysis of plasma samples from 84 participants at risk of coronary artery disease (CAD). All participants took part in a longitudinal study over up to five follow-up time points with blood collection (378 samples) and some participated in a diurnal rhythm study (299 samples). Including all QC samples, the data consisted of 937 injections for LC-MRM analysis in six analytical batches. The QC samples included several types of samples: blanks with and without lipid extraction, blanks containing ISTD mix, technical quality control (TQC) samples prepared by pooling aliquots of extracted samples, and BQC samples prepared by extracting aliquots of pooled plasma alongside the study samples. TQC and BQC samples were injected at regular intervals throughout the analysis to monitor both analytical and instrument performance.

#### Dataset 2: A failed analysis batch from DYNAMO study

The data comes from a human plasma lipidomic study^9^, where the goal was to identify plasma lipid biomarkers of rapid renal function decline in a cohort of type 2 diabetes patients. The overarching project is called <u>D</u>iabetes Stud<u>Y</u> in <u>N</u>ephropathy <u>A</u>nd other <u>M</u>icrovascular c<u>O</u>mplications (DYNAMO). We carried out the lipidomics analysis method following the method by Huynh *et al* with minor modifications^9, 34^. We extracted data from an analytical batch where the analysis was halted after 266 injections, which was eventually excluded in the analysis reported in the paper. The analytical experiment design is equivalent to that of **Dataset 1**. This analysis batch underwent LC column degradation starting after ∼80 injections, and the gradual RT drift and peak intensity decline persisted until two thirds of the injections were completed.

#### Dataset 3: Main Data from DYNAMO study

This dataset is part of the reported data in the DYNAMO study mentioned in **Dataset 2**. The study consists of four different patient cohorts and our demonstration is based on the largest of the four cohorts. Here we extracted raw MS data for samples collected at National University Hospital, Singapore, including 3,816 plasma samples along with 775 process and quality control samples. For biological and clinical conclusions related to the data, we refer the readers to the reference paper^9^.

#### Conversion of raw files to mzML

INTEGRATOR requires raw data to be in the mzML format^30^. All original Agilent raw files (.d) of Datasets 1 – 3 were batch-converted to mzML on a Windows PC (Intel Xeon E-2186G CPU, 32 GB RAM) using msconvert (ProteoWizard, v3.0.2327727)^35^ with the following settings: output format = mzML; binary encoding precision = 32 bit; write index = true; files converted in parallel = 6. Conversion took 9 minutes and 48 minutes for **Datasets 1** and **3**, respectively. This step can be streamlined by performing automated, sample-wise conversions during the analysis.

### MRMhub – INTEGRATOR module

#### Implementation

INTEGRATOR and its viewer app are implemented as a self-contained, light-weight application for macOS (Apple Silicon) and Windows 11 (x64) that requires no installation. It includes a terminal user interface while its viewer comes with GUI. See **Data Availability** for source code, executables and detailed manual.

Considering the scalability to large-scale projects (e.g., >10,000 samples), INTEGRATOR avoids loading all MS raw data onto the memory simultaneously. Instead, it first surveys peak features across all samples, writes the information to file system as binary files, which are reloaded as needed in subsequent steps. This design significantly reduces the memory footprint and processing times for large-scale datasets, rendering it executable on relatively modest hardware configurations, such as standard laptop.

#### Sample and transition list validation

The mzML file names in the sample list file will be checked against actual files in the data directory provided by the user. Files absent from the directory will be reported in a text file so that the user can check (“missing_files.txt”). The input transition list is used to assign transition names and attributes to chromatograms in each mzML file by matching precursor ion m/z, product ion m/z and RT. Transitions that fail to match any of the chromatograms will be skipped and reported in a text file (“missing_compounds.txt”). Chromatograms in mzML files without any assignment will also be excluded from analysis. The user may proceed with the analysis even if some files or compounds are missing.

#### RT shift correction

The computational algorithm begins with initial RT drift estimation. For each transition, the algorithm considers the ion trace in the user designated reference sample(s) as the reference pattern and uses it to track potential RT drift in all samples (**Supplementary Fig. 1b**). The reference samples are expected to be selected from an early part of the analysis sequence and not dispersed throughout the entire analysis sequence. Next, the chromatograms in all samples are aligned to that of the reference samples in individual MRM windows, yielding estimates of RT shifts. During this process, the user can set a maximum allowance of RT drift (in minutes) in the parameter file (param.txt) using “RT_shift” parameter. The syntax is “RT_shift = [x, y]”, where x and y refer to forward and backward drift allowances, respectively. The algorithm interprets that RT drift is no more than x minutes earlier and no more than y minutes later with respect to the expected RT in the reference samples. In addition, “RT_shift_bound” parameter restricts the RT shift between adjacent sample injections. RT shift correction can also be disabled by setting “RT_shift = [0, 0]”, which may be helpful for dataset with significant variability of features and/or interferences in chromatograms where the cross-correlation approach may result in inaccurate results. In the absence of chromatographic peaks for blank samples, their corresponding RT shifts are determined by interpolation from the shifts observed in the adjacent samples of the analytical sequence.

#### Feature detection

Next, the algorithm performs a first-round search of peak features in all files using the continuous wavelet transform-based pattern matching^36^. For each MRM window, using the resulting survey of features, the algorithm first determines the target number of features per transition as the median number of features detected in the initial survey, and considers it the upper limit for the number of stable and consistent features to be reported during the second-round survey. In the next round, the algorithm maps all peak features across the samples, where it connects each peak in one sample to the corresponding peak in the next sample, ensuring that each and every feature is stable (i.e. can be detected as a feature) and consistent. The key part of this mapping is that peaks are matched *accounting for the RT drift* estimated above. Here, the user can set “RT_tol” parameter to indicate the maximum deviation of RT from the RT value indicated in the method table and the detected. If no feature can be detected within this bound, the integrated area will be reported as zero.

#### Determination of peak integration borders

In the last step, the algorithm goes through the transitions to finalize the integration borders. In the simple case of unimodal ion traces, the user can set “peak_width” parameter quartets to define default start and end integration points (**Supplementary Fig. 1a**). Users can customize the option for individual analytes with a column labelled “peak width” in the transition list table. Further, the user can optionally set “uniform width” parameter in its own column in the transition table so that integration width remains the same between samples for the given transition. The user can also set “left/right integration bound” to fix integration points with respect to the reference samples.

In complex cases with ion traces showing multiple peaks with and without baseline separation, the algorithm searches for valleys between features to cut the integration in the drift-corrected RT space, and reconverts the border positions to the original RT space, and prints out all RT borders to a CSV file for user inspection and manual adjustment for calculation of peak areas (RT_matrix.csv).

#### Reporting Results

INTEGRATOR exports the peak integration results as CSV tables containing peak areas and other peak variables (e.g., RT, integration borders). A peak viewer app can be used to visualize the integration results for inspection. Alternatively, chromatograms with peak integration results can be exported as PDF as an option.

### MRMhub - QUANT Module

#### Design

We implemented MRMhub-QUANT as an R package (http://www.github.com/SLINGhub/mrmhub). It uses a S4-based data object as its central data structure for storing raw, intermediate, and processed data along with processing metadata, named MRMhubExperiment. All core functions operate on this data object, which can be serialized to a file. Its modular design with consistent interfaces supports the extension of the QUANT module or generation of external functions with new processing and QC functions that integrate with existing functionalities. Furthermore, generic functions for tasks such as data import and drift correction are provided, enabling users to develop custom extensions for new data sources or user-defined correction algorithms that are fully integrated into the workflow. Detailed unit tests, together with continuous integration on GitHub, ensure correctness and robustness of data processing and visualization functions at deployment. Documentation and tutorials are available at https://slinghub.github.io/mrmhub.

#### Data processing using computational notebooks

Quarto notebooks were used to store and run the code used for the workflows applied to each dataset (see **Supplementary Codes 1 and 2**). All processed data and figures presented in the main text and as Supplementary Figures were generated with these notebooks and are also available via the corresponding rendered PDF reports. Minor modifications on annotation and scaling of the panels were made in Adobe Illustrator. See **Data Availability** for original outputs.

#### Terminology and visualization themes

In MRMhub, the term *analysis* refers to individual measurements whereas the term *sample* represent physical samples (i.e., extracts). *Features* denote individual peaks, peak groups integrated together, or other distinct reported signals. *Analytes* are defined as measured compounds (individual molecular species) or as species that are mixtures of molecular species that cannot be distinguished in the measurement, such as isomers or isobars.Sample types are denoted as *QC types* to distinguish them from other definitions and are annotated using a specific colour and point-shape scheme used by most plots in the QUANT module. Study samples (SPL) are shown as grey. Pooled batch QCs (BQC), which are aliquots of a pooled sample co-extracted with study samples, are shown in red. Technical (instrument) QCs (TQC), representing injections of a pooled extract, are shown in blue. Processed blanks (PBLK), representing extractions without any matrix (plasma), are shown in green. The long-term reference material (LTR), a pooled plasma used as in-house reference material, are shown in purple. See the MRMhub manual (**Data Availability**) for the *QC types* supported and details of the default visual schemes.

#### Data import and validation

The peak integration results of **Dataset 1**, exported from the INTEGRATOR as a long-format CSV file, was imported into the QUANT data object using the *data_import_mrmhub* function. All data and metadata present in this file, i.e., RT, FWHM (full width at half maximum), integration borders, as well as precursor and product m/z values, collision energy, polarity, injection timestamp, and sample type, are imported. For **Dataset 3**, the peak integration results from Agilent MassHunter Quant, a CSV file, was imported using the *data_import_masshunter* function, including all available metadata (**see Datasets**). Other file formats, such as from Skyline and plain text files are also supported by QUANT but were not used in these example workflows.

Metadata describing samples, features/analytes, spiked-in ISTDs, response curve details, and known concentrations in calibrant and reference samples is collected using an Excel-based metadata template distributed with the QUANT package (**see Datasets**) and then imported into the QUANT data object. Other supported metadata input formats, such as CSV files, individual Excel sheets, and in-memory R data frames, were not used in these examples.

After metadata import, QUANT checks for the consistency of sample and feature identifiers between and within the data and metadata, and for completeness and validity of the imported metadata information. A summary table of detected issues, classified into errors, warnings and notes, is returned. Although the user can then choose to ignore warnings and proceed with the workflow, errors will prevent completion of the data import to ensure integrity of the data in subsequent data processing and visualization steps.

#### Visualizing analysis design and timeline

The analysis sequence of the different QC types is displayed using the *plot*_*runsequence* function. In **Fig. 4a**, the x-axis represents the analysis order, but alternatively, the date and time can be used to show an overview of the analysis timeline, including analysis interruptions. The plot also prints the start and end date time of the analysis, as well as the median time between individual sample injections, will generally corresponds to the run time per sample.

#### Relative Log Abundance plot

The Relative Log Abundance (RLA) plot displays the distribution of log_2_ transformed relative abundances per sample^37, 38^. The relative abundances can either be calculated using median values across all batches (*i*) or batch-wise (*ii*), where *x*_*ij*_ is the abundance of feature *i* in sample *j*, and *b* is the batch in which sample *j* was measured.

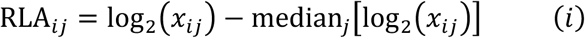

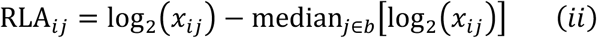

In **Fig. 4b**, only a subset of samples was plotted as illustration, one can display other subsets or the full sequence, see examples in the notebook.

#### Correlating lipid species RTs with chain length and saturation

The median RTs of integrated lipid species peaks in study samples were plotted against the total carbon number of the carbon chains for each lipid class. Robust regression (using the *rla* function from the *MASS* package) were applied to species sharing the same number of double-bonds for visualization or detection of potential outlier that may represent incorrect annotations. Outliers were defined based on either or both criteria: (i) an absolute residual value exceeding 0.15 min, or (ii) a decrease in RT between a species and the following species with higher number of carbons. Outlier data points were flagged with stars, and the corresponding species names are printed to the console. **Fig. 4c** shows the original dataset with annotations errors, which were subsequently corrected (below).

#### Detecting highly correlated feature pairs

In QUANT, feature pairs with a Pearson correlation coefficient > 0.9 can be identified. The module enables comparison of their intensities in individual analyses/samples via scatter plots and allows the user to determine whether the high correlation is not the result of one or few outliers. The different QC types were highlighted using the predefined colour scheme.

#### Updated integration of Dataset 1

After the two previous quality control steps, peak integration of **Dataset 1** was repeated using an updated transition/feature list (**see Datasets**) with corrected expected RTs for the species that were found to be annotated incorrectly. The new updated peak area data was then used in the downstream data processing steps.

#### Plotting time trends of feature variables

Any raw or processed feature variable, such as peak area, RT, ion ratios, or analyte concentration can be plotted across all analyses along the injection sequence as scatter plots via the *plot_runscatter* function. This highly customizable function is used in several sections of this workflow. Features and QC types are subsettable for visualization, e.g., for plotting ISTDs only as shown in **Fig. 4f**. To prevent outliers from skewing the plot and potentially masking trends, this function has an outlier capping option. Control lines (i.e., mean and ±2SD), and overall trends before and after drift or batch correction can also be included (**Fig. 5ab**). The generated plots can either be displayed below the code chunk in notebooks, or exported as a paged PDF for rapid review of large feature numbers and for archival purposes (see **Data Availability**).

#### Matrix effects on internal standards

The peak areas of individually spiked-in internal standards were normalized by their mean across all samples. The resulting values were plotted as box plots, grouped by QC type, with individual data points representing each analysis using the *plot_qc_matrixeffects* function (**Fig. 4g**). This plot assumes that ISTDs were spiked at equal concentrations in all samples.

#### Normalization and quantification by internal standards

For **Dataset 1**, the peak area of each feature in each analysis was normalized to the corresponding peak area of its assigned ISTD, as instructed in the feature metadata. Concentrations were then calculated based on the sample amounts and the amount of the corresponding spiked-in ISTD. The calculation is summarized in (iii) whereby *C*_Analyte_ corresponds to the molar concentration of the analyte in the sample, Area to the peak area of analyte and corresponding ISTD, *V_ISTD_* to the volume of ISTD mix added in µL, Sample_Amount to amount of sample extracted, *C*_ISTD_ to the molar concentration of the ISTD in the ISTD mix.

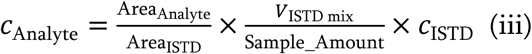

These concentrations are considered as “semi-quantitative concentrations” rather than absolute concentrations, as most ISTDs were non-authentic ISTDs in the lipidomics panels used in this study, with the exception of LPC 18:1[D7], LPE 18:1[D7], and CE 18:1[D7], which were used to normalize corresponding analytes, and because the used ISTDs had only nominal rather than certified absolute concentrations.

#### Response Curves

The response curves, obtained from measuring different amounts of a pooled sample, were plotted using the *plot_responsecurves* function. The corresponding regression results were used later for feature filtering. The sample amount shown on the x-axis corresponds to the percentage of study sample amount injected.

#### Run-order drift correction and batch centering

Correction of run-order drifts was applied using moving average smoothing with Gaussian kernel weights and a kernel size of 20 injections, based on log-transformed study samples raw concentrations only^29^. This correction was applied to all features, separately for each batch, using the *corr_drift_gaussiankernel* function. For comparison, a cubic spline-based smoothing was also performed based on BQC samples using the *corr_drift_cubicspline* function^12, 31^, with the smoothing parameter determined by cross-validation for each feature. After batch-wise drift smoothing, batches were re-centered using batch-wise medians of all study samples within each batch.

#### Feature Filtering

A set of quality control metrics was calculated for each feature and used to filter features. These included: a the signal-to-blank ratio greater than 10 (defined as median of analyte raw intensities in study samples divided by the median in Processed Blanks, PBLK); a median raw intensity study samples greater than 500, a %CV of final concentrations in BQC and TQC samples less than 25%; a D-ratio (the ratio of the standard deviation of final concentrations in study samples to the concentrations in BQCs less than 0.5^20^; an mean R^2^ greater than 0.8 and a mean relative slope greater than 0.5 for the linear regression of the two response curves; and a data missingness in study samples less than 5%. The filtering criteria are applied using the *filter_features_qc* function with parameter defining the individual criteria.

The results of this filtering, i.e., how many species failing one or more of these criteria were summarized per analyte class and overall using the *plot_qc_summary_byclass* and *plot_qc_summary_overall* functions. The latter also returns a Venn diagram showing the number of features failing individual filter categories, i.e., signal-to-noise, analytical CV, and linearity.

#### Data Export

The MRMhubExperiment data object containing all raw and processed data, metadata, QC metrics, and processing status information was exported as serialized R object in the RDS format using the *save_mrmhubexperiment_rds* function. An Excel-based report containing data, metadata, filtered final datasets and processing information was exported as well using the *save_report_xlsx* function. Flat CSV files with feature concentrations as matrix was exported using the *save_dataset_csv* function.

## Supplementary Information

**Supplementary Figures:** Supplementary Figures 1-8

**Supplementary Information**: Supplementary Note 1 with Supplementary Tables 1-2, Supplementary Figures 9-13, and Supplementary References

**Supplementary Code 1**: Code and Results Data Processing for Dataset 1

**Supplementary Code 2**: Code and Results Data Processing for Dataset 3

## Code and Software Availability

Source code for the INTEGRATOR and QUANT modules is available at https://github.com/SLINGhub/MRMhub and has been archived on Zenodo at https://doi.org/10.5281/zenodo.15370293 (ref. ^39^). The repository also contains the source for the MRMhub manuals and tutorials, and the corresponding documentation website is available at https://slinghub.github.io/MRMhub. The Quarto computational notebooks with scripts used for the post-processing of **Datasets 1 and 3** are available at https://github.com/SLINGhub/mrmhub-workflows and have been archived to the same **Zenodo** repository at https://doi.org/10.5281/zenodo.15370293. The corresponding rendered HTML reports are available at https://slinghub.github.io/mrmhub-workflows, and PDFs are provided in **Supplementary Codes 1 and 2.** Executables of INTEGRATOR and its viewer application are available for download from the MRMhub repository.

## Data Availability

MS raw data, analytical metadata, and processed data are available at Zenodo https://doi.org/10.5281/zenodo.1537029339. The MRMhub INTEGRATOR input files and integration result files are also available from the same Zenodo repository, see the corresponding README.txt for data structure and for instructions to reproduce the peak integration step using the INTEGRATOR application. The code used for the post-processing of **Datasets 1 and 3** with the QUANT module is provided as Quarto computational notebooks in **Supplementary Codes 1 and 2**. **Supplementary Code 1** also includes the scripts and original figure files used to generate the figures in this publication. These materials are additionally available as rendered HTML reports at https://slinghub.github.io/mrmhub-workflows. See **Code Availability** for the corresponding underlying code repositories.

## Author contribution

BB, GT and HC developed the algorithms and implemented the software. MRW and MC supervised the SPERFECT study. ET and KV supervised the DYNAMO study. PIB performed all mass spectrometry experiments under the supervision of FT. SHT contributed to sample preparation in the SPERFECT project. BB and ZL performed benchmarking of the existing software solutions. PIB, SJ, JO, PYL, ACG contributed to software testing. HC supervised the MRMhub project. All authors participated in manuscript writing.

## Conflict of interest

All authors declare that the research was conducted without any commercial interest or financial relationships.

## Acknowledgments

The authors thank Jeremy John Selva for providing the Excel metadata template and members of Singapore Lipidomics Incubator, National University of Singapore, for software testing and feedback. BB was supported by Life Sciences Institute, National University of Singapore. This work was supported in part by grants from Singapore Ministry of Education (MOE-000244-00, MOE-000617-00 to HC), National Medical Research Council of Singapore (MOH-000986-00 to MC, HC), and A*STAR (IAF-ICP I1901E0040 to MRW, FT and HC, IAF-ICP I2501E0055 to FT, AMC, MC and HC).

## Notes

https://github.com/SLINGhub/MRMhub

https://doi.org/10.5281/zenodo.15370293

https://slinghub.github.io/mrmhub-workflows

### Competing Interest Statement

The authors have declared no competing interest.

