## Supplementary Figures 1-8 for "MRMhub: one-stop solution for automated processing of large-scale targeted metabolomics data"

### Table of Contents

- Supplementary Figures 1-8

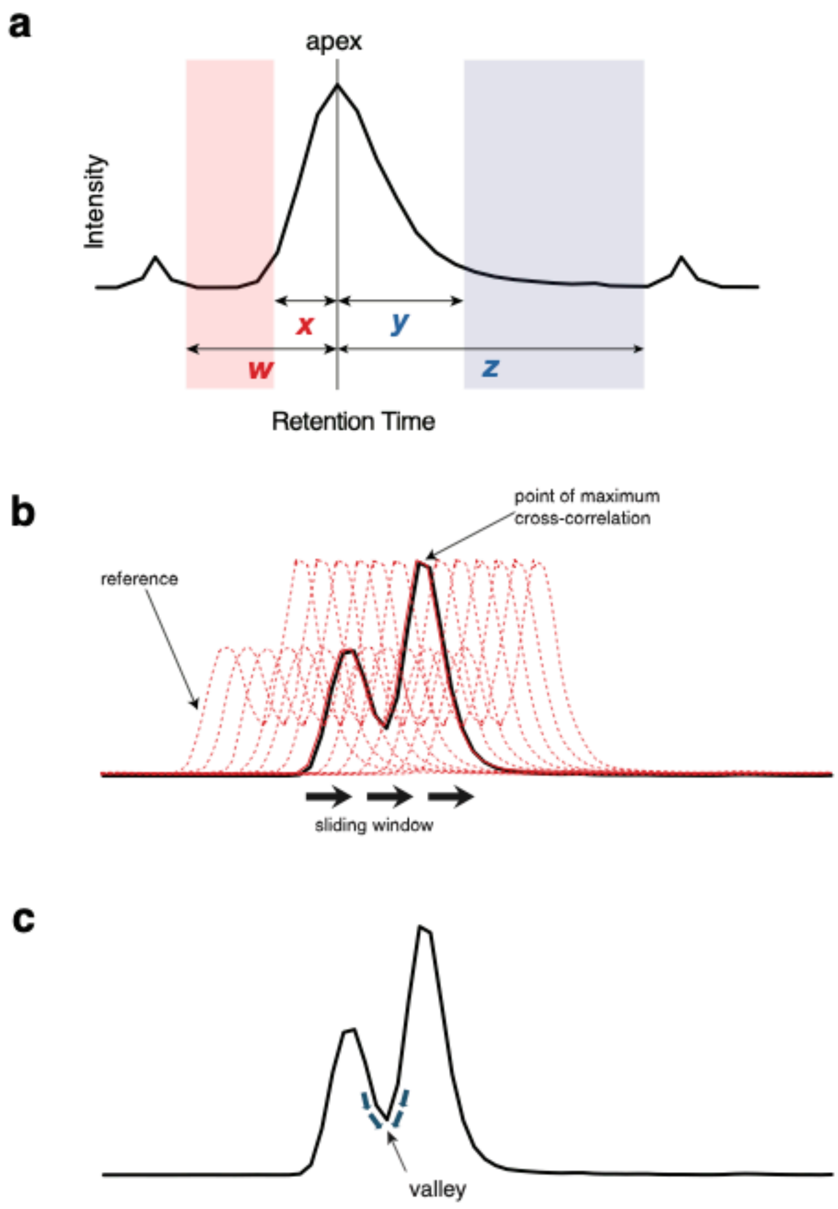


**Supplementary Fig. 1.** Key steps in peak integration by MRMhub-INTEGRATOR **a.** Definition of the peak width variables used with the quartet parameter in the global settings file ("param.txt") and at feature-level in the transition input table. Peak integration will start at the lowest point between w and x and stop at the lowest point between y and z, respectively. **b.** Schematics of the retention time drift estimation. Cross correlation is computed at each potential offset from the reference chromatogram, and RT shift is estimated to be the offset value maximizing the cross correlation. If multiple reference samples were used, it takes the minimum of all estimated RT drifts. **c**. In the case of overlapping features, valleys between modes are identified as integration boundaries.

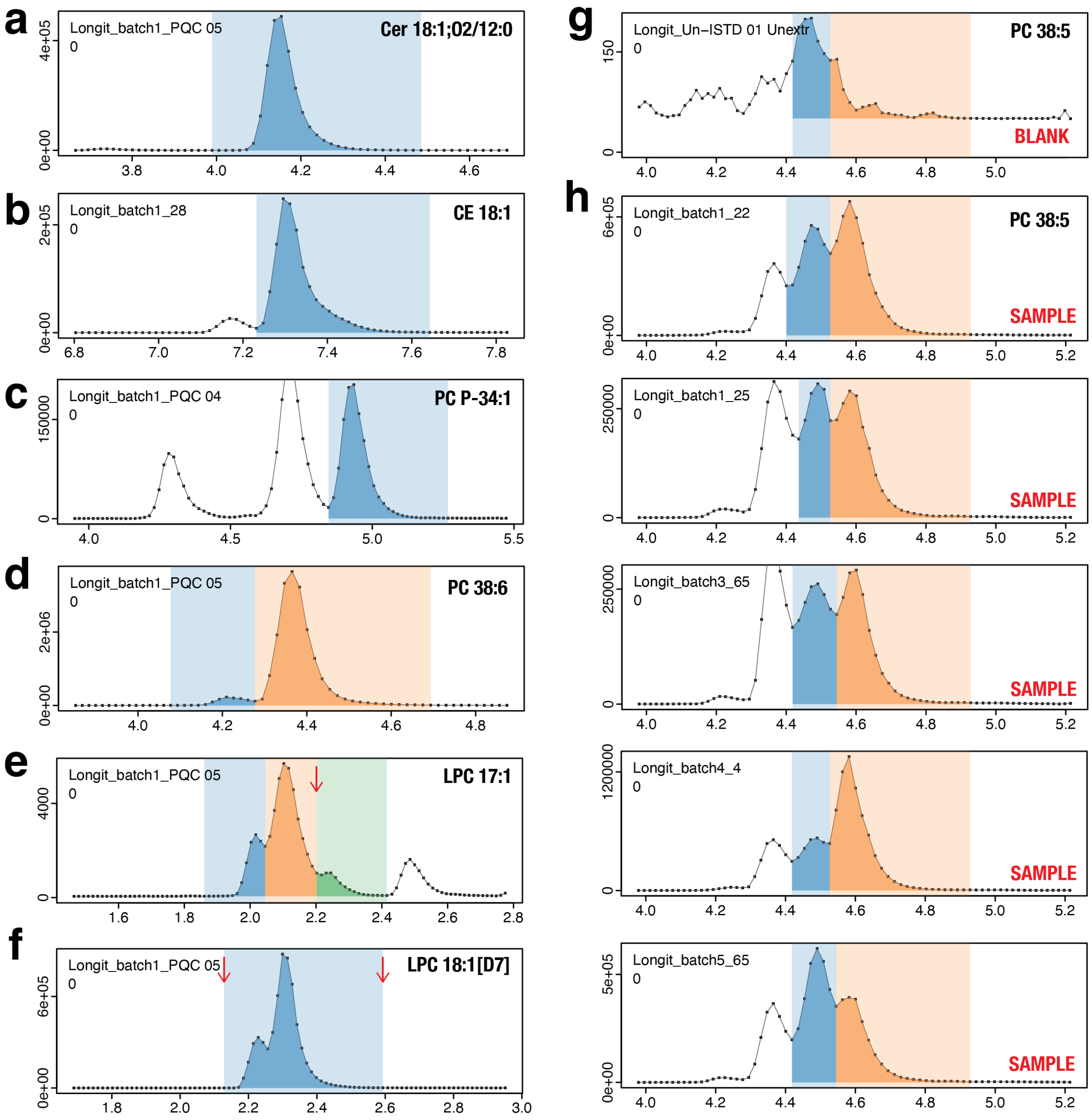


**Supplementary Fig. 2**. Peak integration results for selected features from **Dataset 1**. **a.** Automated integration of a unimodal single peak. **b.** Automated integration of a unimodal peak with increased tailing and adjacent interference arising from the M+2 isotopologue of CE 18:2. **c**. Automated integration of a unimodal peak within a group of isobaric species (PC 33:2 and PC O-34:2). **d.** Automated integration of two isomeric peaks of the lipid species PC 38:6, with differing abundances. **e.** Automated integration of peaks from three partially coeluting isomers. The boundary between the second and third peak, indicated by a red arrow, was set as a fixed value for this transition in the INTEGRATOR feature list input file to ensure consistent integration of this border. **f.** Automatic co-integration of a multimodal peak group representing two coeluting isomers. The left and right borders were set as fixed values (red arrows) to force co-integration of the in vitro formed positional isomers of the internal standard LPC 18:1[D7]. **g-h.** Integration of a blank sample (**g**) using consensus peak boundaries obtained from the integration of samples and QC materials (**h**).

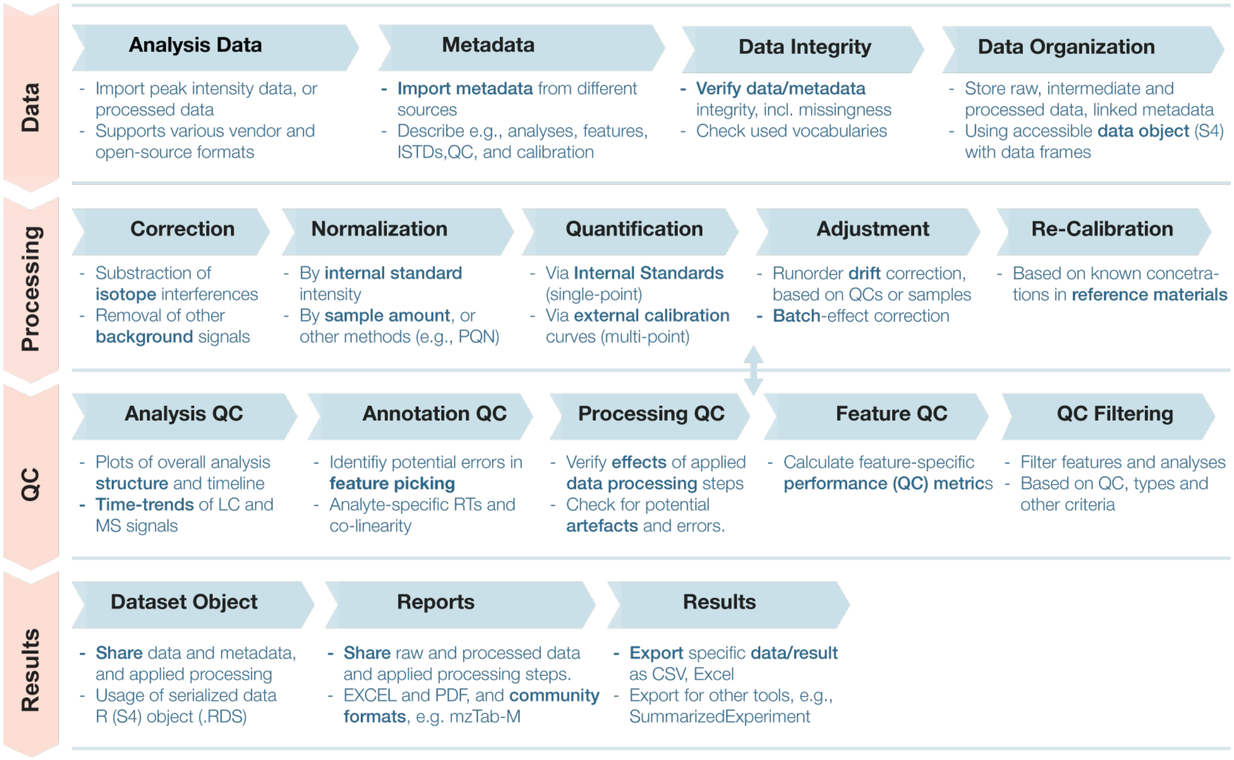


**Supplementary Fig. 3**. Overview of the MRMhub-QUANT functionalities for the data processing and quality control workflow. The workflow is organized into four main sections (red arrows on the left), each comprising specific processing steps (blue arrows). Key functionalities implemented in the QUANT module are shown below each step. Corresponding R functions are provided for each functionality. Detailed descriptions are available in the online manual (see **Data Availability**).

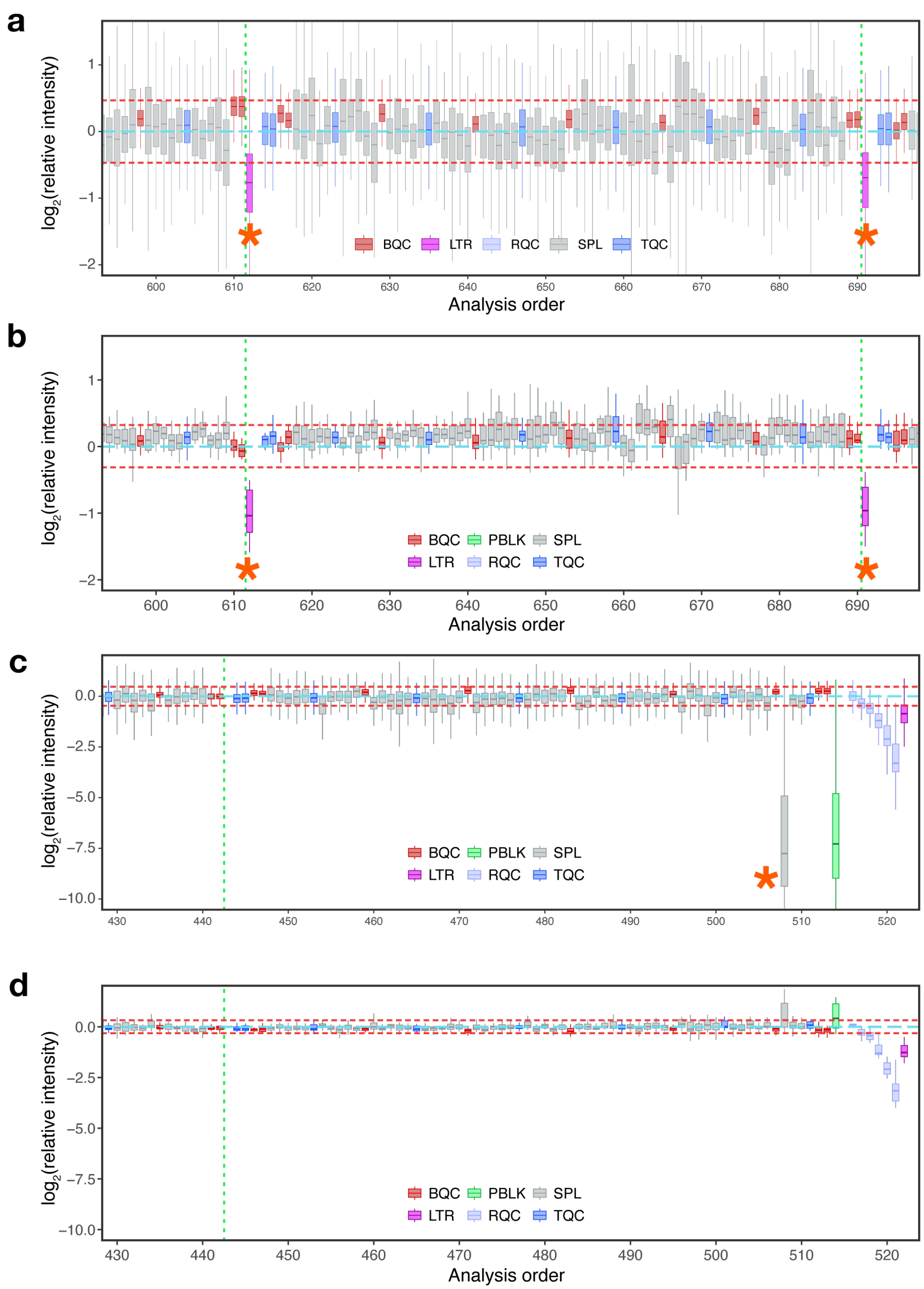


**Supplementary Fig. 4**. Relative abundance (RLA) plots showing the distribution of raw feature intensities per sample. Horizontal orange lines denote fold-difference thresholds used for outlier classification. Vertical dotted lines indicate boundaries between analytical batches. Asterisks (*) denote potential outliers. Boxplot colors and legend refer to QC sample types; see text. **a.** Intensities of non-ISTD features in a selected segment of the analysis sequence, showing that long-term reference (LTR) sample signals may represent potential technical outliers. **b.** As in panel a but displaying only ISTD signals. **c.** Intensities of non-ISTD features at the end of the analysis sequence, showing one outlier study sample in grey between positions 500 and 510. **d.** As in panel c but displaying only ISTD signals. The outlier observed in panel c aligns with other study samples in panel d, indicating ISTD concentrations comparable to those of the other study samples.

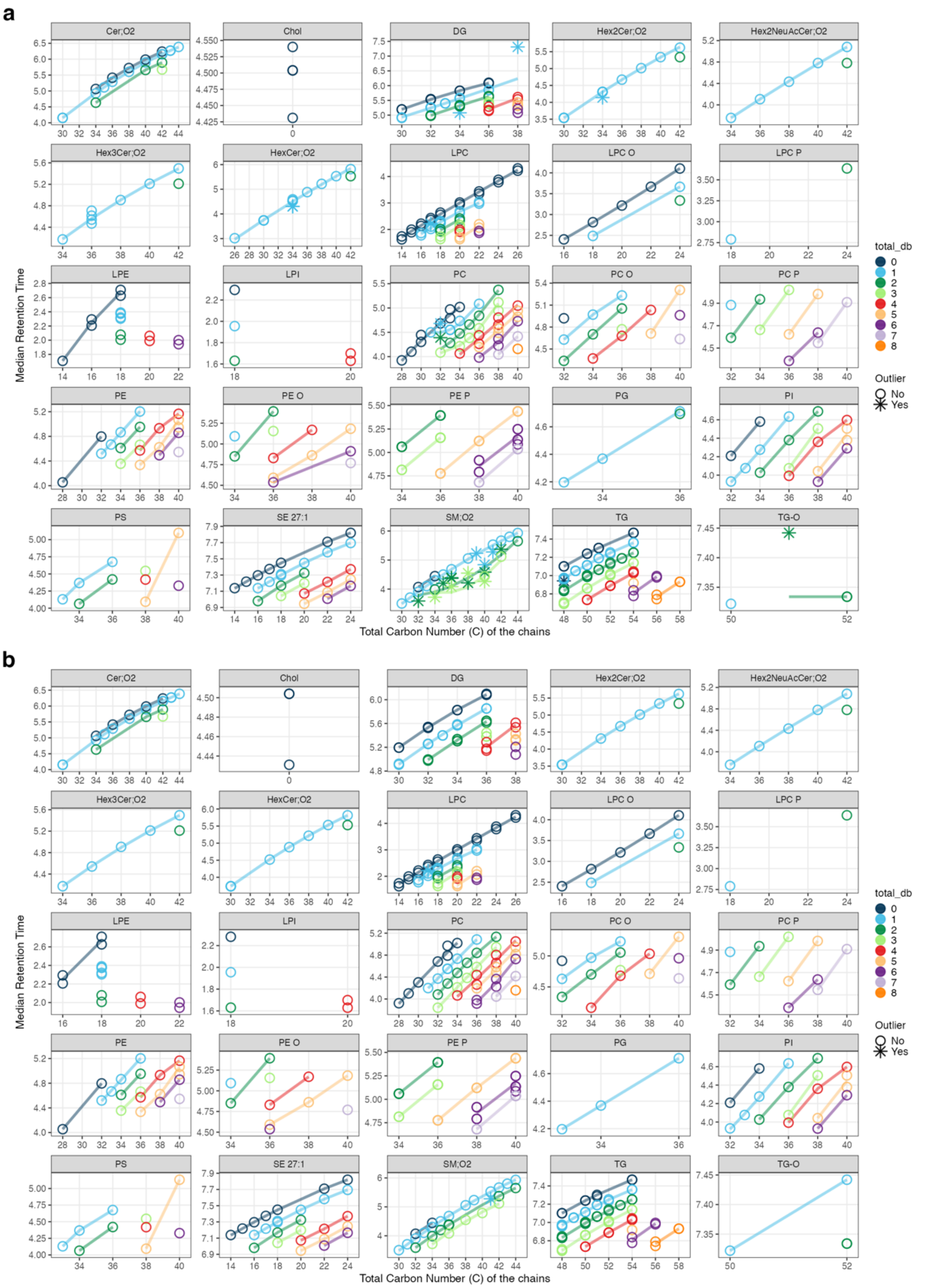


**Supplementary Fig. 5**. Retention time vs total chain carbon number for different chain desaturation levels. **a.** The initial dataset with potential peak annotation errors, particularly in phosphatidylcholines (PC) and sphingomyelins (SM), indicated by stars instead of circle points. **b**. Revised dataset with corrected peak picking.

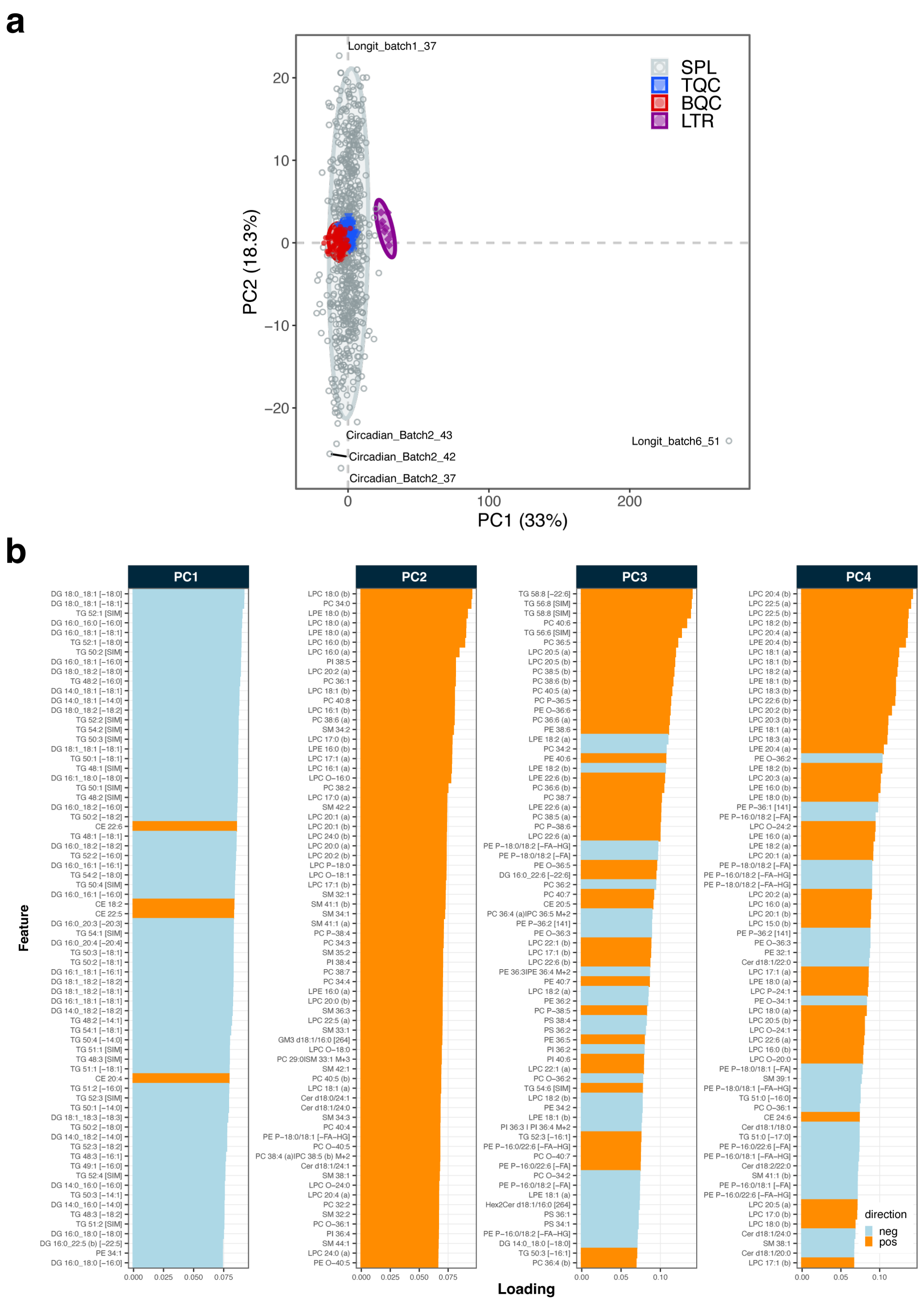


**Supplementary Fig. 6.** PCA of raw peak intensities from samples and QCs based on all measured features. **a.** PCA plot of the initial dataset, with the identified study sample representing a technical outlier in the bottom right corner, labelled with the corresponding sample identifier. **b.** PCA loading plot of the revised dataset. Bars represent absolute loading scores of features for the PC dimensions 1 – 4. Blue indicates negative, orange positive loading.

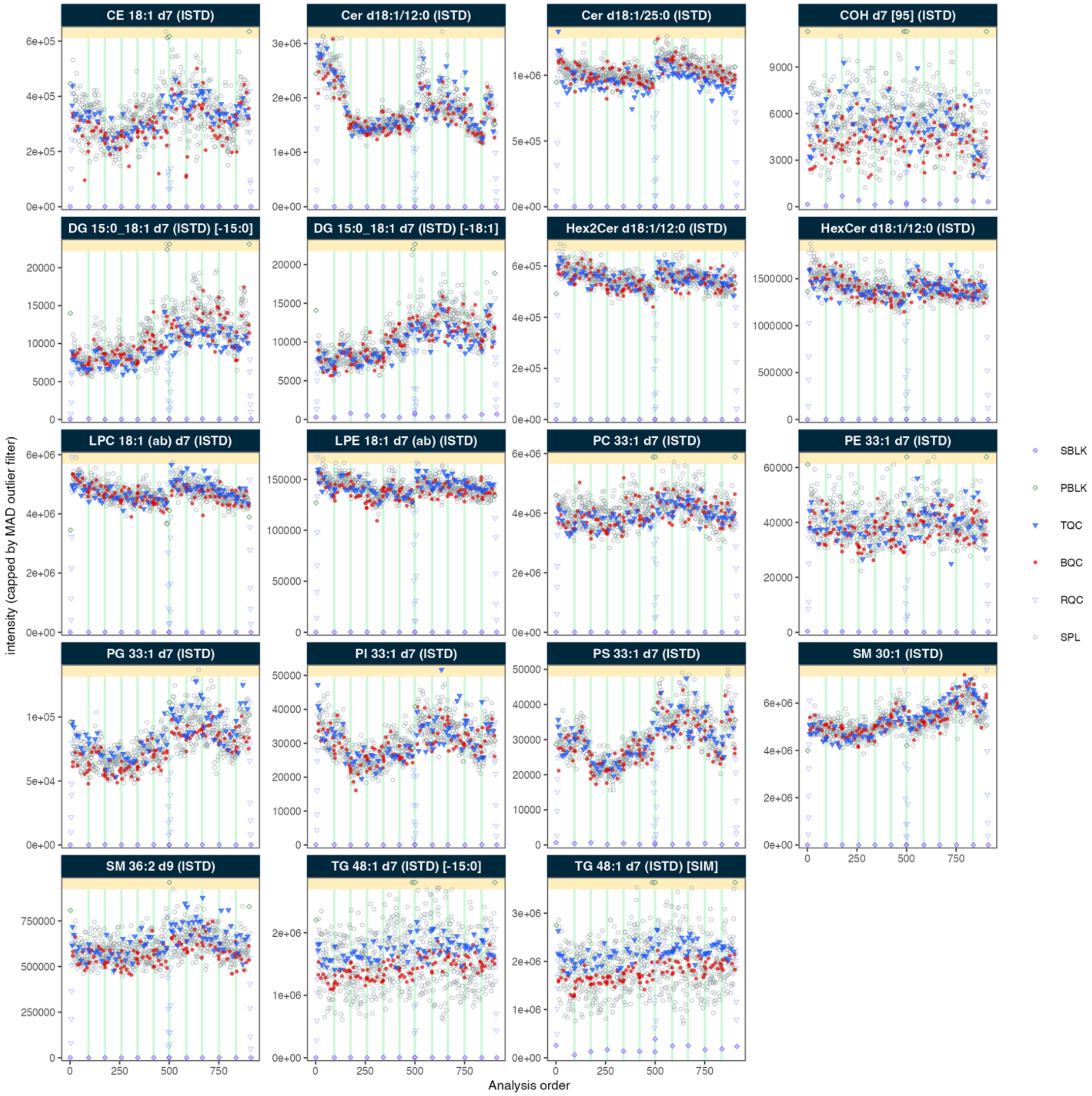


**Supplementary Fig. 7.** Scatter plots of raw peak intensities of all spiked-in internal standards. Batches are separated by vertical green lines. Extreme values exceeding 4x MAD are capped and displayed in the top yellow shaded area of the plots.

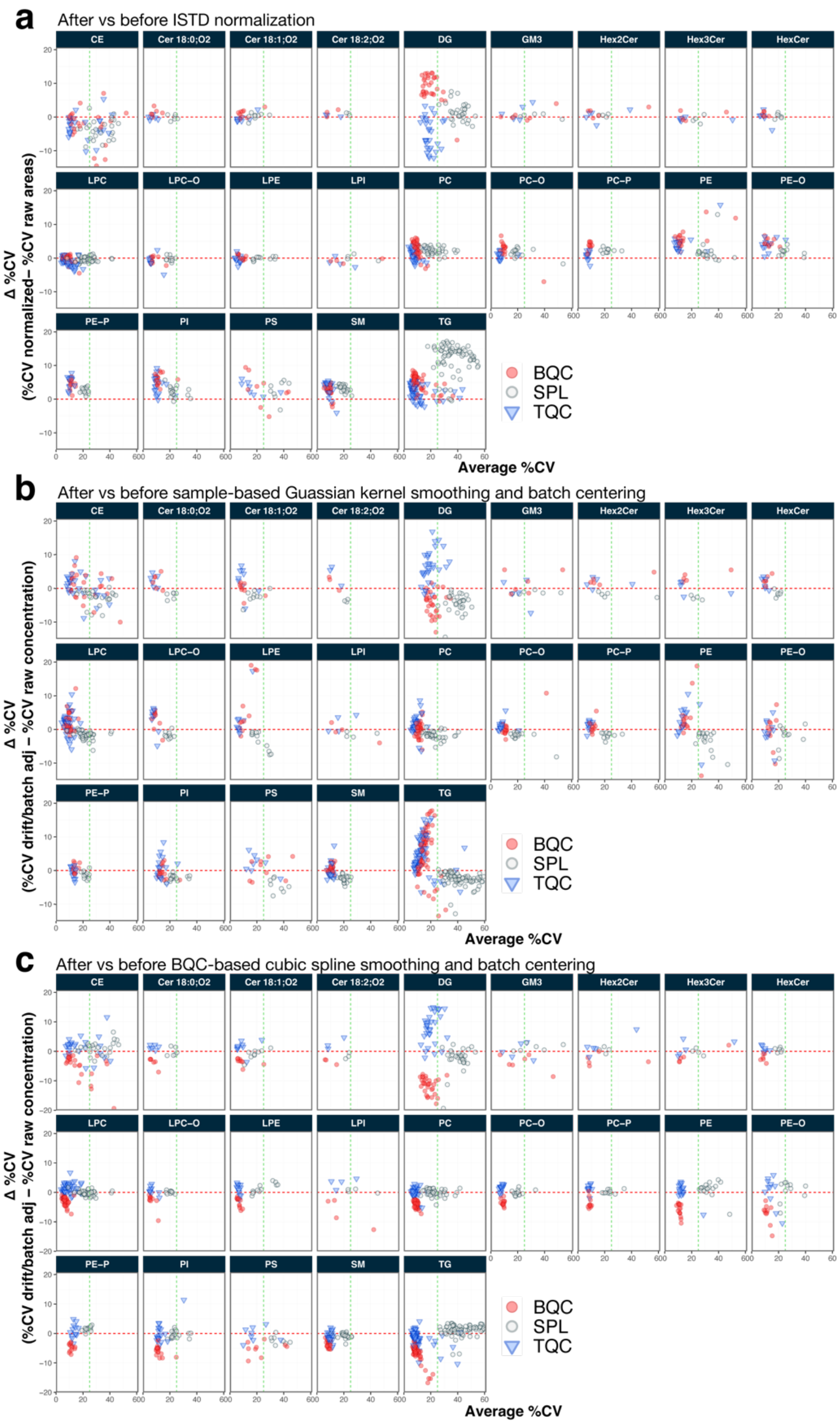


**Supplementary Fig. 8.** Effect of normalization and drift- and batch-correction on CVs of samples and QCs of features from different classes. **a.** Comparison of CVs between raw and ISTD-normalized peak intensities. Features from each class were normalized with the corresponding class-specific ISTD, **b**. Comparison of CVs before and after sample-based Gaussian kernel smoothing followed by median centering. **c.** Comparison of CVs before and after BQC-based cubic spline smoothing with feature-specific parameter optimization, followed by median centering.
