## Supplementary Information for "MRMhub: one-stop solution for automated processing of large-scale targeted metabolomics data"

### Table of Contents

- Supplementary Note
- Supplementary Table 1
- Supplementary Table 2
- Supplementary Figures 9-13
- Supplementary References

### Supplementary Note

We compared the two MRMhub modules to the following available tools with regard to published and reported features: MRMPROBS[1], Skyline [2, 3], MRMQuant [4], and automRm[5] (**Supplementary Tables 1–2**). INTEGRATOR does not have a graphical user interface (GUI) and lacks the ability to use multiple transitions for determining feature boundaries, but it appears to provide the most comprehensive set of features. However, in targeted lipidomics and metabolomics applications, single transitions per feature are often used because including multiple transitions increases cycle times and limits the number of measurable concurrent transitions. INTEGRATOR was therefore designed to operate on single transitions and instead borrow information across samples to detect peaks and determine peak boundaries.

The QUANT module also provides an extensive feature set that exceeds what is currently available in other tools and programming libraries. It is the sole R package to date offering extensive post-processing functionalities for different types of targeted analyses. It also provides a distinct comprehensive test for quality control with flexible plotting options. While it also lacks a GUI, QUANT can serve as a well-tested, encapsulated library for building future applications that may include GUIs.

We additionally conducted tests with five of the tools: *MRMhub-INTEGRATOR*, *MRMPROBS* (v3.73), *SmartPeak (v1.27.0)*, *Skyline* (v24.1), and *MRMQuant* (v2.5). *automRm* was not included, due to its reliance on including at least one qualifier transition and, ideally, of corresponding$^{13}$C-labelled internal standards for each analyte for optimal performance[5]. Dataset 1 could be imported and processed with all tools, and in all cases, processing required less than 2 min for INTEGRATOR, and between 20 to 60 min for the other tools on a PC (Xeon W-1270P CPU, 64 GB RAM, Win 11 x64). The user interface remained mostly responsive in MRMPROBS, SmartPeak, and Skyline, whereas in MRMQuant navigation through the dataset was slow, with considerable waiting times when selecting data subsets.

Next, we evaluated these tools for their performance in automatically integrating chromatograms with multiple overlapping peaks, which are common in lipidomics and metabolomics. We used PC 38:5 as the test case, corresponding to the analyte examples shown in **Fig. 11** and **Extended Fig. 10**. Except MRMhub-INTEGRATOR, none of the other four tools consistently and accurately integrated a single peak from a chromatogram containing overlapping peaks (**Supplementary Fig. 9 – 5)**. This includes those tools with functionality intended to handle convoluted peaks, i.e., SmartPeak and MRMQuant (see **Supplementary Table 1**). An alternative approach is to explicitly set feature-specific peak boundaries that are applied uniformly across all samples. Among the tools tested, only MRMQuant provides this option. However, the boundaries must be defined in the GUI using a reference sample for each dataset and are not corrected for retention time shifts[4].This limitation reduces its applicability for large-scale datasets. Besides, MRMPROBS and Skyline offer sample-specific adjustments of boundaries for individual features via their GUIs, which, however, is impractical and error-prone with large-scale datasets. While all tools provide interactive visualization of integration results as individual or overlaid chromatograms, only MRMPROBS allowed for efficient review of all peak integrations for features across all samples via the GUI. None of the tools offered an option to export integrations as reports, such as PDF.

### Supplementary Table 1: Comparison of the functionalities and properties of INTEGRATOR with freely available tools.

Classifications were based on the corresponding publications and available online resources (documentation and source code) for each tool evaluated. + indicates that the feature is available, - indicates that the feature is absent, and o indicates that the information was not available, or no conclusion could be drawn.

| | MRMhub INTEGRATOR | MRMPROBS | SmartPeaks / OpenMS | Skyline | MRMQuant | automRm |
|---|---|---|---|---|---|---|
| **Design** | | | | | | |
| Type | app | app | app | app | app | R pkg/ ShinyUI |
| Graphical User Interface (GUI) | - | + | + | + | + | + |
| Cross-Platform | + | - | + | - | + | + |
| Multithreaded | + | - | + | + | - | - |
| Portable | + | - | - | - | - | - |
| Programmatic access/API | + | - | + | + | - | + |
| **Data Import** | | | | | | |
| Direct import of native (vendor) raw data | - | - | - | + | - | - |
| Processing of only subsets of transitions or samples | + | + | + | + | - | + |
| **Peak Picking and Integration** | | | | | | |
| Retention time correction | + | + | + | - | - | + |
| RT window definable | + | + | + | + | + | + |
| Use of qualifiers in peak detection | - | + | + | + | - | + [1] |
| Integration parameter adjustable per feature | + | - | - | - | + | - |
| Option of fixed boundaries per feature / adjusted for RT shifts | +/+ | - | - | - | + [2]/- | - |
| Option to use of consensus boundaries | + | - | - | - | + | + [3] |
| Option to set min/max peak widths per feature / left and right of apex separately definable | +/+ | +[4, 5]/- | + [5] | - | +[4, 5]/- | + [5] |
| Multiple features per transition | + | + | + | + | + | + [1] |
| Integration of overlapping features (e.g. deconvolution) | + [6] | - | + [7] | - | +[8] | + [1] |
| Detailed Feature QC metrics | + [9] | - | + | + [9] | - | + |
| Manual modification of boundaries per sample | + [10] | + | - | + | + | + |
| Efficient review of peak integration via plot grids (traces of one feature across all samples) | + | + | - | - | - | - |
| **Tests conducted with Dataset 1** [11] | | | | | | |
| Import and initial processing successful | + | + | + | + | + | o [1] |
| Application responsive | + | + | + | + | - | o [1] |
| Robust automated integration of overlapping peaks | + | - | - | - | - | o [1] |
| Option to efficiently review integration in all samples | + | + | - | - | - | o [1] |

[1] Requires at least one qualifier transition and ideally a corresponding 13C-labelled internal standard for optional performance, see Notes 1
[2] Via the GUI
[3] As result of the machine learning
[4] Only minimum
[5] Only globally
[6] Drop-to-baseline
[7] Exponentially Modified Gaussian (EMG) model fitting
[8] Gaussian model fitting
[9] Integration borders, estimated FWHM, height
[10] Via editing a peak table
[11] See below, Supplementary Information Note 1

### Supplementary Table 2: Comparison of the functionalities and properties of QUANT with freely available tools.

Classifications were based on the corresponding publications and available online resources (documentation and source code) for each tool evaluated. + indicates that the feature is available, - indicates that the feature is absent, and o indicates that the information was not available, or no conclusion could be drawn. See text for references concerning the tested tools.

| | MRMhub QUANT | MRMPROBS | SmartPeak | Skyline | MRMQuant | tidyMS | nPYc - Toolbox | lipidr | QC:MXP |
|---|---|---|---|---|---|---|---|---|---|
| **Data Processing** | | | | | | | | | |
| Type | R pkg | App | App | App | App | Python pkg | Python pkg | R pkg | App |
| Graphical User Interface (GUI) | - | + | + | + | + | - | - | - | + |
| Support computational notebooks | + | - | - | - | - | + | + | + | - |
| Open source | + | - | + | + | + | + | + | + | + |
| Cross-Platform | + | - | + | + | + | - | + | + | + |
| Multithreaded | + | - | + | + | | + | + | - | + |
| Programmatic access/API | + | - | + | + | - | + | + | + | - |
| Does not require additional coding | + | + | + | + | + | - | - | + | + |
| Includes detailed unit tests | + | o | + | + | - | + | + | + | - |
| **Data Formats** | | | | | | | | | |
| Direct import MRMhub INTEGRATOR results | + | - | - | - | - | - | - | - | - |
| Direct import of results from external tools | +[1] | - | - | - | - | - | +[2] | +[3] | + |
| Flexible import of wide and long tabular data files with option to map columns | + | - | - | - | - | - | | + | - |
| Import of more than one feature variable (e.g. area and RT) | + | - | - | - | - | - | + | - | - |
| **Data Processing** | | | | | | | | | |
| Normalization with Internal Standard (ISTD) / Feature-specific | +/- | +/+ | +/+ | +/+ | +/ | -/- | -/- | +/- | -/- |
| Quantitation via spiked-in ISTD | + | - | - | + | - | + | + | | + |
| Quantitation via external calibration | + | + | + | - | + | - | - | - | - |
| QC-based drift correction / batch-wise / multiple models | +/+/+ | +/-/+ | -/-/- | -/-/- | +/+/- | +/-/- | - | | +/+ |
| Sample-based drift correction available | + | - | - | - | - | - | - | - | + |
| Stand-alone batch correction / selectable ref. sample type | +/+ | -/- | -/- | -/- | +/- | -/- | +/+ | -/- | +/- |
| Technical outlier detection and exclusion | +/+ | - | - | ? | - | - | +/- | - | - |
| **Feature Filtering** | | | | | | | | | |
| Support for various QC types / User-defined QC types | +/+ | +/- | +/- | | | | | | |
| Feature filter QC variability/D-ratio/signal-to-blank /missingness | +/+/+/+ | - | - | - | - | +/+/+/+ | + | +/-/-/- | +/+/+/- |
| Response (dilution) curve correlation filtering / multiple curves / slope | +/+/+ | - | - | - | - | +/-/- | +/-/- | - | - |
| Filter applicable to different feature variables / QC type [4] | +/+ | | - | | | | - | - | - |
| **QC Metrics + Plots** | | | | | | | | | |
| Run-order scatter plots / outlier-capping | +/+ | - | - | +/- | - | - | +/- | - | +/- |
| PCA / loading plot | +/- | +/- | - | - | - | | +/- | | |
| (Relative) log abundance box plots | + | - | - | - | - | - | - | + | - |
| Heatmaps | - | - | + | - | + | - | - | - | - |
| Feature Annotation QC plots[6] | +[6] | - | - | - | - | - | - | - | - |
| Response (dilution) curve plots | + | - | - | - | - | - | - | - | - |
| Feature filterring summary plots | + | - | - | - | - | - | - | - | - |
| Output feature-specific plots as facetted, multi-page plots to PDF/console | +/+ | - | - | - | - | - | - | - | - |
| **Documentation + Data Sharing** | | | | | | | | | |

| | | | | | | | | | |
|---|---|---|---|---|---|---|---|---|---|
| Data processing workflow documentation | + | | + | | | + | + | | |
| Export report | + | | + | | | | | | |
| Customizable formats (long/wide, variables, sample types) | + | - | - | - | + | - | - | - | - |
| Export to community formats | | - | + | + | | | +[7] | | |

[1] Currently formats from Skyline and Agilent MassHunter Quantitative are supported
[2] Skyline and Agilent MassHunter Quantitative are currently supported
[3] Progenesis QI, XCMS, XCMS online, MZmine, MS-DIAL, Biocrates are currently supported
[4] Skyline and Metabolomics Workbench (mwTAB) are currently supported
[5] Whether filter can be used for e.g. raw and normalized peak areas, or concentrations, and based on different QC or Blank types
[6] For lipid species, based on retention time and total carbon number, saturation, or effective carbon number (ECN)
[7] in ISA-TAB format

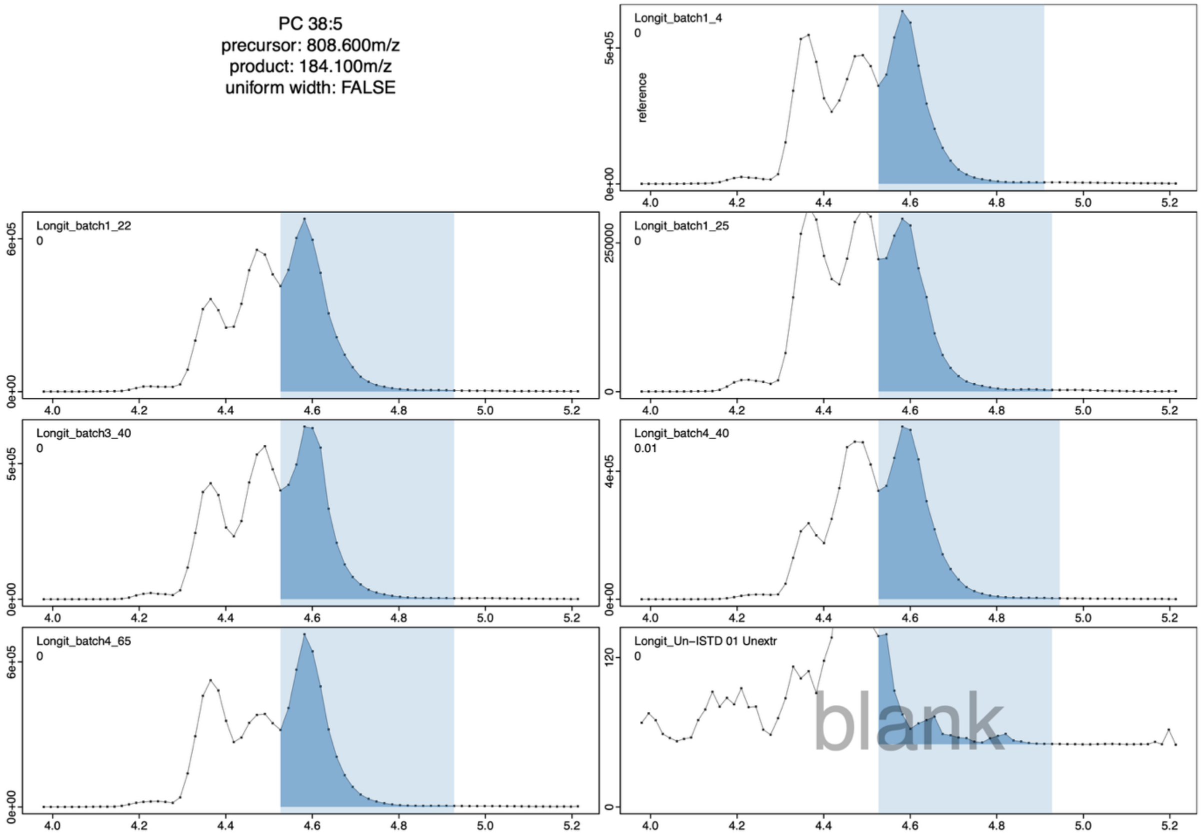


**Supplementary Fig. 9: MRMhub-INTEGRATOR integration results of PC 38:5 from selected samples of Dataset 1.** The feature targeted by the feature input list is the rightmost peak (isomer *b*). Each panel corresponds the chromatogram of one sample. The integrated peaks are depicted by dark blue shaded areas, the left and right integration boundaries, respectively, by light blue shaded areas.

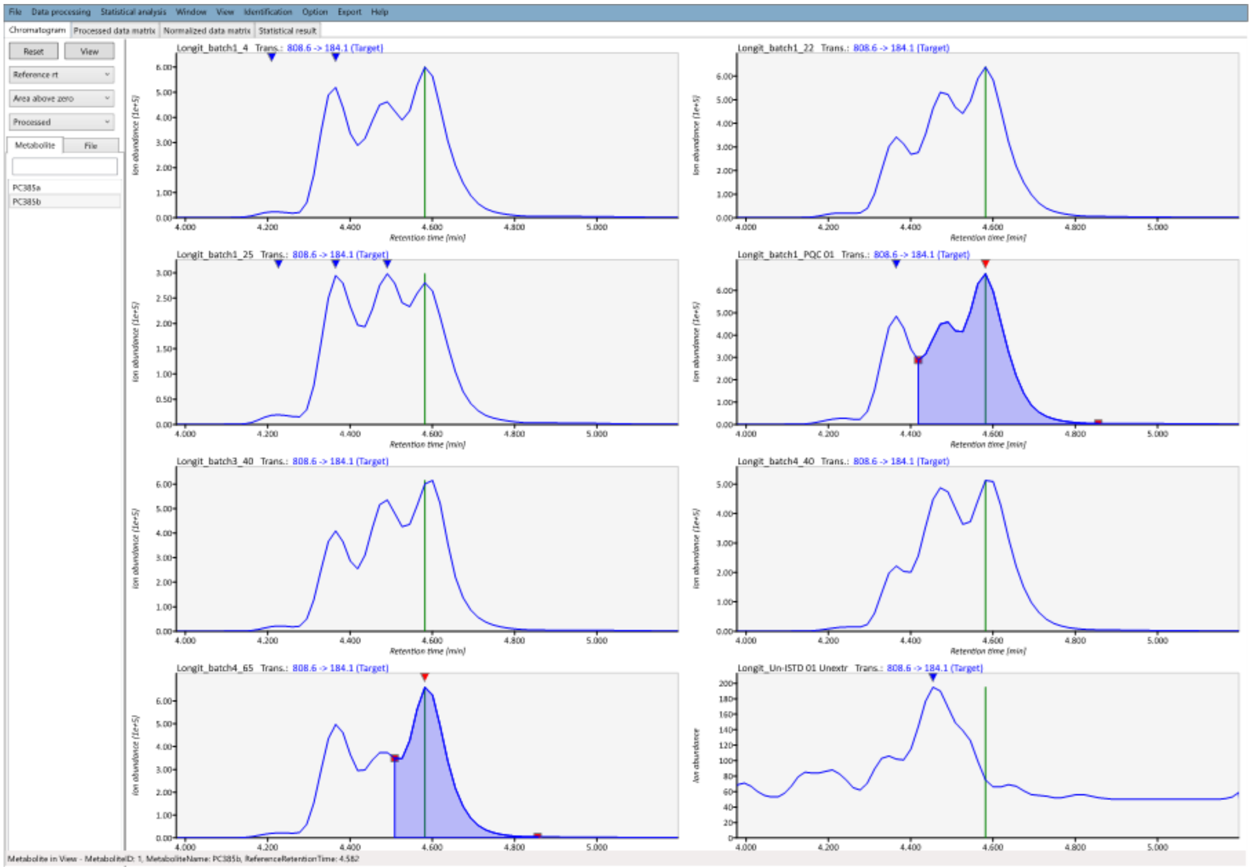


**Supplementary Fig. 10: MRMPROBS integration results of PC 38:5 from selected samples of Dataset 1.** The peak targeted by the feature input list is the rightmost peak (isomer *b*), indicated by the green vertical line. Each panel corresponds the chromatogram of one sample, using the same samples as shown in **Supplementary Fig. 9**, using the same samples. The peak detected within the target RT range is indicated by a red triangular arrow, while other peaks are marked by blue triangular arrows. The integrated range is depicted by the blue shaded area.

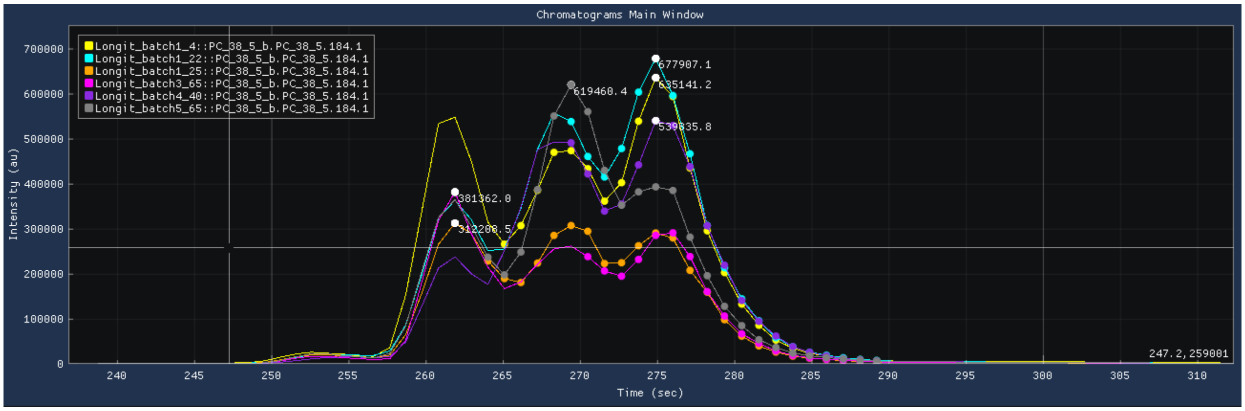


**Supplementary Fig. 11: SmartPeak integration results of PC 38:5 from selected samples of Dataset 1.** The peak targeted by the feature input list is the right most peak (isomer *b*). Each trace corresponds to the chromatogram of one sample. The determined peak apex for each trace is indicated with the number corresponding to the signal height. The integrated range is depicted by the shown datapoints for each trace, whereby segments of the middle and left peaks were included in most samples. Regions beyond these points are not part of the detected peaks.

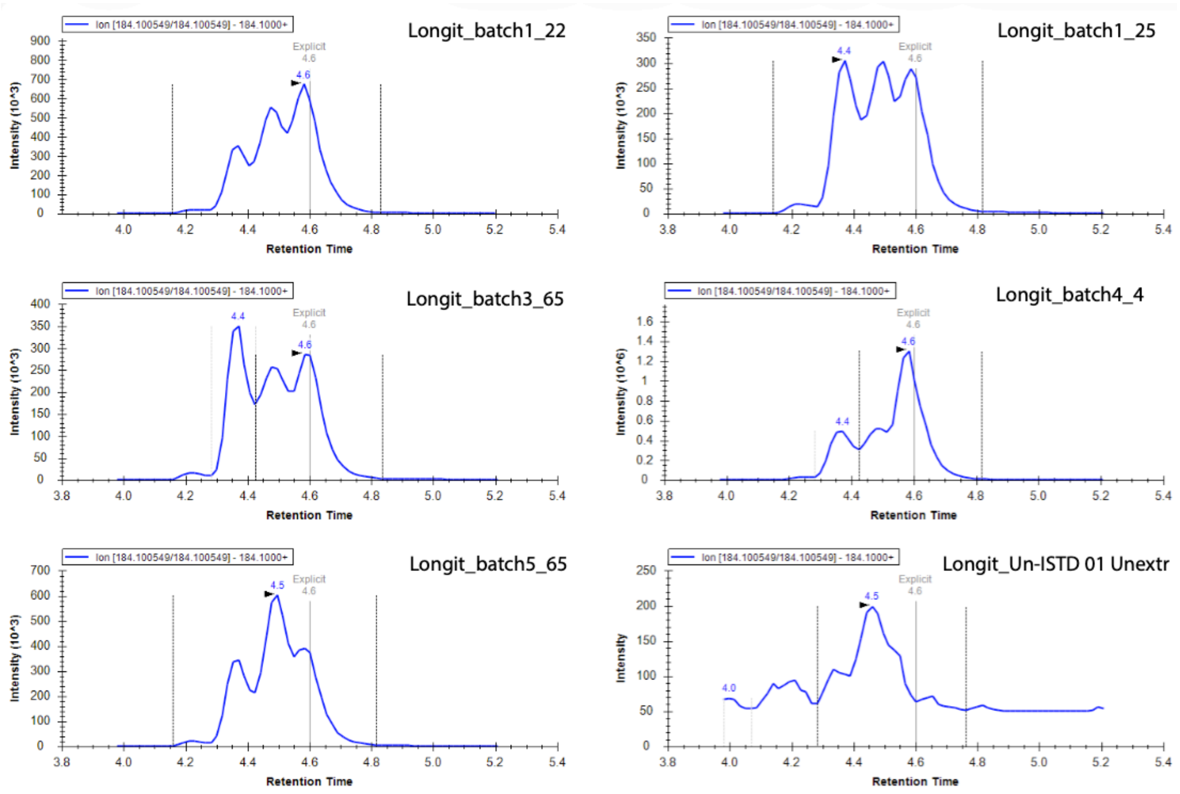


**Supplementary Fig. 12: Skyline integration results for PC 38:5 from selected samples of Dataset 1.** The peak targeted by the feature input list is the rightmost peak, indicated by the vertical line at 4.6 min. Each panel corresponds the chromatogram of one sample, using the same samples as shown in **Supplementary Fig 9.** The determined peak apex for each trace is indicated with a triangular arrow. The detected peak is depicted by the left and right most vertical lines.

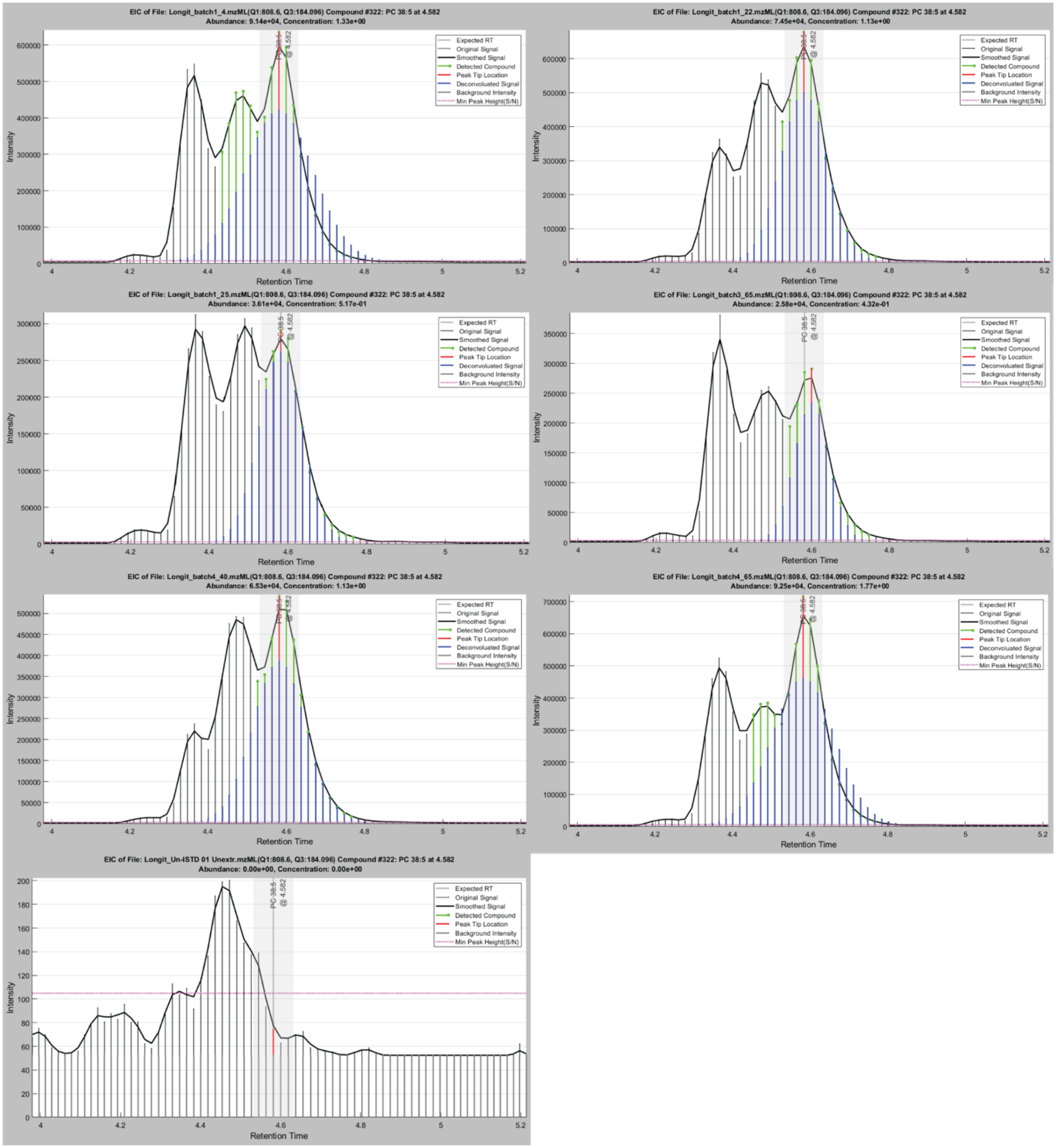


**Supplementary Fig. 13: MRMQuant integration results for PC 38:5 from selected samples of Dataset 1.** The peak targeted by the feature input list is the rightmost peak (isomer *b*), indicated by the vertical line at 4.582 min. The shaded grey band show the RT window used for peak detection. Each panel corresponds the chromatogram of one sample, using the same samples as shown in **Supplementary Fig 9**. The determined peak apex for each trace is indicated with a vertical red line. The detected peak is shown in green, and the corresponding modelled Gaussian peak shape is represented by blue vertical lines.
