## Supplementary Code 1 for "MRMhub: one-stop solution for automated processing of large-scale targeted metabolomics data"

### MRMhub Data Processing Workflow for Dataset 1

**Supplementary Code 1**

Bo Burla Guo Shou Teo Hyungwon Choi

2025-11-24

**References 60**

### 1 Overview Dataset

This example workflow uses a targeted lipidomics dataset measuring 503 features across 937 samples (Tan *et al.* (Tan et al. 2022)) and demonstrates the complete MRMhub data processing pipeline, including peak integration, quantitation, quality control, and reporting. This document covers many elements of the MRMhub workflow for demonstration purposes, some of which may not be applicable or necessary when processing other datasets.

> **!** Important
>
> All datasets used and generated within this workflow have been deposited in the **Zenodo** repository https://zenodo.org/records/15370293. Within this repository, the raw mass

spectrometry files and the MRMhub-INTEGRATOR input files required for peak integration are available under the record "**Dataset 1**". The data and metadata files used and produced by the code in this Quarto notebook can be found under the "**mrmhub-workflows**" record. Please note that due to size limitations, the corresponding GitHub repository does not include any of these datasets.

### 2 Raw Data Processing: Peak Picking and Integration

This section describes key aspects of raw data processing workflow, i.e. peak picking and peak integration, used for this dataset. For more details on the usage of the INTEGRATOR application, see the INTEGRATOR Manual.

#### 2.1 Conversion of vendor raw files to mzML

INTEGRATOR requires the raw files to be in the mzML format. The original raw data files (Agilent .d) were converted to mzML using msconvert from the ProteoWizard software (https://proteowizard.sourceforge.io) with the following settings: output format = mzML; binary encoding precision = 32 bit; write index = true. See the MRMhub Manual for more details.

#### 2.2 Preparing the INTEGRATOR input files

INTEGRATOR requires three different input files. Templates and examples are included in the downloaded MRMhub-INTEGRATOR release archive. The final input files used for this dataset are available from the Zenodo repository (see above). The following subsections describe each required input file and the key settings used for this example.

##### 2.2.1 Global Settings File

The parameter mz_tol, defining the maximum tolerance in m/z values for identifying transitions, was set to 0.06. The RT_tol parameter, defining the window around the expected RT (see Feature/Transition table below) in which features are searched, was set to 0.1 min. This relatively narrow RT window was chosen to reduce selection of incorrect peaks in complex chromatograms, as observed in this dataset. In this dataset peaks exhibited some tailing, thus the *peak_width* was set to [0.15, 0.5, 0.1, 0.35], with the right-side ranges set to longer than the left-side ranges. For specific features with broader or convoluted peaks, feature-specific peak widths were applied, overwriting the global settings (see Feature Table below). The maximum allowed RT shift correction was set to (-0.2, 0.2), and the maximum allowed sample-to-sample RT shift was 0.1 min. The full set of settings in param.txt is shown in the figure below.

```
### location of .mzML files
mzML_files =  'mzML/*.mzML'

### table indicating file name, batch, sample type and reference sample
batch_info = 'run_order_20251009.csv'

### Table with transition information:
transition_list =  'transition_list_20251009_FINAL.csv'

# Number of threads to use for parallel processing
num_threads = 14

### m/z tolerence, maximum difference between m/z in transition list and in mzML file
mz_tol = 0.06

### RT tolerence, maximum difference indicated RT and detected RT
RT_tol =  0.1

### Peak width limits: [w, x, y, z], the left integration bound will be between w and x mir
peak_width =  [0.17, 0.05, 0.1, 0.35]

### if "RT_shift = [x, y]", the estimation will assume that RT shift is no more than x minu
RT_shift = [-0.2, 0.2]

### this parameter restricts RT shift between adjacent samples
RT_shift_bound = 0.1

### if false, empty fields in "RT_matrix.csv" will be errors, if true, empty fields will be
#fill_RT = true
```

Figure 1: **Global settings file (param.txt).** Screenshot of the param.txt file used for this dataset, showing all configured parameters and their values.

##### 2.2.2 Sample Table

In the sample table (run_order_20251009.csv) the samples to be processed were listed by their .mzML file names. The first two BQC (batch quality control) samples were selected as retention time reference anchors. The sample type for each sample was defined in column C (sample_type) using the MRMhub QC type nomenclature. This is optional and INTEGRATOR distinguishes only between blanks (identified by the presence of the text `BLK`), and non-blanks.

| | A | B | C | D |
|---|---|---|---|---|
| 1 | file name | batch | sample_type | reference |
| 14 | Longit_batch1_PQC 01.mzML | L1 | BQC | x |
| 15 | Longit_batch1_PQC 02.mzML | L1 | BQC | x |
| 16 | Longit_batch1_1.mzML | L1 | SPL | |
| 17 | Longit_batch1_2.mzML | L1 | SPL | |
| 18 | Longit_batch1_3.mzML | L1 | SPL | |

Figure 2: **Sample Table.** Screenshot of a portion of the CSV file used for this dataset (run_order_20251009.csv). Two samples are selected as RT reference samples.

##### 2.2.3 Feature/Transition Table

The feature/transition table (transition_list_20251009.csv) defined the transitions to be extracted from the .mzML files by precursor and product m/z values. Each feature (peak or peak group) was specified by its expected retention time (RT, in minutes). For some transitions, multiple features were defined. The corresponding internal standard feature for each entry was included in the table, however, these internal standards were not used by INTEGRATOR but were exported in the integration results.

The feature and transition table, provided as a CSV file, specifies the transitions to be extracted from the mzML files and the features (peaks) to be integrated from each transition (chromatogram).

| 1 | Compound Name | Transition Name | ISTD | Precursor Ion | Product Ion | RT | uniform_width | left integratio | right integratio |
|---|---|---|---|---|---|---|---|---|---|
| 235 | PC 36:3 | PC 36:3 | PC 33:1 d7 (IS | 784.6 | 184.1 | 4.567 | n | | |
| 236 | PC 36:4 (a) \| PC 36 | PC 36:4 | PC 33:1 d7 (IS | 782.6 | 184.1 | 4.25 | n | | |
| 237 | PC 36:4 (b) | PC 36:4 | PC 33:1 d7 (IS | 782.6 | 184.1 | 4.433 | n | | |
| 238 | PC 36:5 | PC 36:5 | PC 33:1 d7 (IS | 780.6 | 184.1 | 4.2 | n | | |
| 239 | PC 36:6 (a) | PC 36:6 | PC 33:1 d7 (IS | 778.5 | 184.1 | 3.88 | n | 3.79 | 3.917 |
| 240 | PC 36:6 (b) | PC 36:6 | PC 33:1 d7 (IS | 778.5 | 184.1 | 4 | n | 3.917 | 4.21 |
| 241 | PC 38:2 | PC 38:2 | PC 33:1 d7 (IS | 814.6 | 184.1 | 5.15 | n | | |

Figure 3: **Feature/Transition Table.** Screenshot of a portion of the CSV file (transition_list_20251009_Final.csv) used for this dataset. Fixed integration borders were set for two adjacent features to ensure correct integration.

For this dataset, fixed borders were defined for some features that were poorly separated from adjacent peaks or that had noisy chromatograms to ensure correct and consistent integration, for example PC 36:6. In some other cases, isomers were intentionally co-integrated, such as the LPC and LPE internal standards (see Figure 3).The feature and transition table used in this workflow was optimized for an earlier version of INTEGRATOR. Some feature-specific integration settings included in that table may not be necessary with newer versions of the software. The final version of the feature/transition table (transition_list_20251009_FINAL.csv) is the result from several rounds of parameter optimization with INTEGRATOR and from peak picking QC using the QUANT module, see below.

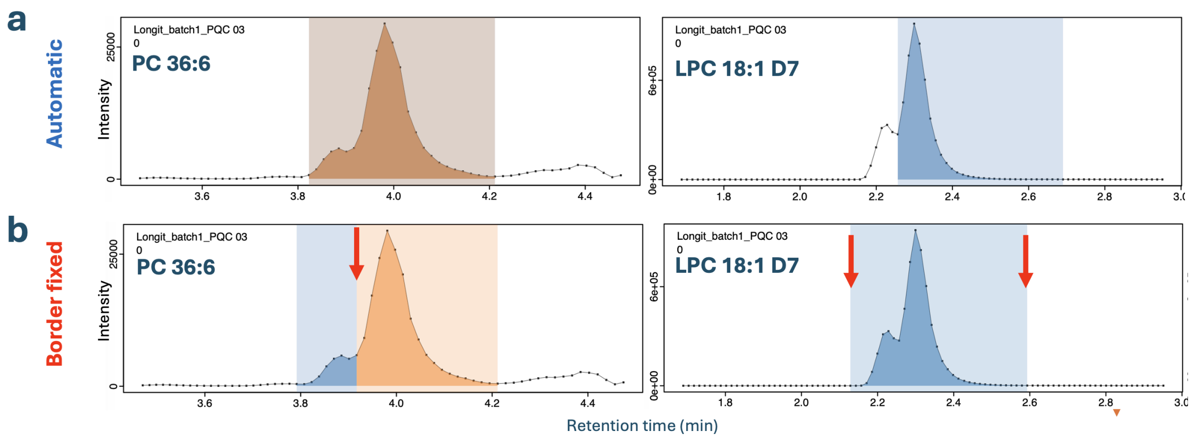

Figure 4: **Examples of manually set peak boundaries for MRMhub-INTEGRATOR**. (a) Automatic integration resulting in co-integration of both overlapping peaks (PC 36:6) or integration of only one peak (LPC 18:1 D7). (b) The same features integrated with fixed boundaries for one and two peak boundaries, respectively. Red arrows indicate feature-specific fixed integration border(s).

#### 2.3 Running MRMhub-INTEGRATOR and Review of Results

The most recent release OF INTEGRATOR was obtained from https://github.com/SLINGhub/MRMhub/releases. After preparing all input files, the mrmhub application is started. Steps 1 to 4 then run consecutively without further input to perform peak detection, peak picking, integration, and export of data and PDF results. The integration results are reported in the generated file 'long.csv', a long-format table that includes peak areas, actual retention time, peak width, and other metadata. This data file will be used for subsequent data postprocessing (see the following section). A wide-format table with peak areas is available as 'quant_raw.csv'. PDFs of the integrated transitions are available in the by_* folders. See the INTEGRATOR Manual for more details.

The peak integration results were inspected for all features across samples using the generated PDFs (see above). In this dataset, RT shifts across the entire run of 937 samples were less than 0.02 min

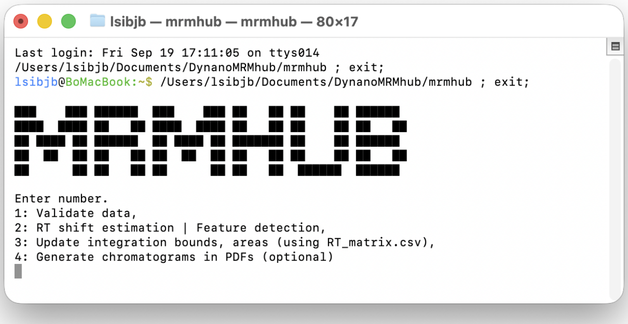

Figure 5: **MRMhub-INTEGRATOR application.** The sequential steps of the MRMhub-INTEGRATOR workflow are initiated by selecting the corresponding number on the keyboard. The intermediate and final datasets generated by this application are stored in the same directory.

### 3 Data Postprocessing and QC

The postprocessing (Quantification, Quality Control and Reporting) of the data obtained from the peak integration above are conducted using MRMhun-QUANT, which is implemnted as the {mrmhub} R package. A Quarto computational notebook is used the run, document and report the data post processing.

#### 3.1 Setting up a Quarto Notebook and Installation of {mrmhub}

A Quarto notebook was created in RStudio the {mrmhub} R package was installed by running following code in the R console, which also installs two other R packages used in this workflow.

```
if (!require("remotes")) install.packages("remotes")
remotes::install_github("SLINGhub/MRMhub", subdir = "quant")
install.packages("here") # Used for parallel processing
```

The Quarto notebook code generated in this example workflow is available from https://github.com/SLINGhub/MRMhub-workflows (file Dataset3.qmd). In the subsequent section, the data postprocessing is encoded and documented step-by-step into the notebook as individual code chunks using {mrmhub} functions. Certain code chunks include additional R code to showcase specific examples for illustrative purposes. In real-world applications, this supplementary may not be necessary.

#### 3.2 Postprocessing workflow

##### 3.2.1 Load mrmhub and other required packages

To improve performance multi-threading was used for some of the calculations and plottings, for which the R package {mirai} needs to be installed and loaded.

```
library(mrmhub)
library(mirai)  # Used for parallel processing
```

##### 3.2.2 Import the MRMhub-INTEGRATOR results (initial version of the dataset)

We import the results of the initial peak integration iteration obntained with the INTEGRATOR workflow, described in the previous section. The corresponding original result file 'long.csv' has been renamed to 'sPerfect_LongCirc_MRMkit_Initial.csv'. The results contain peak areas (`intensity`), retention time (`rt`), and peak full width at half maximum (`fwhm`). Available metadata, i.e., sample type, acquisition time stamp, precursor and product m/z values are imported as well (`import_metadata = TRUE`). The data is stored in a `MRMhubExperiment` object, which will be used in subsequent steps as data object (container).

```
data_path <- "./data/dataset-1/data/sPerfect_LongCirc_MRMkit_Initial.csv"
mexp_initial <- MRMhubExperiment()
mexp_initial <- import_data_mrmhub(mexp_initial, data_path, import_metadata = TRUE)
#> v Imported 937 analyses with 503 features
#> i `feature_area` selected as default feature intensity. Modify with `set_intensity_var()`
#> v Analysis metadata associated with 937 analyses.
#> v Feature metadata associated with 503 features.
```

##### 3.2.3 Analytical design and timeline

An overview of the analysis structure, detailing the types and sequence of QC samples analyzed is provided by the plot below. It also shows information on the start and end dates, total duration, and median run time for each sample.

```
fig4a <- plot_runsequence(
  mexp_initial,
  show_batches = TRUE,
  qc_types = c("SPL", "BQC", "TQC", "PBLK", "UBLK", "RQC", "SBLK", "LTR"),
  batch_zebra_stripe = TRUE,
  base_font_size = 6,
  batch_fill_color = "#fffbdb",
```

```
  segment_linewidth = 0.5,
  show_timestamp = FALSE) +
theme(plot.title = element_text(size = 5))
fig4a
```

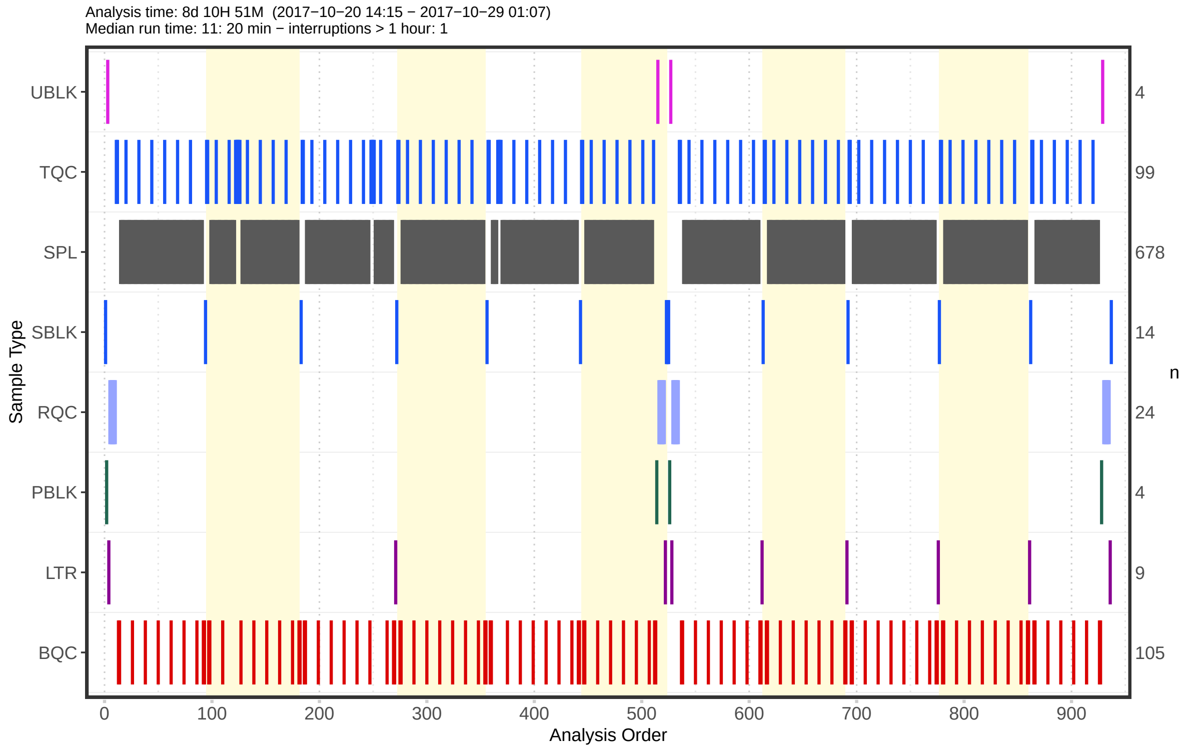

##### 3.2.4 Overall trends and check for possible outliers

To examine overall technical trends and issues affecting most analytes (features), the Relative Log Abundance (RLA) plot is useful (De Livera et al., 2015 (Livera et al. 2015)). In an RLA plot, each feature is normalised to the across-sample or within-batch median and the resulting values are shown as boxplots for each sample. This visualization helps identify technical problems such as pipetting errors, sample loss, or changes in injection volume or instrument sensitivity.

First, an overview of all analyses (samples) is plotted.

```
plot_rla_boxplot(
  data = mexp_initial,
  rla_type_batch = c("within"),
```

```
  variable = "intensity",
  qc_types = c("BQC", "SPL", "LTR", "TQC", "RQC", "PBLK"),
  filter_data = FALSE,
  #analysis_range = get_batch_boundaries(myexp, batch_indices = c(5,6)),
  y_lim = c(-3,3),
  show_timestamp = FALSE,
  outlier_method = "iqr",
  outlier_k = 1.5,
  base_font_size = 6,
  outlier_exclude = FALSE,
  x_gridlines = FALSE,
  batch_zebra_stripe = FALSE,
  linewidth = 0.1, show_plot = TRUE)
#> i Found 26 outliers in the 919 shown analyses
```

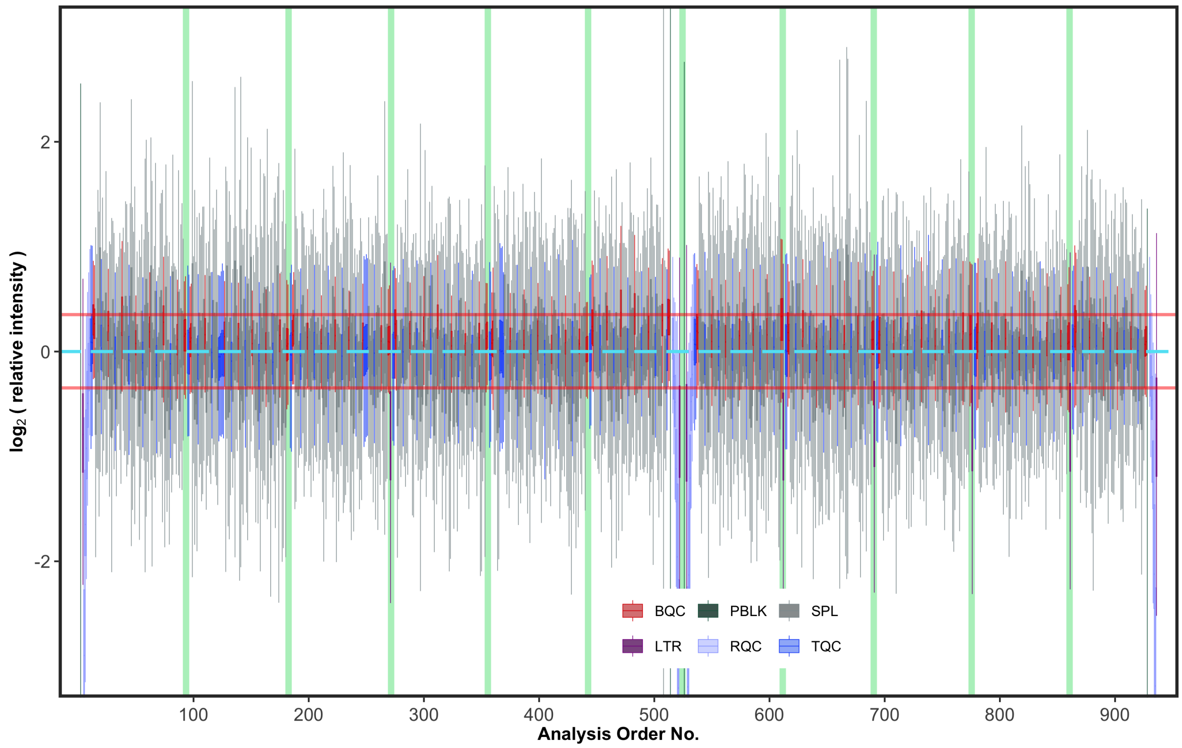

A zoom-in reveals that the engogenous (non-ISTD) feratures of one QC sample type, the LTRs (long-term reference) have in average an approximately 1/2 lower intensity compared to other samples.

```r
# An outlier sample, discussed later, will be excluded here
# for demontration purposes.

mexp_initial_temp <- mexp_initial
mexp_initial_temp  <- exclude_analyses(
  mexp_initial_temp,
  analyses = "Longit_batch6_51",
  clear_existing = TRUE)
#> i 1 analyses were excluded for downstream processing. Please reprocess data.
res <- plot_rla_boxplot(
  data = mexp_initial_temp,
  rla_type_batch = c("across"),
  variable = "intensity",
  qc_types = c("BQC", "SPL", "LTR", "TQC", "RQC"),
  filter_data = FALSE,
  exclude_feature_filter = "ISTD",
  plot_range =c(595,696),
  y_lim = c(-2,1.5),
  include_istd = FALSE,
  include_qualifier = FALSE,
  min_feature_intensity = 500,
  show_timestamp = FALSE,
  outlier_method = "iqr",
  outlier_k = 1.5,
  #outlier_k = c(-1, 1),
  outlier_exclude = FALSE,
  x_gridlines = FALSE,
  batch_zebra_stripe = FALSE,
  base_font_size = 6,
  show_plot = FALSE,
  linewidth = 0.1
)
#> i Found 19 outliers in the 914 shown analyses
fig4b <- res$plot
fig4b
```

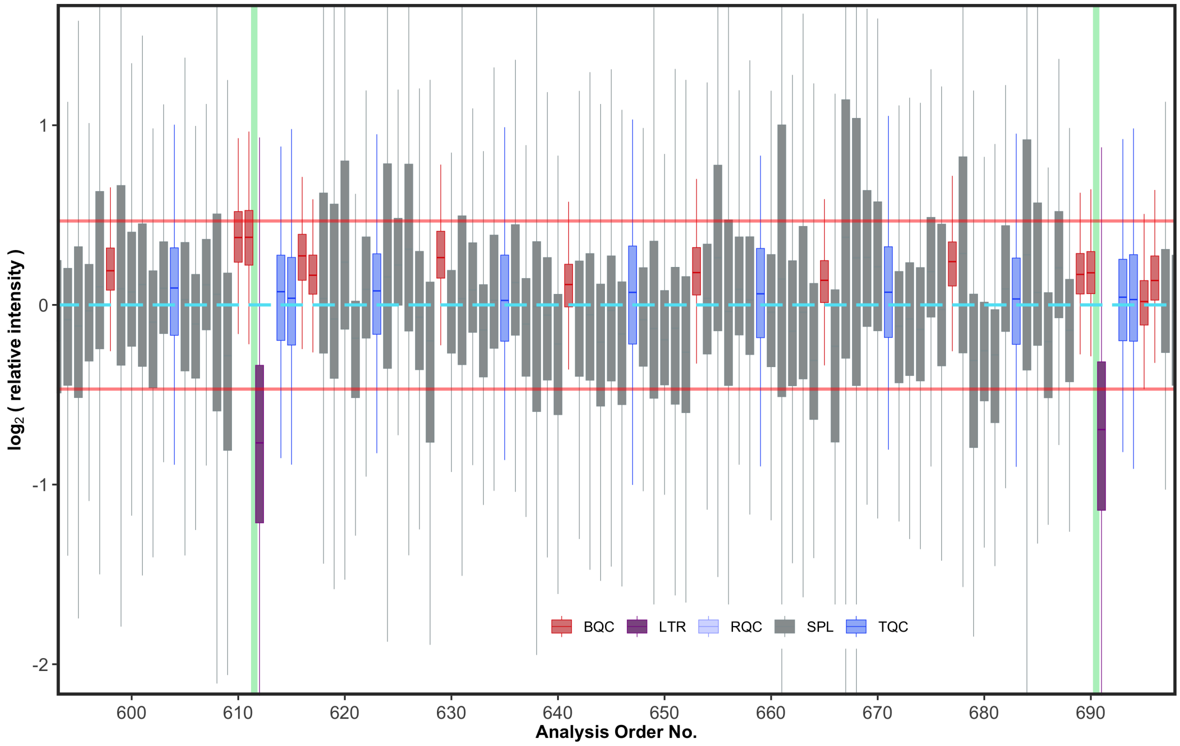

Now we plot the distributions of the ISTDs only. We see that also they also have an approximately 1/2 lower intensity in the LTR samples compared to the other samples. This may suggest that the measured sample amount was lower for the LTR samples, or that twice the amount of extraction solvent was added, both of which will lead to reduction of all features, including ISTDs.

```
extfig2b <- plot_rla_boxplot(
  data = mexp_initial,
  rla_type_batch = c("across"),
  variable = "intensity",
  qc_types = c("BQC", "SPL", "LTR", "TQC", "RQC", "PBLK"),
  filter_data = FALSE,
  include_feature_filter = "ISTD",
  plot_range =c(595,696),
  y_lim = c(-2,1.5),
  include_istd = TRUE,
  include_qualifier = FALSE,
  min_feature_intensity = 500,
  show_timestamp = FALSE,
  outlier_method = "iqr",
  outlier_k = 1.5,
```

```
  outlier_exclude = FALSE,
  x_gridlines = FALSE,
  batch_zebra_stripe = FALSE,
  show_plot = FALSE,
  base_font_size = 8,
  linewidth = 0.2
)$plot
#> i Found 13 outliers in the 919 shown analyses
extfig2b
```

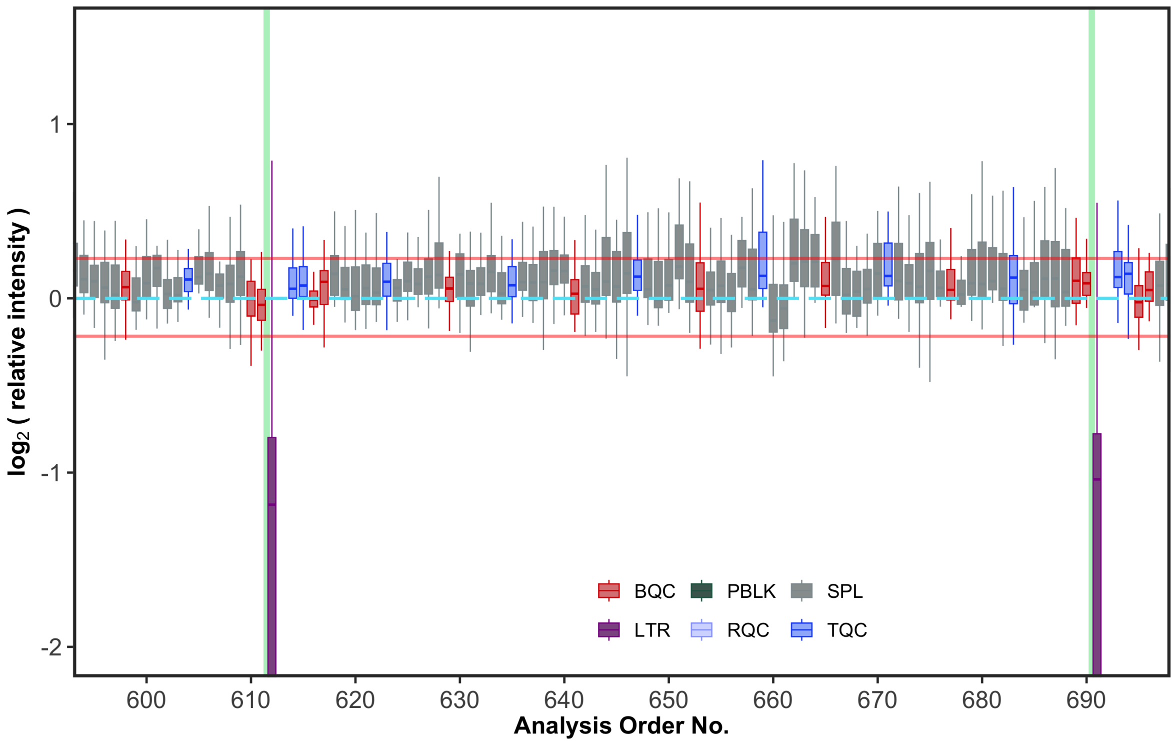

A further, detailed, batch-by-batch inspection, revealed that in batch 6, the sample "Longit_batch6_51" at position 464 had significantly lower intensities of non-ISTD feature, with intensities comparable to Blanks.

```
extfig2c <- plot_rla_boxplot(
  data = mexp_initial,
  rla_type_batch = c("across"),
  variable = "intensity",
  qc_types = c("BQC", "SPL", "LTR", "TQC", "RQC", "PBLK"),
  filter_data = FALSE,
```

```r
  plot_range = c(430,522),
  y_lim = c(-10,1.5),
  include_istd =  FALSE,
  exclude_feature_filter = "ISTD",
  include_qualifier = FALSE,
  min_feature_intensity = 500,
  show_timestamp = FALSE,
  outlier_method = "iqr",
  outlier_k = 1.5,
  outlier_exclude = FALSE,
  x_gridlines = FALSE,
  batch_zebra_stripe = FALSE,
  base_font_size = 6,
  show_plot = FALSE,
  linewidth = 0.2)$plot +
    theme(
      strip.text = ggplot2::element_text(size = 8),
      #aspect.ratio = 0.9,
      legend.position = "inside",
      axis.text = element_text(size = 8),
      #legend.direction = "vertical",
      legend.text = element_text(size = 8),
      legend.title = element_text(size = 8),
      legend.key.size = unit(8, "pt"),
      legend.position.inside = c(0.3, 0.32))
#> i Found 20 outliers in the 919 shown analyses
extfig2c
```

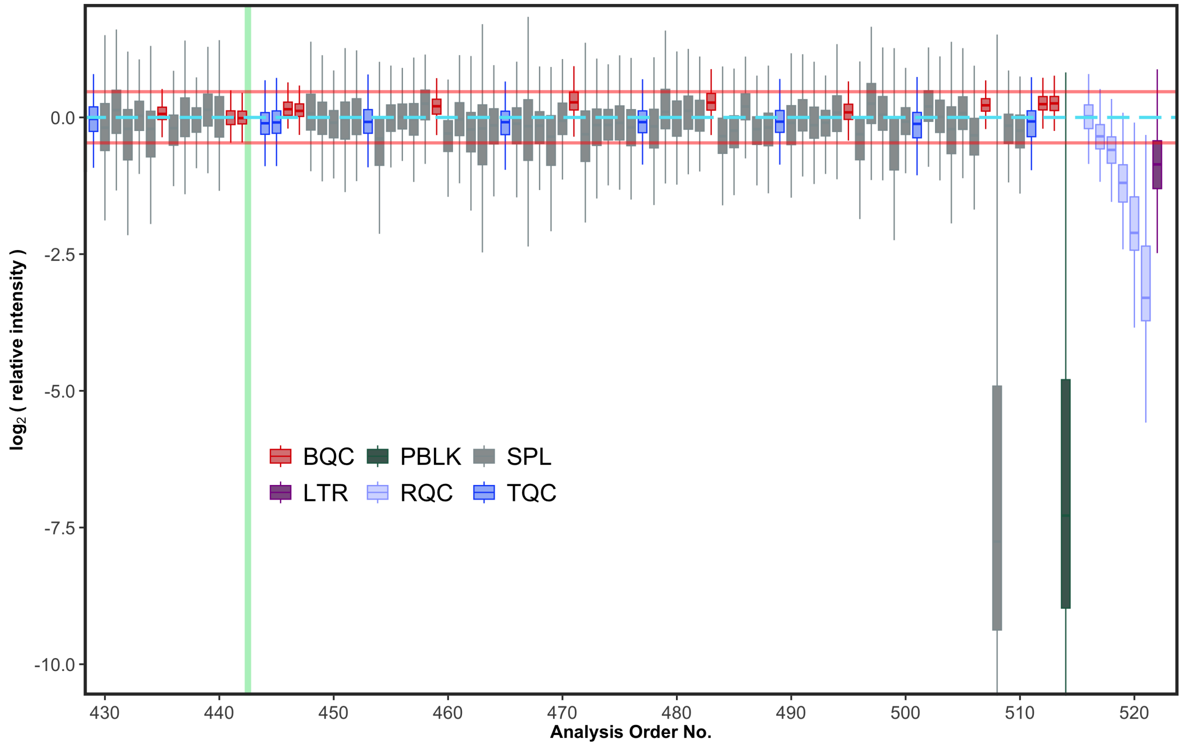

However, the ISTDs signals were comparable with other samples. This likely suggests that this study sample was not added to the extraction vial before extraction. The sample will be excluded from further analysis.

```
mexp_initial_temp2 <- mexp_initial
mexp_initial_temp2@dataset <- mexp_initial_temp2@dataset |> filter(is_istd)

extfig2d <- plot_rla_boxplot(
  data = mexp_initial_temp2,
  rla_type_batch = c("across"),
  variable = "intensity",
  qc_types = c("BQC", "SPL", "LTR", "TQC", "RQC", "PBLK"),
  filter_data = FALSE,
  plot_range = c(430,522),
  y_lim = c(-10, 1.5),
  include_istd = TRUE,
  include_feature_filter = "ISTD",
  include_qualifier = FALSE,
  min_feature_intensity = 500,
  show_timestamp = FALSE,
  outlier_method = "iqr",
```

```
  outlier_k = 1.5,
  outlier_exclude = FALSE,
  x_gridlines = FALSE,
  batch_zebra_stripe = FALSE,
  base_font_size = 8,
  show_plot = FALSE,
  linewidth = 0.2)$plot +
    theme(
    strip.text = ggplot2::element_text(size = 8),
    #aspect.ratio = 0.9,
    legend.position = "inside",
    axis.text = element_text(size = 8),
    #legend.direction = "vertical",
    legend.text = element_text(size = 8),
    legend.title = element_text(size = 8),
    legend.key.size = unit(8, "pt"),
    legend.position.inside = c(0.3, 0.32))
#> i Found 20 outliers in the 919 shown analyses
extfig2d
```

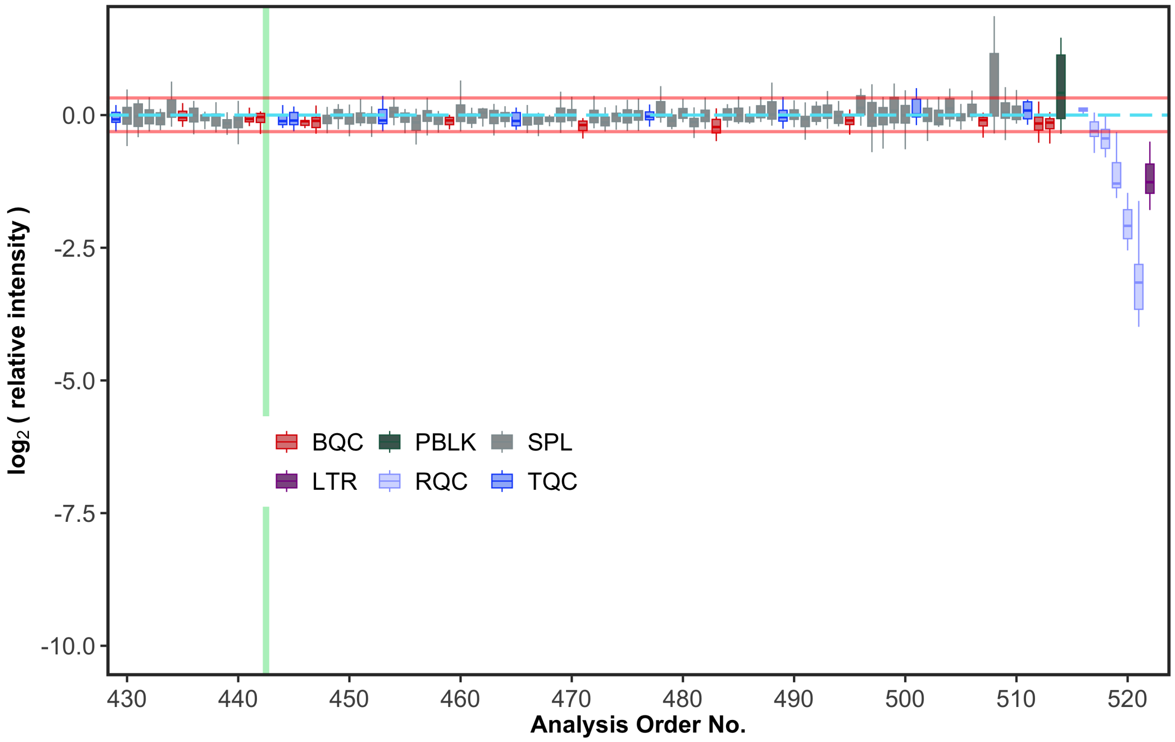

##### 3.3 Peak annotation QC

To check for potential peak picking errors during the peak integration step (see section above), the retention time (RT) of lipid features is plotted against the total carbon number and double bond number. This QC plot reveals several potential peak picking errors, especially among the SM, which are likely caused by erroneous picking of the isobaric isotopologue peak of PCs.

To check for potential peak picking errors during the peak integration step (see section above), the retention time (RT) of lipid features are plotted against the total carbon count and double bond number. The currently imported datas was the result of an interative process of data review using the QC plot and curation, this actual dataset corresponds to final dataset after curation.

Lipid species flagged as potential misannotations were manually inspected in the chromatograms and compared with an online resource for lipid RTs measured using the same LC-MS method (https://metabolomics.baker.edu.au/method/lipids). These features were deemed likely correct.

```
mexp_initial <- exclude_analyses(mexp_initial,
                                  analyses = "Longit_batch6_51",
                                  clear_existing = TRUE)
#> i 1 analyses were excluded for downstream processing. Please reprocess data.
mexp_temp <- mexp_initial
mexp_temp@dataset <- mexp_temp@dataset |> filter(str_detect(feature_id, "^PC \\d"))

fig4c <- plot_rt_vs_chain(
  mexp_temp,
  qc_types = "SPL",
  x_var = "total_c",
  base_font_size = 6) +
  theme(
      legend.position = "right",
      legend.direction = "vertical",
      legend.text = element_text(size = base_font_size*0.8),
      legend.title = element_text(size = base_font_size*0.8),
      legend.key.size = unit(base_font_size *0.8, "pt"),
      legend.position.inside = c(0.1, 0.7)
      )
#> i The following features were flagged as potential annotation outliers: PC 30:0 (b), PC 3
fig4c
```

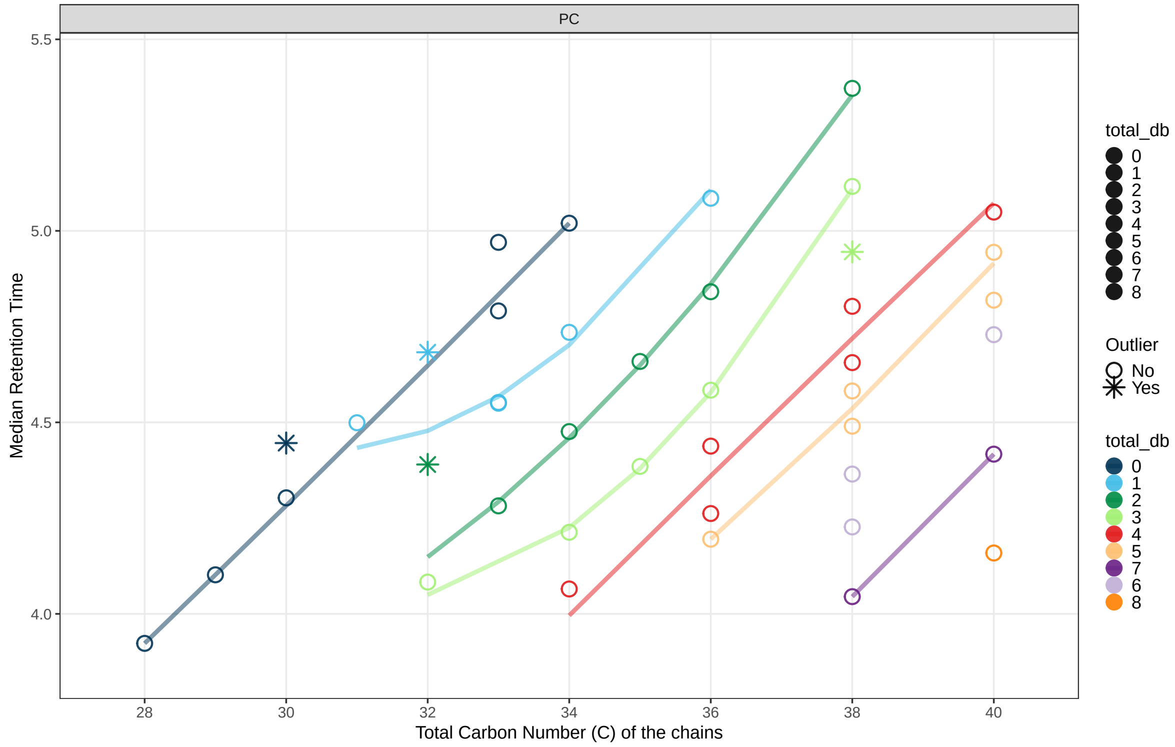

#### 3.4 7. Feature correlation analysis

As another check for potential peak picking errors, highly correlating features are plotted.

```
# this below is to exclude a sample that has a very low intensity for all features, see next
# details
#| fig-format: png
#| dpi: 600
mexp_initial <- exclude_analyses(
  mexp_initial,
  analyses = "Longit_batch6_51",
  clear_existing = TRUE)
#> i 1 analyses were excluded for downstream processing. Please reprocess data.
fig4d <- plot_feature_correlations(
  mexp_initial,
  variable = "intensity",
  include_feature_filter = c(
    "Cer d18:1/24:0", "Cer d18:0/24:1"),
  qc_types = c("SPL", "BQC", "TQC"),
  cor_min = 0.1,sort_by_corr = TRUE, return_plot = TRUE,show_progress = F,
```

```
  log_scale = FALSE, cols_page = 1, rows_page = 1,
  font_base_size = 6)[[1]] +
  theme(
    strip.text = ggplot2::element_text(size = 6),
    #aspect.ratio = 0.9,
    legend.position = "inside",
    axis.text = element_text(size = 5),
    legend.direction = "vertical",
    legend.text = element_text(size = 6*0.7),
    legend.title = element_text(size = 6*0.7),
    legend.key.size = unit(6 *0.7, "pt"),
    legend.position.inside = c(0.9, 0.12)
)
#> Generating plots (1 page)
fig4d
```

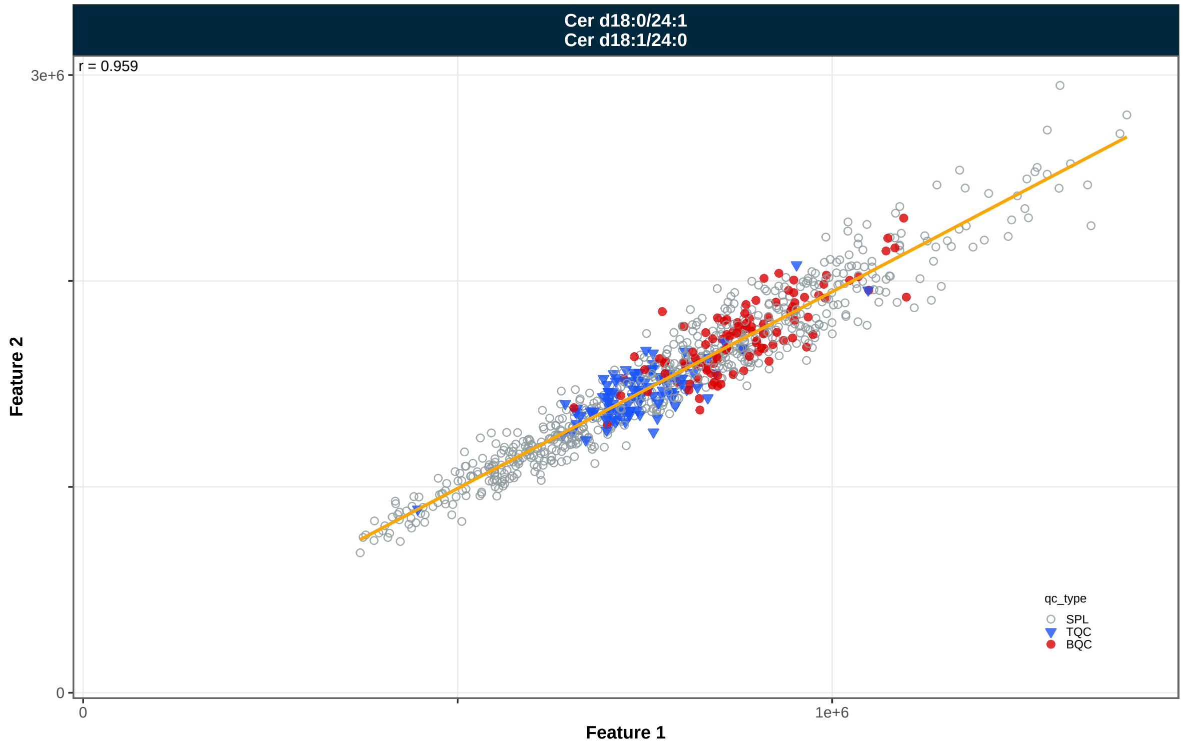

#### 3.5 8. Fix peak integration and re-import data (V2)

We now went back to the MRMkit workflow and corrected the peak integration errors The corrected data is available in the file `sPerfect_LongCirc_MRMkit.csv`. We import the corrected data, which we will use from now.

```
mexp <- MRMhubExperiment()
mexp <- import_data_mrmhub(
  mexp,
  "./data/dataset-1/data/sPerfect_LongCirc_MRMkit_Final.csv"
)
#> v Imported 937 analyses with 482 features
#> i `feature_area` selected as default feature intensity. Modify with `set_intensity_var()`
#> v Analysis metadata associated with 937 analyses.
#> v Feature metadata associated with 482 features.
```

We also directly plot the RT against chain lengths again, which confirms that the peak picking errors seems to have been corrected.

```
extfig3a <- plot_rt_vs_chain(mexp_initial, qc_types = "SPL", x_var = "total_c", base_font_si
  theme(
      legend.position = "right",
      legend.direction = "vertical",
      legend.text = element_text(size = base_font_size*0.8),
      legend.title = element_text(size = base_font_size*0.8),
      legend.key.size = unit(base_font_size *0.8, "pt"),
      legend.position.inside = c(0.1, 0.7))
#> i The following features were flagged as potential annotation outliers: DG 18:1_20:0 [-18
extfig3a
```

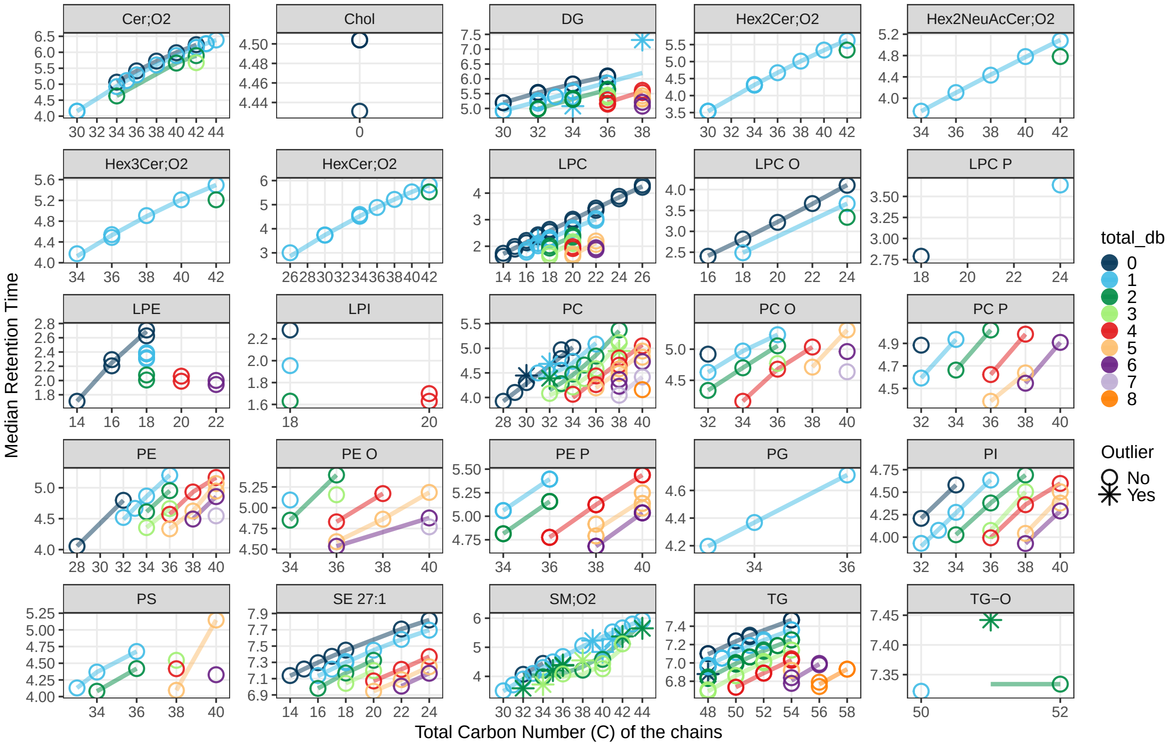

```
extfig3b <- plot_rt_vs_chain(mexp, qc_types = "SPL", x_var = "total_c", base_font_size = 6)+
  theme(
      legend.position = "right",
      legend.direction = "vertical",
      legend.text = element_text(size = base_font_size*0.8),
      legend.title = element_text(size = base_font_size*0.8),
      legend.key.size = unit(base_font_size *0.8, "pt"),
      legend.position.inside = c(0.1, 0.7))
#> i The following features were flagged as potential annotation outliers: LPC 17:1 (c), SM
extfig3b
```

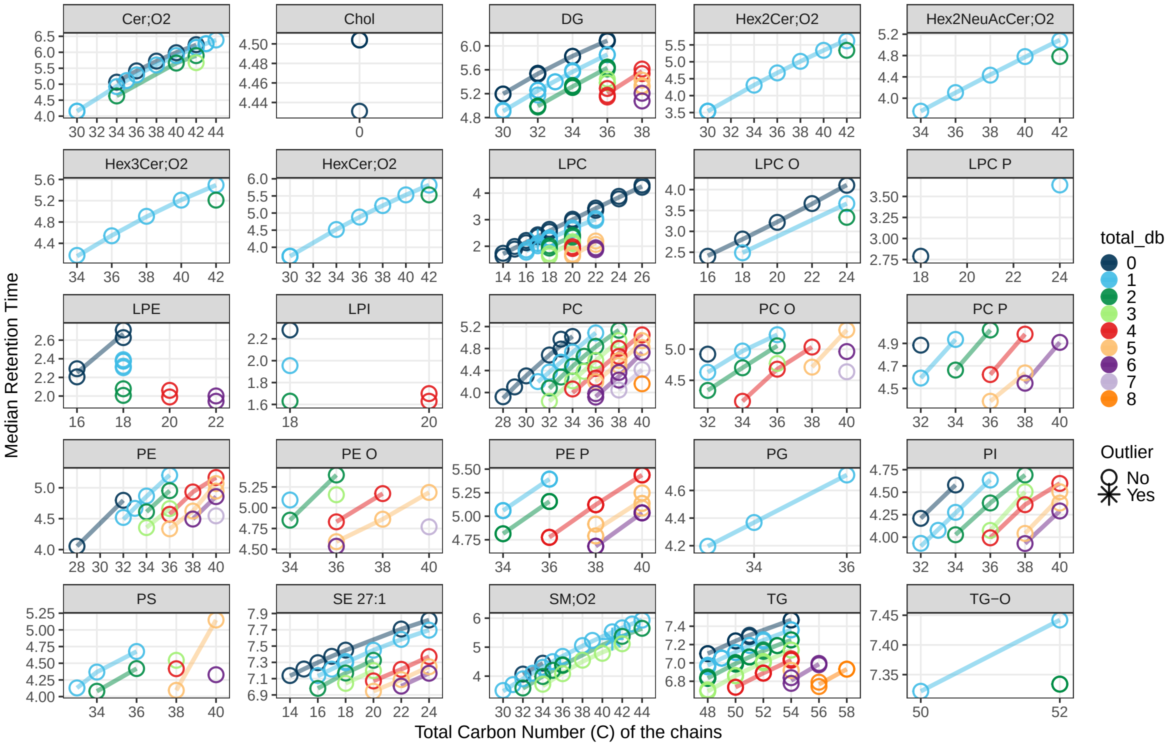

#### 3.6 PCA to check for potential technical outliers

```
extfig4a <- plot_pca(
  data = mexp,
  variable = "intensity",
  filter_data = FALSE,
  pca_dims = c(1,2),
  labels_threshold_mad = 4,
  qc_types = c("SPL", "BQC", "TQC", "LTR"),
  ellipse_variable = "qc_type",
  log_transform = TRUE,
  point_size = 1, point_alpha = 0.7, font_base_size = 8, ellipse_alpha = 0.3,
  include_istd = FALSE,show_labels = TRUE,label_font_size = 2,
  shared_labeltext_hide = NA)    + theme(
    plot.title = element_blank(),
    aspect.ratio = 1,
        legend.position = "inside",
      legend.direction = "vertical",
      legend.text = element_text(size = 9*0.7),
```

```
      legend.title = element_text(size = 9*0.7),
      legend.key.size = unit(7 * 0.7, "pt"),
      legend.position.inside = c(0.83, 0.83))
#> ! 9 features contained missing or non-numeric values and were exluded.
#> i The PCA was calculated based on `feature_intensity` values of 465 features.
extfig4a
```

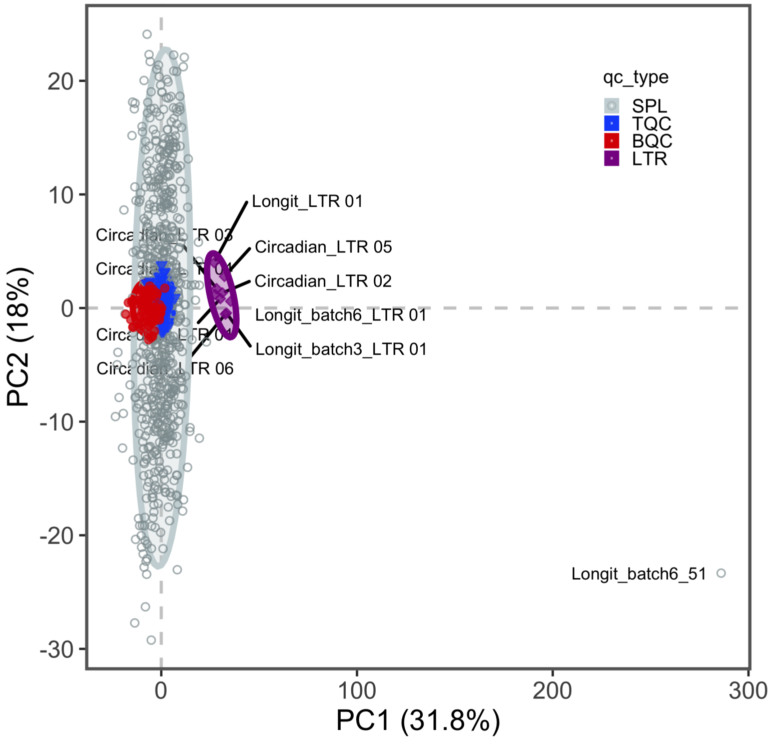

#### 3.7 Exclude outlier analysis/sample and replot PCA

```
mexp <- exclude_analyses(
  mexp,
  analyses = "Longit_batch6_51", clear_existing = FALSE)
#> i A total of 1 analyses are now excluded for downstream processing. Please reprocess data
fig4e <- plot_pca(
  data = mexp,
  variable = "intensity",
  filter_data = FALSE,
  pca_dims = c(1,2),
```

```
  labels_threshold_mad = 4,
  qc_types = c("SPL", "BQC", "TQC"),
  ellipse_variable = "qc_type",
  log_transform = TRUE,
  point_size = 1, point_alpha = 0.7, font_base_size = 6, ellipse_alpha = 0.3,
  include_istd = FALSE,show_labels = FALSE,label_font_size = 2,
  shared_labeltext_hide = NA)   +
    theme(
      plot.title = element_blank(),
      legend.position = "inside",
      legend.direction = "vertical",
      legend.text = element_text(size = 12*0.7),
      legend.title = element_text(size = 12*0.7),
      legend.key.size = unit(8 *0.7, "pt"),
      legend.position.inside = c(0.89, 0.89))
#> ! 9 features contained missing or non-numeric values and were exluded.
#> i The PCA was calculated based on `feature_intensity` values of 465 features.
fig4e
```

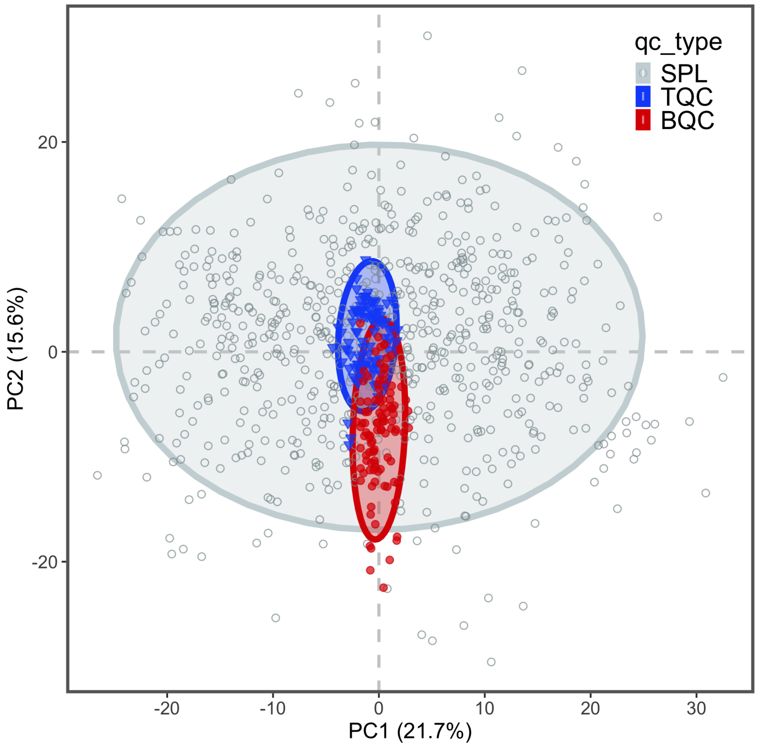

```r
extfig4b <- plot_pca_loading(
  data = mexp,
  variable = "feature_intensity",
  include_istd = FALSE,
  pca_dims = c(1,2,3,4),
  top_n = 70,
  font_base_size = 7,
  #qc_types = c("SPL", "BQC", "TQC", "LTR"),
  log_transform = TRUE)   +
    theme(
      plot.title = element_blank(),
      legend.position = "inside",
      legend.direction = "vertical",
      legend.text = element_text(size = 8*0.7),
      legend.title = element_text(size = 8*0.7),
      legend.key.size = unit(6 *0.7, "pt"),
      legend.position.inside = c(0.96, 0.06))
extfig4b
```

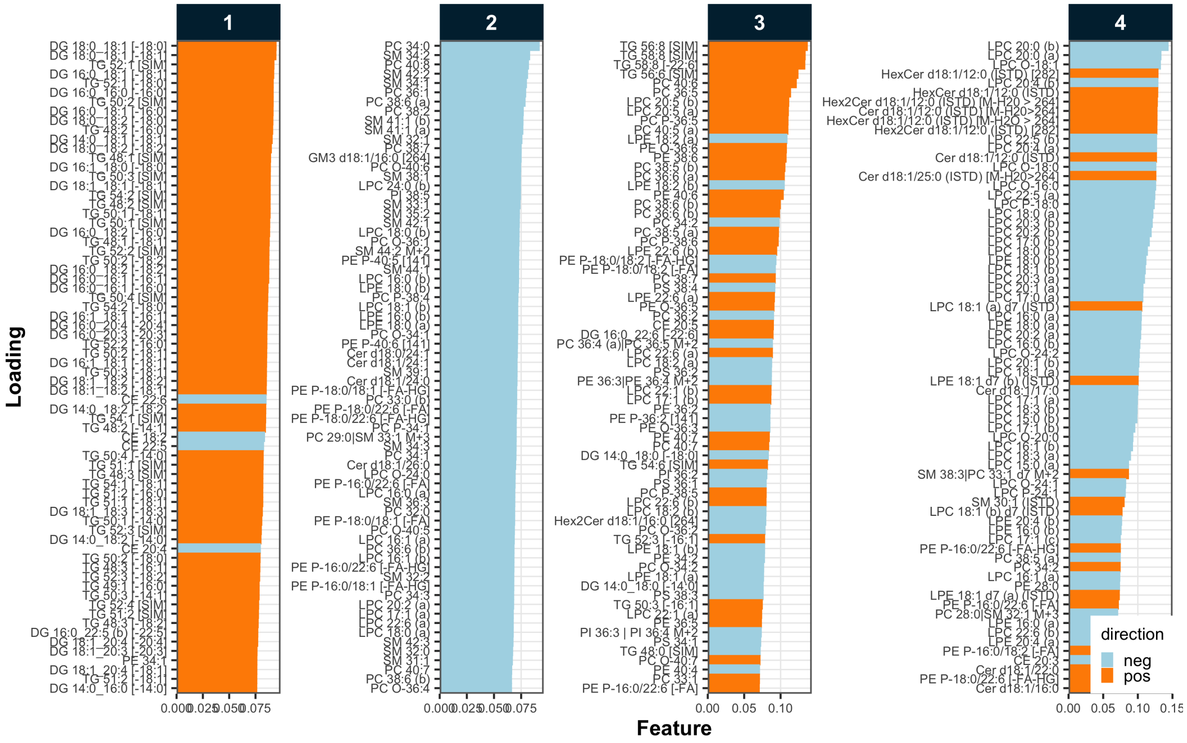

##### 3.8 11. Importing detailed metadata

To proceed with further processing, we require additional metadata describing the analyes/samples, features, internal standards (ISTDs), QC samples, and response curves. We are using the `MRMhub MSorganizer template`, an Excel template that provides a user-friendly solution to collect, organise and pre-validate analysis metadata. However, other formats to import metatadata are available as well, such as from CSV files, individual Excel tables, or R data frames.

After metadata import, a summary of the validation of data/metadata integrity is returned. For example, duplicate or missing data, such as IDs will be highlighted. The user can chose to ignore warnings (`ignore_warnings = TRUE`) and proceed with the analysis. Errors however, such as missing IDs or duplicate IDs must always be fixed before analysis can proceed.

```
file_path <- "./data/dataset-1/data/sPerfect_LongCirc_Metadata.xlsm"
mexp <- import_metadata_msorganiser(mexp, file_path, ignore_warnings = TRUE)
#> ! Metadata has following warnings and notifications:
#> ------------------------------------------------------------------------------------------------
#>   Type Table    Column     Issue                            Count
#> 1 W*   Features feature_id Feature(s) not in analysis data    28
#> 2 W*   Features feature_id Feature(s) without metadata         5
#> ------------------------------------------------------------------------------------------------
#> E = Error, W = Warning, W* = Supressed Warning, N = Note
#> ------------------------------------------------------------------------------------------------
#> v Analysis metadata associated with 937 analyses.
#> v Feature metadata associated with 477 features.
#> i 4 invalid features (as defined in the metadata) were excluded
#> v Internal Standard metadata associated with 18 ISTDs.
#> v Response curve metadata associated with 24 annotated analyses.
```

##### 3.9 12. RunScatter plots

We now plot the peak areas (`intensity`) of all ISTD features across all analyses/samples.

```
fig4f <- plot_runscatter(mexp, variable = "intensity",
                #include_feature_filter = "ISTD",
                qc_types = c("SPL", "BQC", "TQC", "PBLK", "RQC", "SBLK"),
                include_feature_filter = example_species_istd,
                show_reference_lines = TRUE,
                ref_qc_types = "SPL",
                reference_fill_color = "#111111",
                reference_k_sd = NA,
```

```r
                 point_size = 1,
                 base_font_size = 6,
                 cols_page = 1,
                 rows_page = 1,
                 cap_outliers = TRUE,
                 reference_sd_shade = FALSE,
                 #batch_zebra_stripe = TRUE,
                 output_pdf = FALSE,
                 show_progress = FALSE,
                 path = "rt_all.pdf",
                 return_plots = TRUE)[[1]]+
  theme(
    legend.position = "inside",
    legend.direction = "horizontal",
    legend.text = element_text(size = base_font_size*0.8),
    legend.title = element_text(size = base_font_size*0.8),
    legend.key.size = unit(base_font_size *0.8, "pt"),
    legend.position.inside = c(0.7, 0.1))
#> Generating plots (1 page)
```

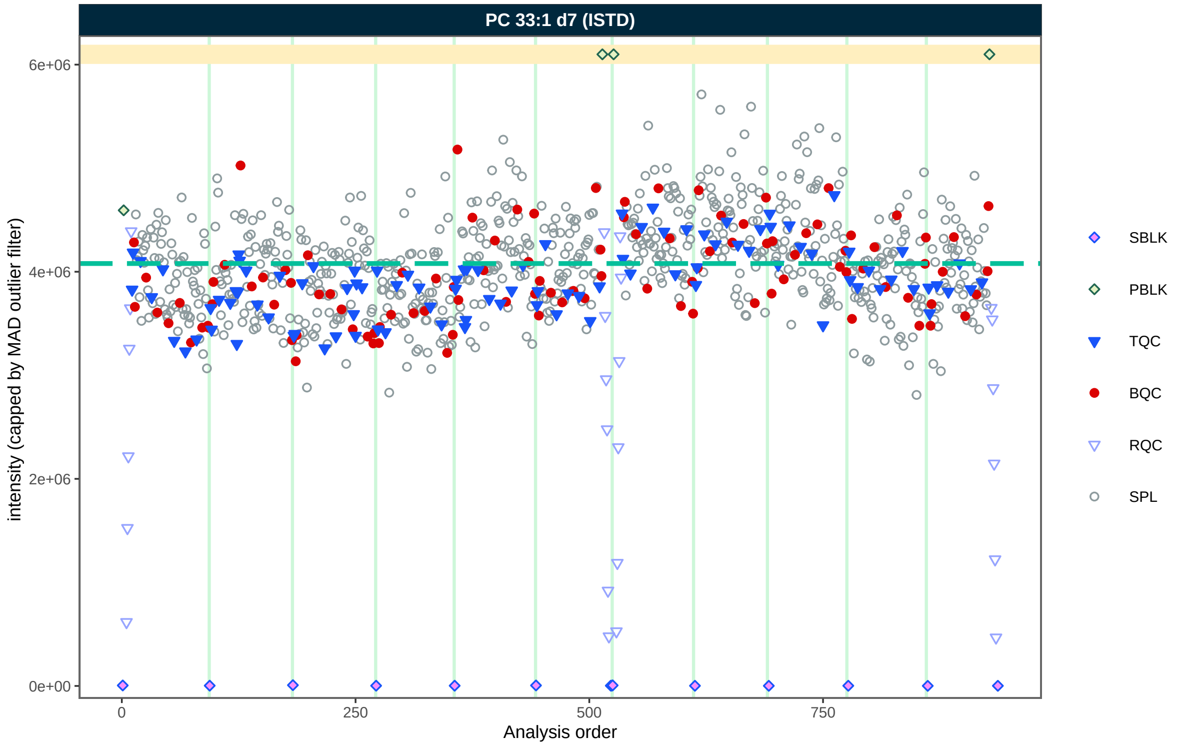

Additionally, we plot all ISTD features in a multi-page PDF to check for potential issues with individual ISTDs.

```
extfig5 <- plot_runscatter(mexp, variable = "intensity",
                include_feature_filter = "ISTD", include_qualifier = F,
                qc_types = c("SPL", "BQC", "TQC", "PBLK", "RQC", "SBLK"),
                show_reference_lines = TRUE,
                ref_qc_types = "SPL",
                reference_fill_color = "#111111",
                reference_k_sd = NA,
                point_size = .7,
                point_border_width = 0.2,
                point_transparency = .7,
                base_font_size = 6,
                cols_page = 4,
                rows_page = 5,
                cap_outliers = TRUE,
                reference_sd_shade = FALSE,
                #batch_zebra_stripe = TRUE,
                show_progress = FALSE,
                output_pdf = FALSE,
                path = "output/area_istd_all.pdf", return_plots = TRUE)
#> Generating plots (1 page)
```

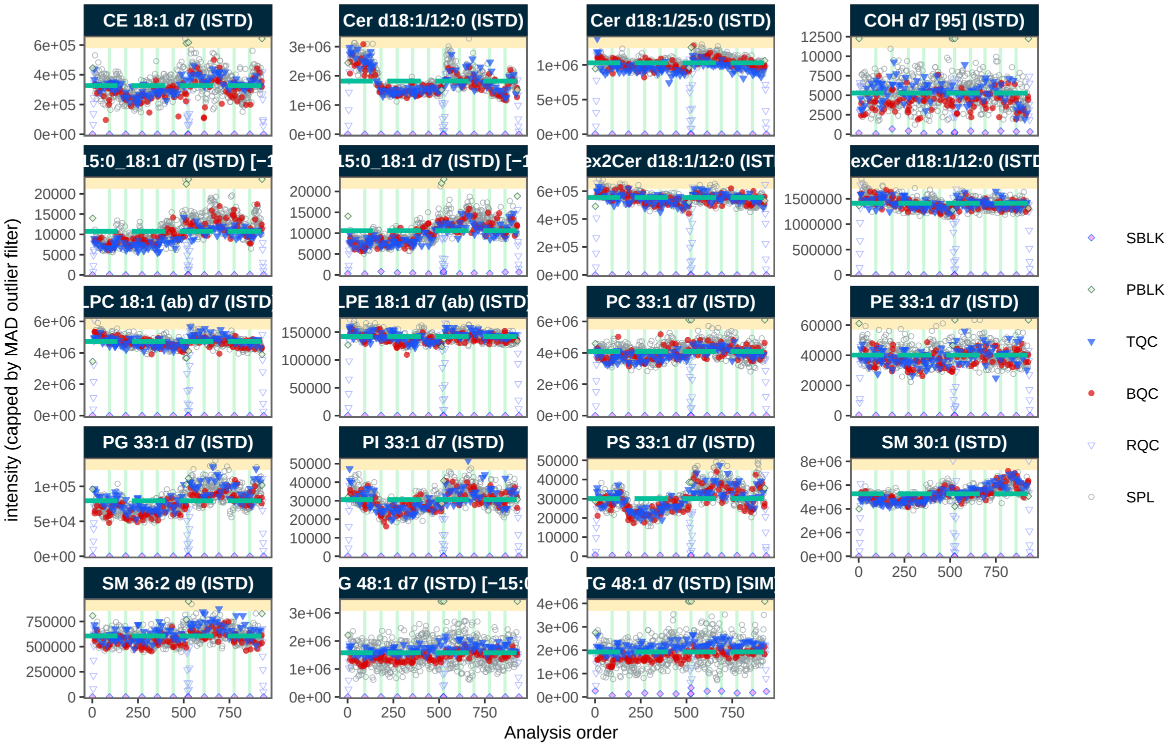

#### 3.10 13. Matrix effects

To check for potential matrix effects, we plot the peak areas (`intensity`) of all ISTD featuresin different QC types.

```
mexp_temp <- mexp
mexp_temp@dataset <- mexp_temp@dataset |> filter(!str_detect(feature_id, "DG 15:0\\_18:1 d7

fig4g <- plot_qc_matrixeffects(
  mexp_temp,
  variable = "intensity",
  y_lim = c(0,250),
  point_alpha = 0.55,
  box_alpha = 0.3,
  point_size = 0.1,
  box_linewidth = 0.3,
  font_base_size = 6,
  include_feature_filter = "P[CIE]|DG|TG|Cer|CE",
  min_median_value = 3000)
```

```
fig4g
```

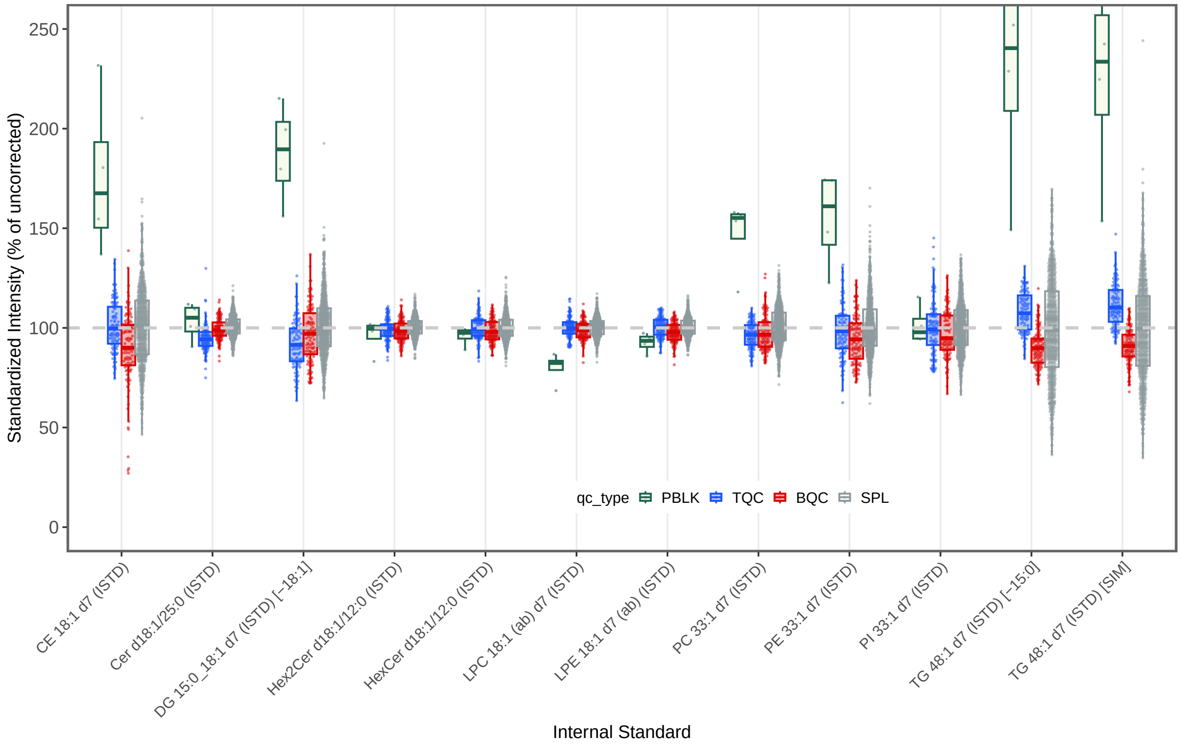

#### 3.11 14. Normalization and Quantification

```
mexp <- normalize_by_istd(mexp)
#> ! Interfering features defined in metadata, but no correction was applied. Use `correct_i
#> v 435 features normalized with 17 ISTDs in 937 analyses.
mexp <- quantify_by_istd(mexp)
#> v 449 feature concentrations calculated based on 28 ISTDs and sample amounts of 937 analy
#> i Concentrations are given in  mol/L.
```

#### 3.12 15. Drift and Batch Correction

```
mexp <- correct_drift_gaussiankernel(
  mexp,
```

```
  variable = "conc",
  ref_qc_types = "SPL",
  batch_wise = TRUE,
  kernel_size = 20,
  outlier_filter = TRUE,
  outlier_ksd = 3,
  recalc_trend_after = TRUE,
  show_progress = FALSE)
#> i Applying `conc` drift correction...
#> i 6 feature(s) contain one or more zero or negative `conc` values. Verify your data or use
#> ! 3 features showed no variation in the study sample's original values across analyses.
#> ! 3 features have invalid values after smoothing. NA will be be returned for all values of
#> v Drift correction was applied to 446 of 449 features (batch-wise).
#> i The median CV change of all features in study samples was -0.52% (range: -7.93% to 1.56%
mexp <- correct_batch_centering(
  mexp,
  ref_qc_types = "SPL",
  log_transform_internal = FALSE,
  variable = "conc")
#> i Adding batch correction on top of `conc` drift-correction.
#> v Batch median-centering of 11 batches was applied to drift-corrected concentrations of al
#> i The median CV change of all features in study samples was -0.97% (range: -30.20% to 50.8
```

##### 3.13 16. Drift and Batch Correction

```
fig5c <- plot_runscatter(mexp, variable = "conc_raw",
                #include_feature_filter = "ISTD",
                qc_types = c("SPL", "BQC", "TQC", "LTR"),
                include_feature_filter = example_species,
                #y_min = 0.00,
                #y_max = 0.15,
                #plot_range = c(0, 910),
                show_reference_lines = TRUE,ref_qc_types = "SPL",
                reference_fill_color = "#b83c3cff",
                reference_line_color = "#37c2f0ff",
                reference_k_sd = NA,
                show_trend = TRUE,
                point_size = 1,
                base_font_size = 6,
                cols_page = 1,
```

```
                rows_page = 1,
                cap_outliers = FALSE,
                reference_sd_shade = FALSE,
                #batch_zebra_stripe = TRUE,
                show_progress = FALSE,
                output_pdf = FALSE,
                path = "rt_all.pdf", return_plot = TRUE)[[1]]+
  theme(
    legend.position = "inside",
    legend.direction = "horizontal",
    legend.text = element_text(size = base_font_size*0.8),
    legend.title = element_text(size = base_font_size*0.8),
    legend.key.size = unit(base_font_size *0.8, "pt"),
    legend.position.inside = c(0.7, 0.1))
#> Generating plots (1 page)
```

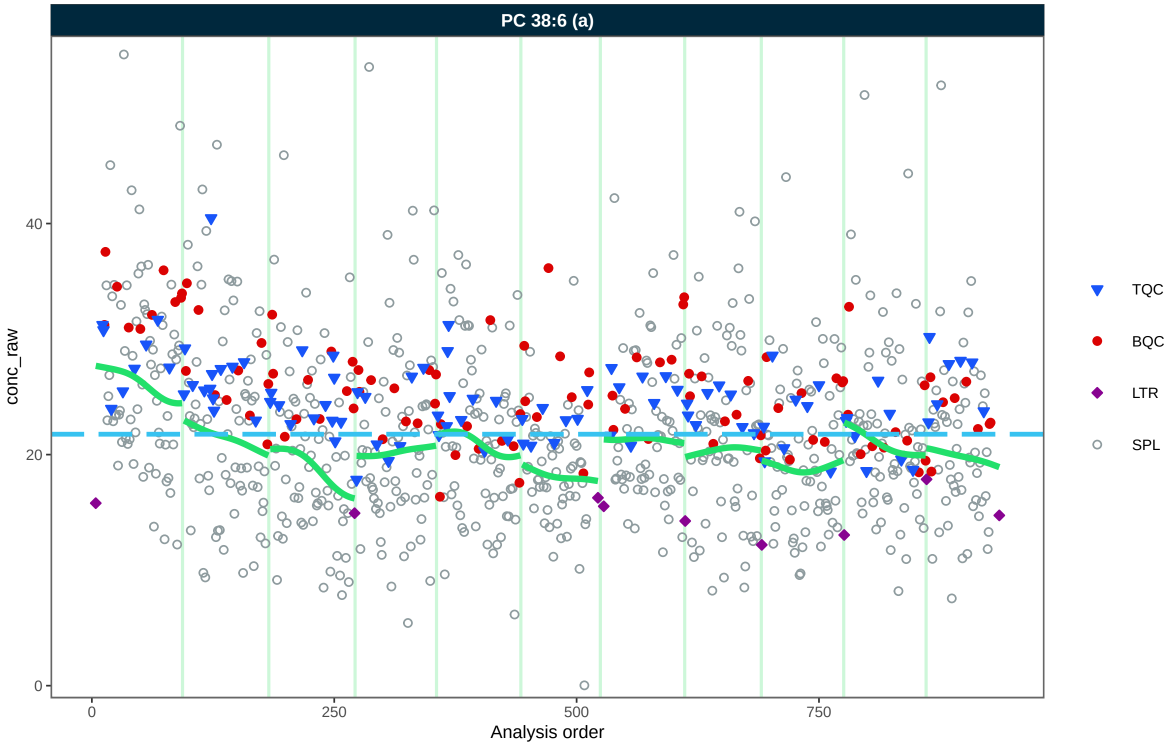

```
fig5d <- plot_runscatter(mexp, variable = "conc",
                #include_feature_filter = "ISTD",
                qc_types = c("SPL", "BQC", "TQC", "LTR"),
                include_feature_filter = example_species,
```

```
                 #y_min = 0.00,
                 #y_max = 0.9,
                 #plot_range = c(0, 910),
                 show_reference_lines = FALSE,
                 ref_qc_types = "SPL",
                 reference_fill_color = "#111111",
                 reference_line_color = "#37c2f0ff",
                 reference_k_sd =  NA,
                 show_trend = TRUE,
                 point_size = 1,
                 base_font_size = 6,
                 cols_page = 1,
                 rows_page = 1,
                 cap_outliers = FALSE,
                 reference_sd_shade = FALSE,
                 #batch_zebra_stripe = TRUE,
                 show_progress = FALSE,
                 output_pdf = FALSE,
                 path = "rt_all.pdf", return_plot = TRUE)[[1]]+
  theme(
    legend.position = "inside",
    legend.direction = "horizontal",
    legend.text = element_text(size = base_font_size*0.8),
    legend.title = element_text(size = base_font_size*0.8),
    legend.key.size = unit(base_font_size *0.8, "pt"),
    legend.position.inside = c(0.7, 0.1))
#> Generating plots (1 page)
```

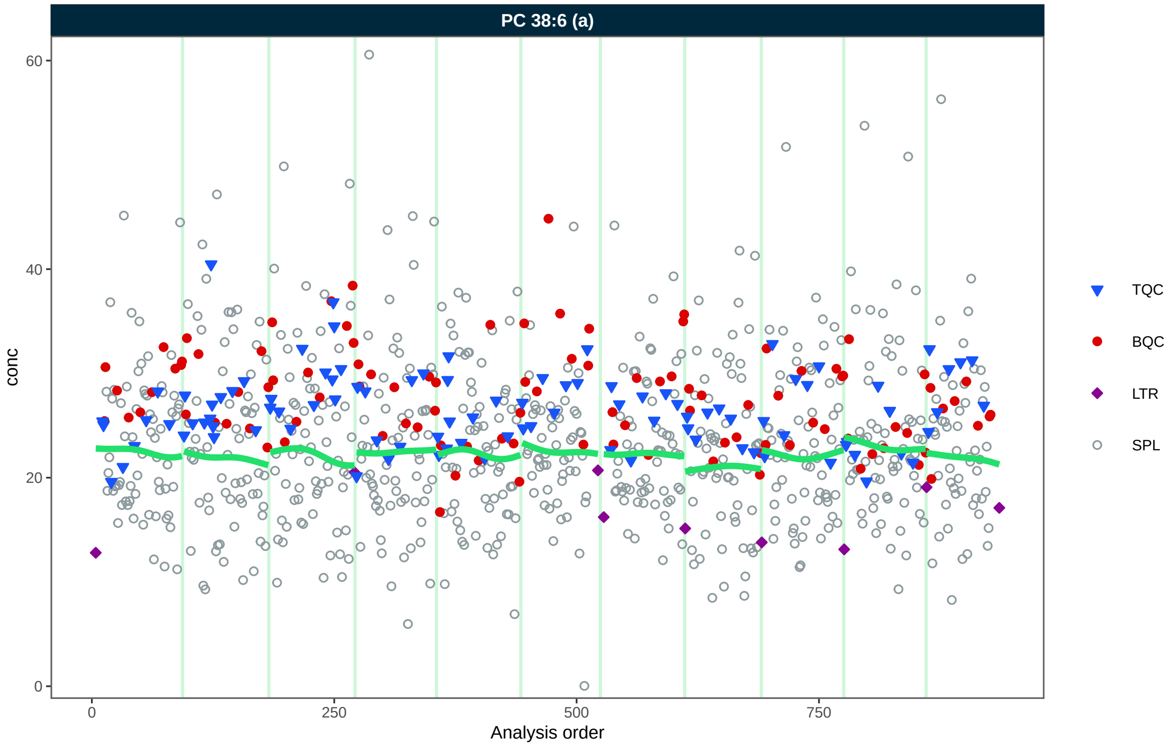
PC 38:6 (a)
conc
Analysis order
0
20
40
60
0
250
500
750
TQC
BQC
LTR
SPL

```
fig5d
```

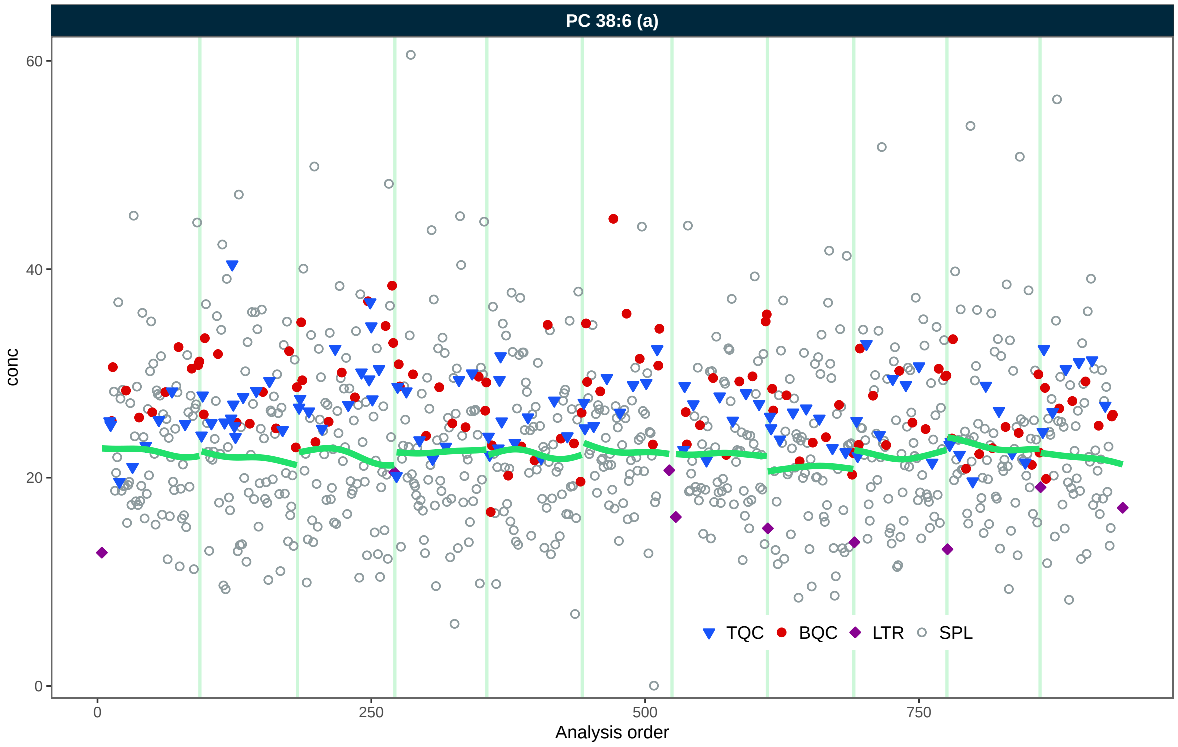

#### 3.14 17. Normalization and Correction QC

```
mexp_t1 <- calc_qc_metrics(mexp, use_robust_cv = TRUE, use_batch_medians = FALSE)
mexp_t1@metrics_qc <- mexp_t1@metrics_qc |>
  dplyr::filter(str_detect(feature_class, "^PC$|TG$"))

mexp_t2 <- calc_qc_metrics(mexp)
mexp_t2@metrics_qc <- mexp_t2@metrics_qc |>
  dplyr::filter(str_detect(feature_class, "^PC$|TG$"))

fig5a <- plot_normalization_qc(
  plot_type = "diff",
  data = mexp_t1,
  before_norm_var = "intensity",
  after_norm_var = "norm_intensity",
  y_lim = c(-15,20),
  x_lim = c(0,75),
  qc_types = c("TQC", "BQC", "SPL"),
  cols_page = 3,
```

```r
  font_base_size = 5,
  point_size = 1,
  facet_by_class = TRUE,
  include_qualifier = FALSE) +
    theme(
      strip.text = ggplot2::element_text(size = 5),
      #aspect.ratio = 0.9,
      legend.position = "inside",
      legend.position.inside = c(0.77, 0.27),
      axis.text = element_text(size = 4),
      #axis.title = element_blank(),
      legend.direction = "vertical",
      legend.text = element_text(size = 6*0.7),
      legend.title = element_text(size = 6*0.7),
      legend.key.size = unit(6 *0.7, "pt"))

fig5b <- plot_normalization_qc(
  data = mexp_t2,
  plot_type = "diff",
  before_norm_var = "norm_intensity",
  after_norm_var = "conc",
  y_lim = c(-15,20),
  x_lim = c(0,75),
  qc_types = c("TQC", "BQC", "SPL"),
  cols_page = 2,
  facet_by_class = TRUE,
  font_base_size = 5,
  point_size = 1,
  include_qualifier = FALSE) +
    theme(
      strip.text = ggplot2::element_text(size = 5),
      #aspect.ratio = 0.9,
      legend.position = "inside",
      legend.position.inside = c(0.77, 0.27),
      axis.text = element_text(size = 4),
      #axis.title = element_blank(),
      legend.direction = "vertical",
      legend.text = element_text(size = 6*0.7),
      legend.title = element_text(size = 6*0.7),
      legend.key.size = unit(6 *0.7, "pt"))
fig5a
```

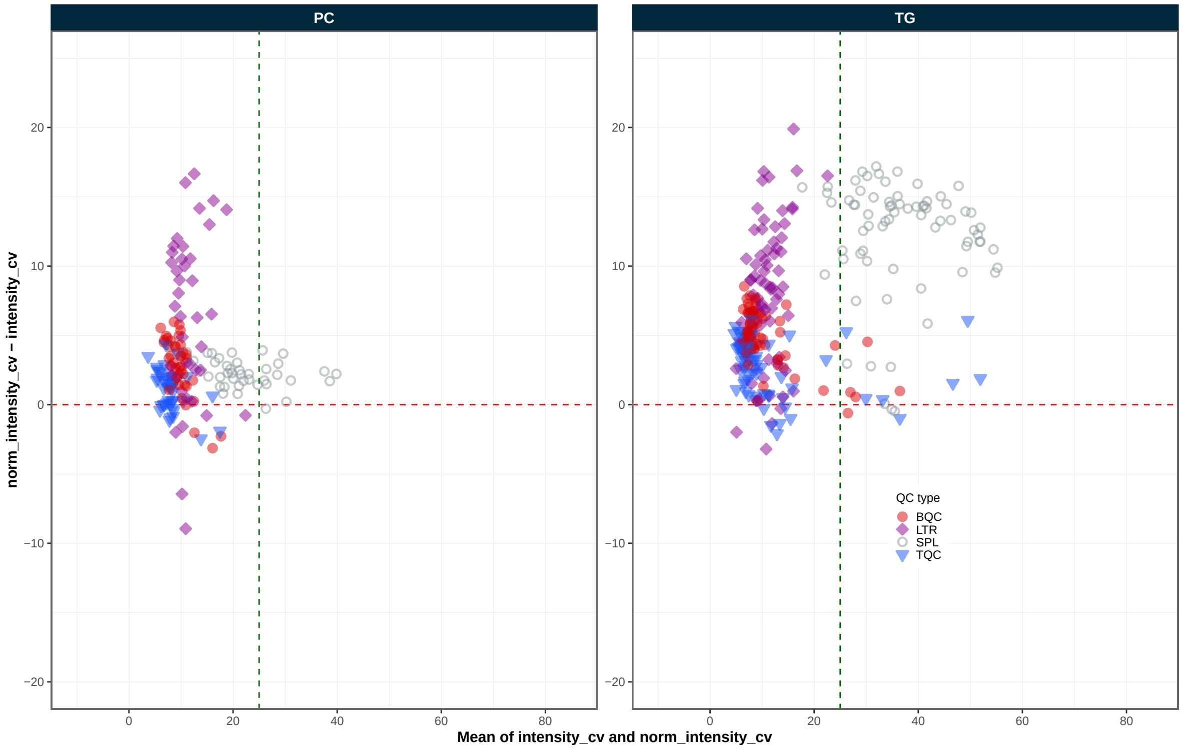
PC
TG
norm_intensity_cv – intensity_cv
Mean of intensity_cv and norm_intensity_cv
QC type
BQC
LTR
SPL
TQC
20
10
0
−10
−20
0
20
40
60
80

```
fig5b
```

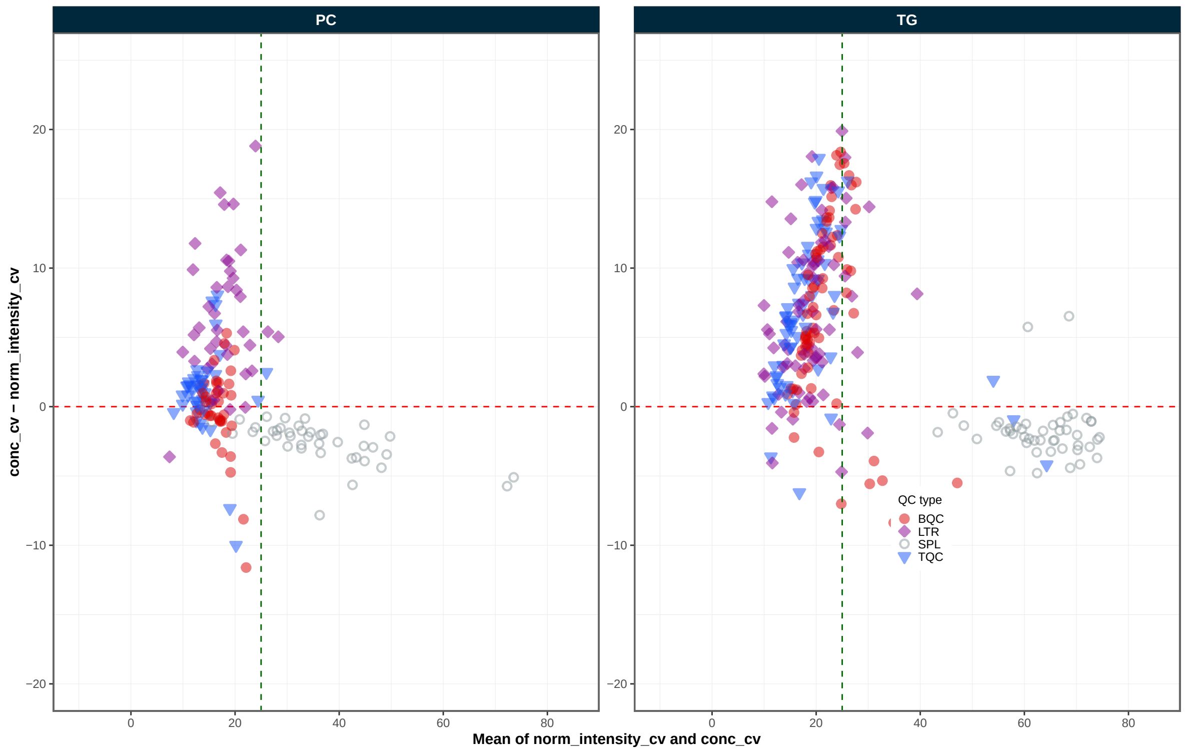

```r
mexp_temp <- calc_qc_metrics(mexp, use_robust_cv = TRUE,use_batch_medians = FALSE)

mexp_temp@metrics_qc <- mexp_temp@metrics_qc |>
  dplyr::filter(!str_detect(feature_id, "^TG O|COH|PG"))

extfig6a <- plot_normalization_qc(
  data = mexp_temp,
  before_norm_var = "intensity",
  after_norm_var = "norm_intensity",
  plot_type = "diff",
  y_lim = c(-15,20),
  x_lim = c(0,60),
  qc_types = c("TQC", "BQC", "SPL"),
  cols_page = 9,
  font_base_size = 5,
  point_size = 0.85,
  facet_by_class = TRUE,
  include_qualifier = FALSE) +
    theme(
      strip.text = ggplot2::element_text(size = 5),
```

```
      legend.position = "inside",
      legend.position.inside = c(0.9, 0.10),
      axis.text = element_text(size = 4),
      legend.direction = "vertical",
      legend.text = element_text(size = 6*0.7),
      legend.title = element_text(size = 6*0.7),
      legend.key.size = unit(6 *0.7, "pt"))
extfig6a
```

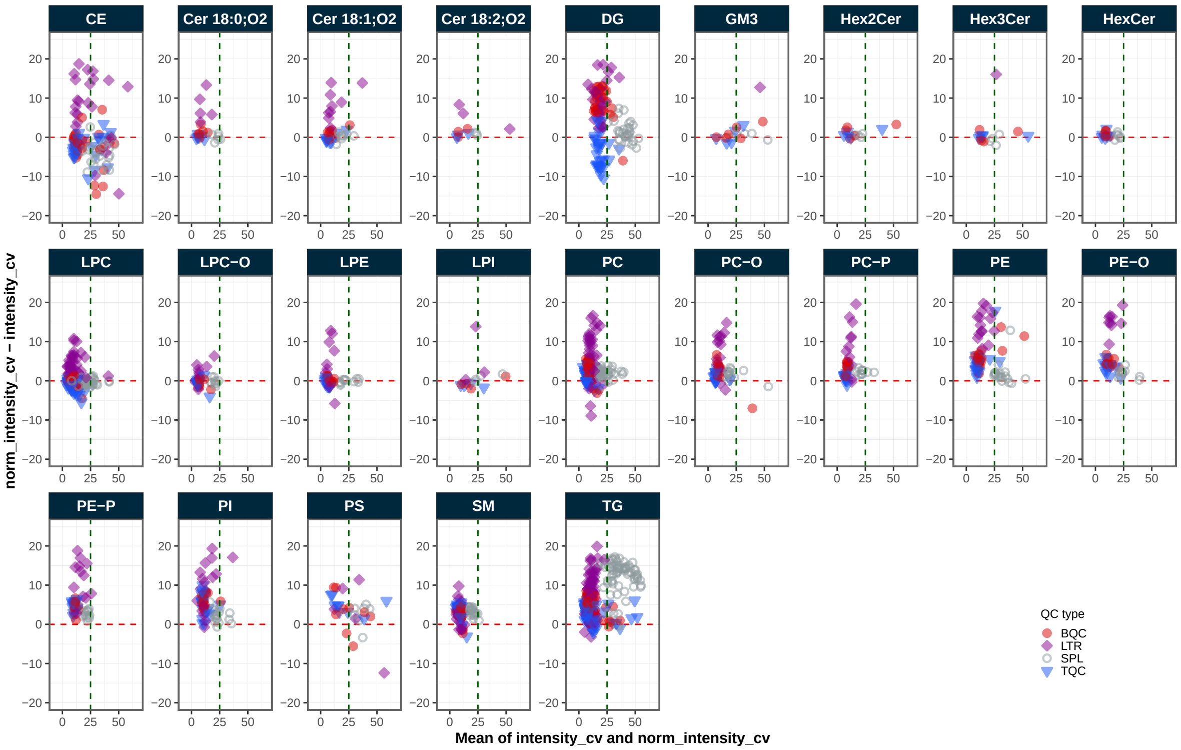

```
extfig6b <- plot_normalization_qc(
  data = mexp_temp,
  before_norm_var = "norm_intensity",
  after_norm_var = "conc",
  plot_type = "diff",
  y_lim = c(-15,20),
  x_lim = c(0,60),
  qc_types = c("TQC", "BQC", "SPL"),
  cols_page = 9,
  font_base_size = 5,
  point_size = 0.85,
```

```
  facet_by_class = TRUE,
  include_qualifier = FALSE) +
    theme(
      strip.text = ggplot2::element_text(size = 5),
      #aspect.ratio = 0.9,
      legend.position = "inside",
      legend.position.inside = c(0.9, 0.10),
      axis.text = element_text(size = 4),
      #axis.title = element_blank(),
      legend.direction = "vertical",
      legend.text = element_text(size = 6*0.7),
      legend.title = element_text(size = 6*0.7),
      legend.key.size = unit(6 *0.7, "pt"))
extfig6b
```

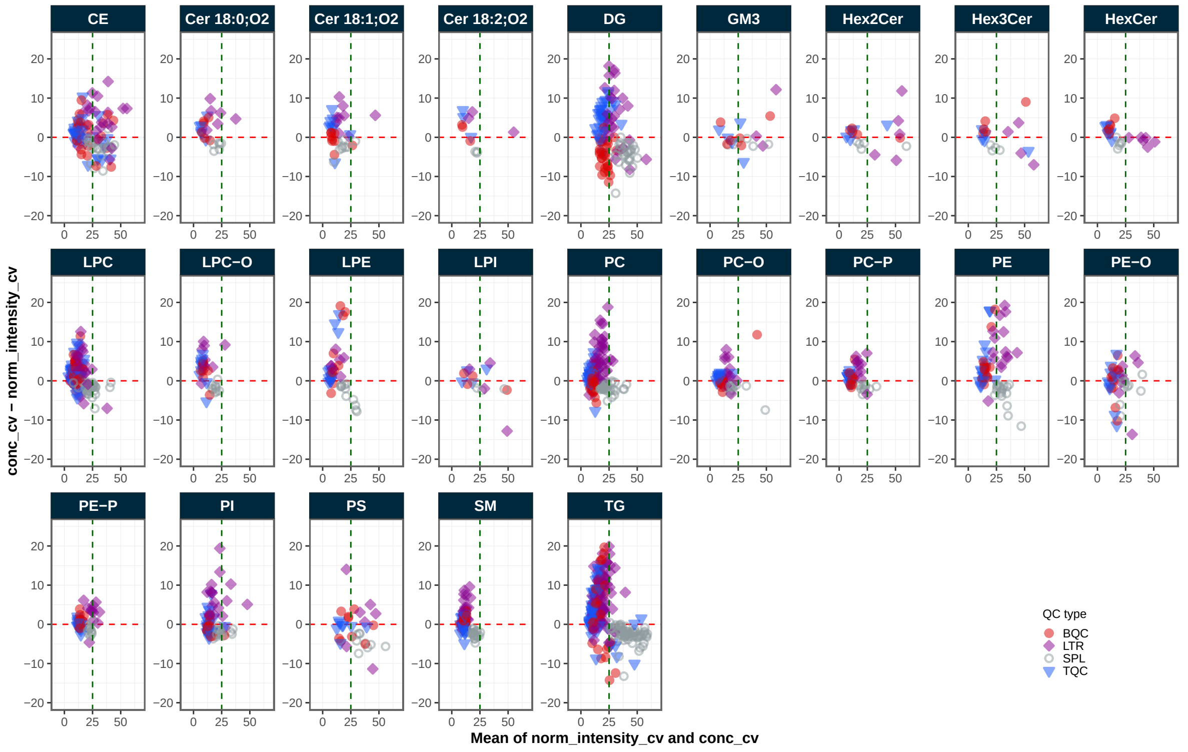

```
mexp_spline <- correct_drift_cubicspline(
  mexp,
  variable = "conc",
  ref_qc_types = "BQC",
  batch_wise = TRUE,
```

```
  lambda = 0.5,
  cv = TRUE,
  recalc_trend_after = TRUE, replace_previous = TRUE)
#> i Replacing previous `conc` drift and batch corrections...
#> i 6 feature(s) contain one or more zero or negative `conc` values. Verify your data or use
#> ! 3 features showed no variation in the study sample's original values across analyses.
#> ! 4 features have invalid values after smoothing. NA will be be returned for all values of
#> ! Smoothing failed for 4 feature(s) in all batches. Please check data, metadata, and fit p
#> ! Smoothing failed for 6 feature(s) in at least one batch: CE 15:0, COH [161], LPI 18:0, I
#> v Drift correction was applied to 445 of 449 features (batch-wise).
#> i The median CV change of all features in study samples was 0.08% (range: -3.49% to 14.38%
mexp_spline <- correct_batch_centering(
  mexp_spline,
  ref_qc_types = "BQC",
  variable = "conc")
#> i Adding batch correction on top of `conc` drift-correction.
#> v Batch median-centering of 11 batches was applied to drift-corrected concentrations of al
#> i The median CV change of all features in study samples was 0.20% (range: -28.20% to 121.9
mexp_spline <- calc_qc_metrics(
  mexp_spline,
  use_robust_cv = TRUE,
  use_batch_medians = FALSE)

mexp_spline@metrics_qc <- mexp_spline@metrics_qc |>
  dplyr::filter(!str_detect(feature_id, "^TG O|COH|PG"))

extfig6c <- plot_normalization_qc(
  data = mexp_spline,plot_type = "diff",
  before_norm_var = "norm_intensity",
  after_norm_var = "conc",
  y_lim = c(-20,20),
  x_lim = c(0,60),
  qc_types = c("TQC", "BQC", "SPL"),
  cols_page = 9,
  font_base_size = 5,cv_threshold_value = 25,
  point_size = 0.85,
  facet_by_class = TRUE,
  include_qualifier = FALSE) +
    theme(
      strip.text = ggplot2::element_text(size = 5),
      #aspect.ratio = 0.9,
      legend.position = "inside",
```

```
      legend.position.inside = c(0.9, 0.10),
      axis.text = element_text(size = 4),
      #axis.title = element_blank(),
      legend.direction = "vertical",
      legend.text = element_text(size = 6*0.7),
      legend.title = element_text(size = 6*0.7),
      legend.key.size = unit(6 *0.7, "pt"))
extfig6c
```

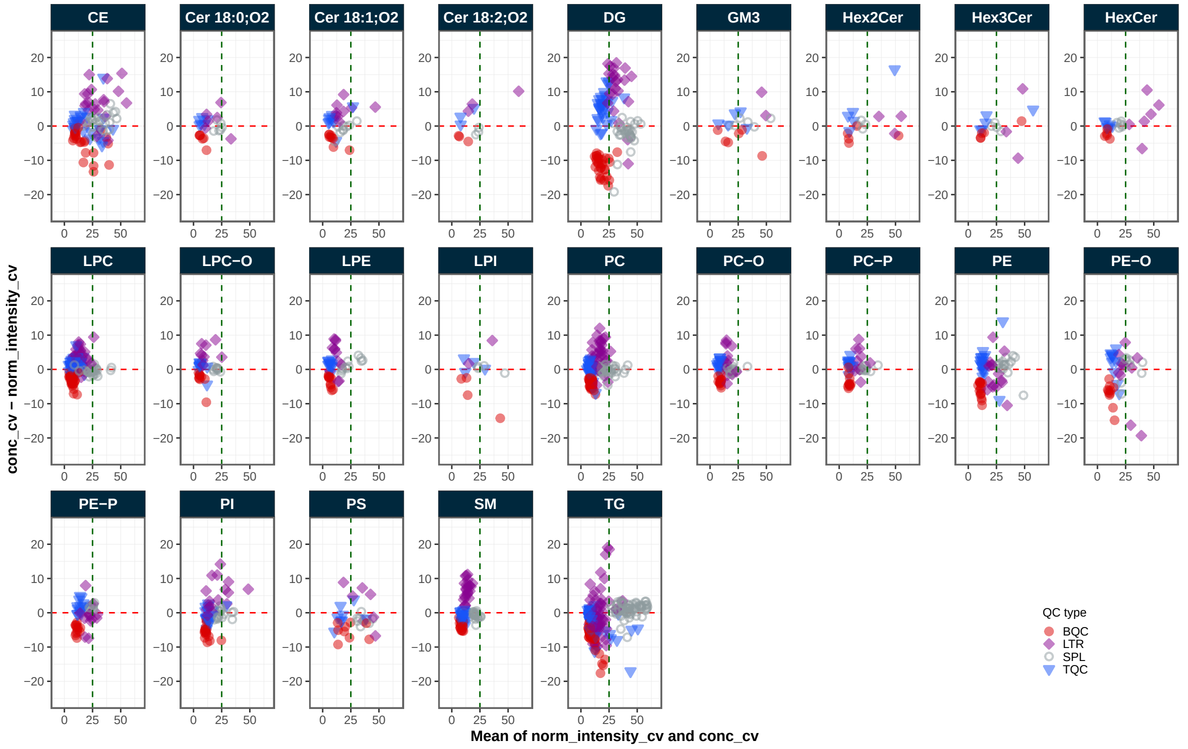

#### 3.15 18. Technical variability versus intensity

Comparing technical variability of features with their abundances can reveal the intensity threshold at which feature signals exhibit increased noise and variability. This information can be applied during peak picking to set appropriate intensity thresholds for feature detection and to reduce time spent on integrating low abundant features.

```
mexp <-  calc_qc_metrics(mexp, use_robust_cv = FALSE,use_batch_medians = TRUE)

fig5e <- plot_qcmetrics_comparison(
```

```
  mexp,
  plot_type = "scatter",
  y_shared = FALSE,
  "intensity_median_bqc",
  "intensity_cv_bqc",
  log_scale = FALSE,
  equality_line = FALSE,
  facet_by_class = FALSE,
   point_size = 1.1,
  font_base_size = 6,
  #x_lim = c(0, Inf),
  y_lim = c(0, 100)
)+
ggplot2::geom_smooth(method = "loess", se = FALSE, span = 0.75)+
geom_hline(yintercept = 20, linetype = "dashed", color = "grey70") +
scale_x_log10(expand = ggplot2::expansion(mult = c(0, 0.00)),
             breaks = c(1E2, 1E3, 1E4, 1E5, 1E6, 1E7, 1E8)) +

scale_y_continuous(expand = ggplot2::expansion(mult = c(0.03, 0.00)),breaks = c(0, 20, 40, 6

    theme(
      strip.text = ggplot2::element_text(size = 6),
      #aspect.ratio = 0.9,
      legend.position = "none",
      #panel.grid.major = element_line(linewidth = 0.3),
      legend.position.inside = c(0.9, 0.10),
      axis.text = element_text(size = 6),
      #axis.title = element_blank(),
      legend.direction = "vertical",
      legend.text = element_text(size = 6*0.7),
      legend.title = element_text(size = 6*0.7),
      legend.key.size = unit(6 *0.7, "pt"))
#> Scale for x is already present.
#> Adding another scale for x, which will replace the existing scale.
#> Scale for y is already present.
#> Adding another scale for y, which will replace the existing scale.
fig5e
#> `geom_smooth()` using formula = 'y ~ x'
```

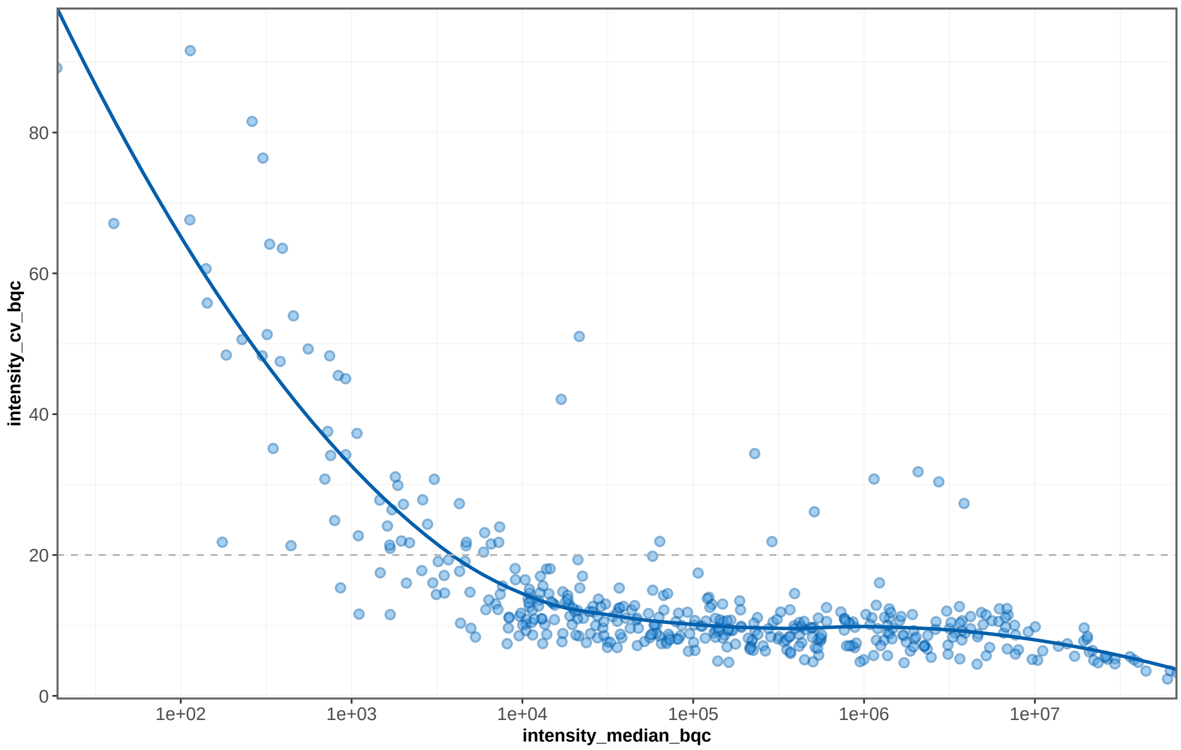

#### 3.16 19. BQC vs TQC variability

Comparison of the variability in BQCs vs TQCs may be useful to undertand the source of variability in the dataset. If the variability in BQCs is much higher than in TQCs, this may suggest that the sample processing is a major contributor to overall variability. If the variability in TQCs and BQCs is similar, this may suggest that technical variability is a major contributor to overall variability. We first look at two sphingomyelins (SM), phosphatidylcholines (PC), and triglycerides (TG). PC and TG species consistently show higher variability in BQCs than in TQCs, suggesting that sample processing, i.e., lipid extraction was a major contributor to overall variability, whereby it is unclear what factor(s) in the sample processing contributed to this variability.

SMs on the other hand show a similar variability in BQCs and TQCs, suggesting that the sample processing lesss contributed to the overall variability. This may be due to the fact that SMs are generally more abundant and thus less affected by variability in extraction efficiency.

```
mexp_temp <-  calc_qc_metrics(mexp, use_robust_cv = TRUE,use_batch_medians = FALSE)

plot_qcmetrics_comparison(
  mexp_temp,
```

```r
  plot_type = "diff",
  y_shared = FALSE,
  "norm_intensity_cv_tqc",
  "norm_intensity_cv_bqc",
  log_scale = FALSE,
  equality_line = TRUE,
  facet_by_class = TRUE,cols_page = 8,
    point_size = 1.3,
  font_base_size = 5,
  x_lim = c(0, 25),
  y_lim = c(-6, 10)) +
    theme(
      strip.text = ggplot2::element_text(size = 8),
      #aspect.ratio = 0.9,
      legend.position = "inside",
      legend.position.inside = c(0.9, 0.10),
      axis.text = element_text(size = 6),
      #axis.title = element_blank(),
      legend.direction = "vertical",
      legend.text = element_text(size = 9*0.7),
      legend.title = element_text(size = 9*0.7),
      legend.key.size = unit(6 *0.7, "pt"))
```

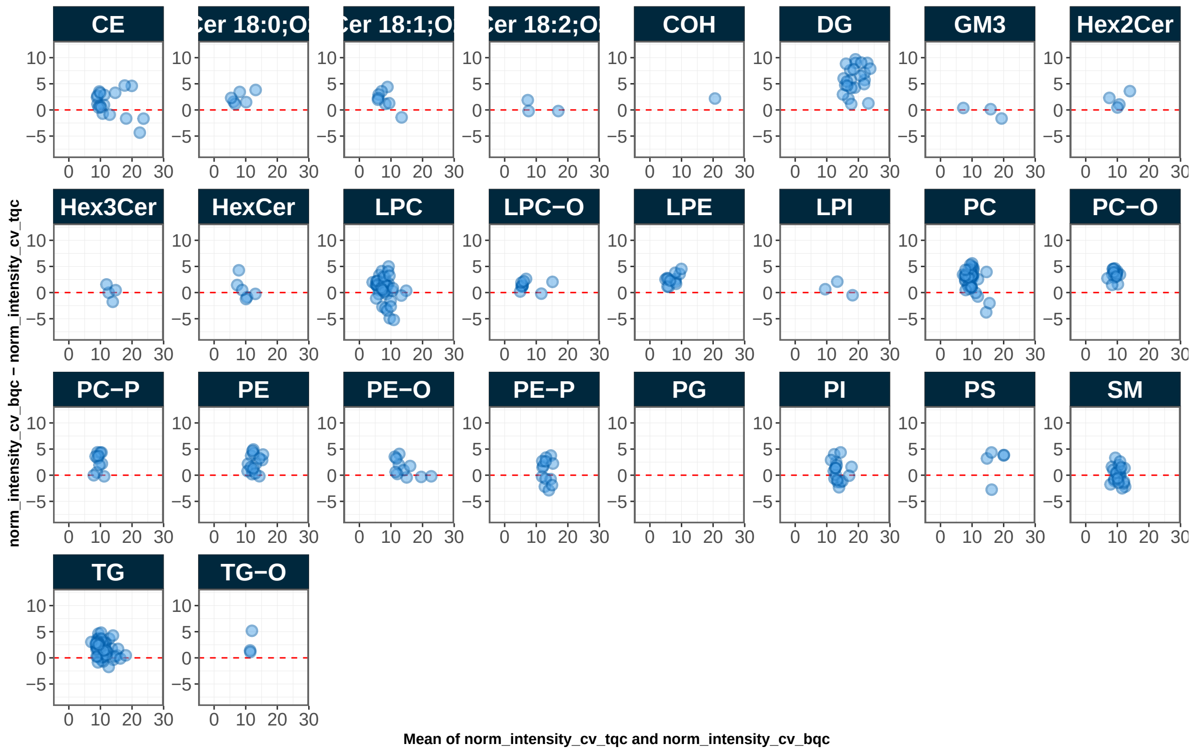

#### 3.17 20. Response curves

```
sel_species <- c("PC 33:1 d7 (ISTD)", "PC 32:1", "PC 34:2", "PE 32:1")
mexp_temp <- mexp
mexp_temp@dataset <- mexp_temp@dataset %>%
  filter(feature_id %in% sel_species) %>%
  mutate(feature_id = factor(feature_id, levels = sel_species)) %>%
  arrange(feature_id)

mexp_temp@annot_responsecurves <- mexp_temp@annot_responsecurves |> filter(curve_id %in% c("

fig5f <- plot_responsecurves(
  data = mexp_temp,
  variable = "intensity",
  filter_data = FALSE,font_base_size = 4, line_width = 0.5, point_size = 1.2,
  include_feature_filter = sel_species,
  output_pdf = FALSE, path = "response-curves.pdf",
  show_progress = FALSE,
  cols_page = 2, rows_page = 2,return_plots = T)[[1]]+
```

```
  theme(
    panel.grid.minor = element_blank(),
    legend.position = "inside",
    legend.direction = "horizontal",
    legend.text = element_text(size = base_font_size*0.6),
    legend.title = element_blank(),
    legend.key.size = unit(base_font_size *0.6, "pt"),
    strip.text = element_text(size = 6),
    legend.position.inside = c(0.9, 0.01))
#> Registered S3 methods overwritten by 'ggpp':
#>   method                  from
#>   heightDetails.titleGrob ggplot2
#>   widthDetails.titleGrob  ggplot2
#> Generating plots (1 page)...  done!
fig5f
```

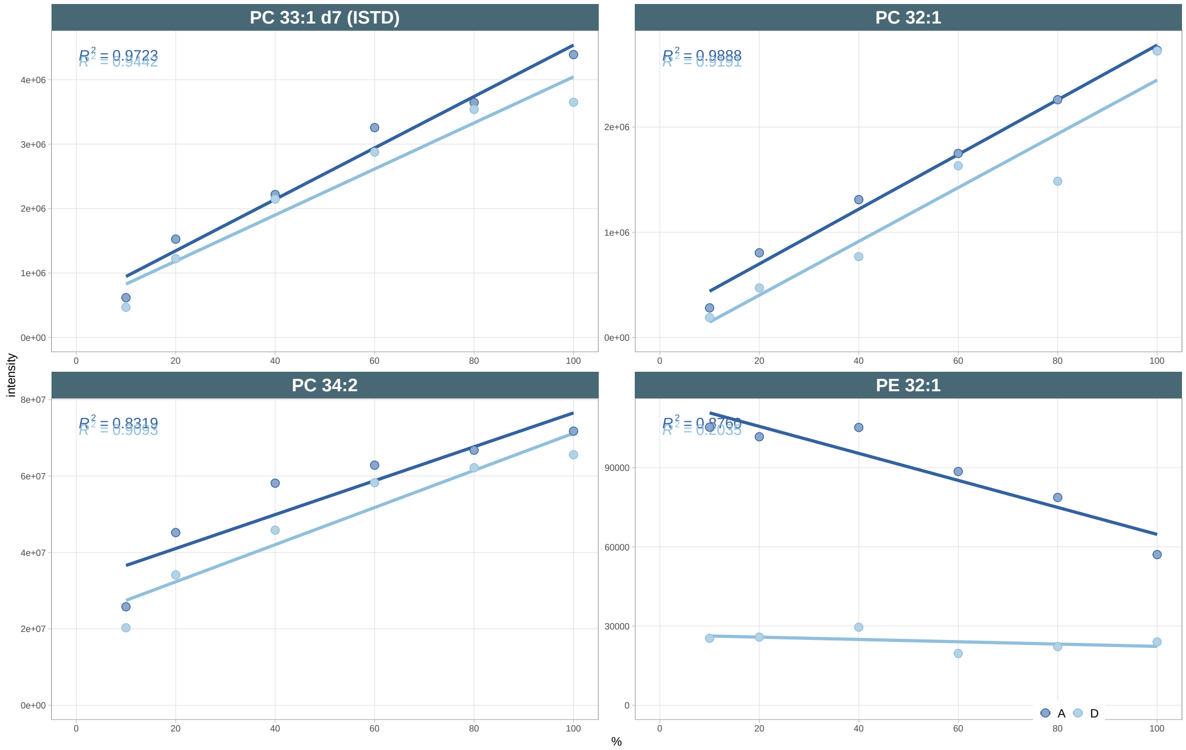

##### 3.18 21. Feature filter

```
mexp_final  <- filter_features_qc(
  data =  mexp,
  recalc_metrics = TRUE,
  clear_existing = TRUE,
  use_batch_medians = TRUE,
  include_qualifier = FALSE,
  include_istd = FALSE,
  response.curves.selection = c(1,2),
  response.curves.summary = "mean",
  min.rsquare.response = 0.8,
  min.slope.response = 0.5,
  max.yintercept.response = 0.5,
  min.signalblank.median.spl.pblk = 10,
  min.intensity.median.spl = 100,
  max.cv.conc.bqc = 25,
  max.dratio.sd.conc.bqc = 0.75,
  max.prop.missing.conc.spl = 100,
  features.to.keep = c("CE 20:4", "CE 22:5", "CE 22:6", "CE 16:0", "CE 18:0")
)
#> Calculating feature QC metrics - please wait...
#> ! The following features were forced to be retained despite not meeting filtering criteri
#> v
New feature QC filters were defined: 316 of 412 quantifier features meet QC criteria (not in
```

##### 3.19 22. Feature filter Results

```
lipidCat_order <- c("Cer 18:0;O2", "Cer 18:1;O2", "Cer 18:2;O2",  "SM",  "Hex2Cer", "Hex3Cer
                    "LPC", "LPE", "LPI", "LPC-O", "PC", "PE", "PI", "PG", "PS",
                    "PC-O", "PC-P", "PE-O", "PE-P", "DG", "TG", "TG-O", "COH", "CE")

mexp_final@metrics_qc$feature_class <-
  factor(mexp_final@metrics_qc$feature_class, lipidCat_order)
mexp_final@metrics_qc <- mexp_final@metrics_qc |> arrange(feature_class)

fig5h <- plot_qc_summary_byclass(mexp_final) +
  theme(
    strip.text = ggplot2::element_text(size = 5),
```

```
    #aspect.ratio = 0.9,
    legend.position = "inside",
    axis.text = element_text(size = 4),
    axis.title = element_text(size = 6, face = "plain"),
    axis.text.y.right =  element_text(size = 4, face = "plain"),
    legend.direction = "vertical",
    legend.text = element_text(size = 6*0.7),
    legend.title = element_blank(),
    legend.key.size = unit(6 *0.7, "pt"),
    legend.position.inside = c(0.77, 0.27))
fig5h
```

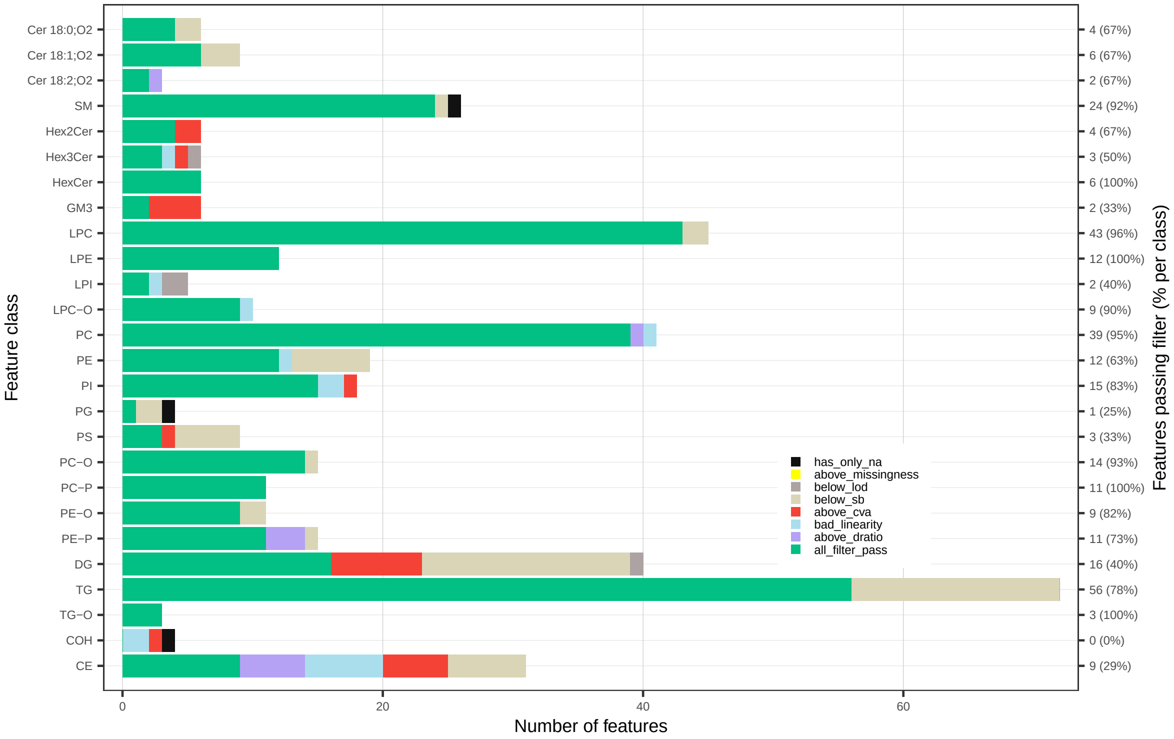

#### 3.20 23. Overview analytical variability

The plot below provides an overview of the analytical variability of the filtered dataset, showing the distribution of coefficients of variation (CVs) in the BQCs for all features. We aslo see that the cholesteryl esters (CE), which we included in the final dataset despite failing QC, show a higher variability. In subsequent statistical analyses and interpretation, it will be important to consider that these CE species for forced to be included in the dataset.

```
plot_abundanceprofile(
  data = mexp_final,
  use_qc_metrics = TRUE,
  log_scale = FALSE,
  filter_data = TRUE,
  variable = "conc_cv_bqc",
  density_strip = TRUE,
  qc_types = "SPL",
  #analysis_range = c(1,4000),
  show_sum = FALSE,
  #x_lim = c(6.5, 7.5),
  x_label = NA,
  feature_map = "lipidomics")
```

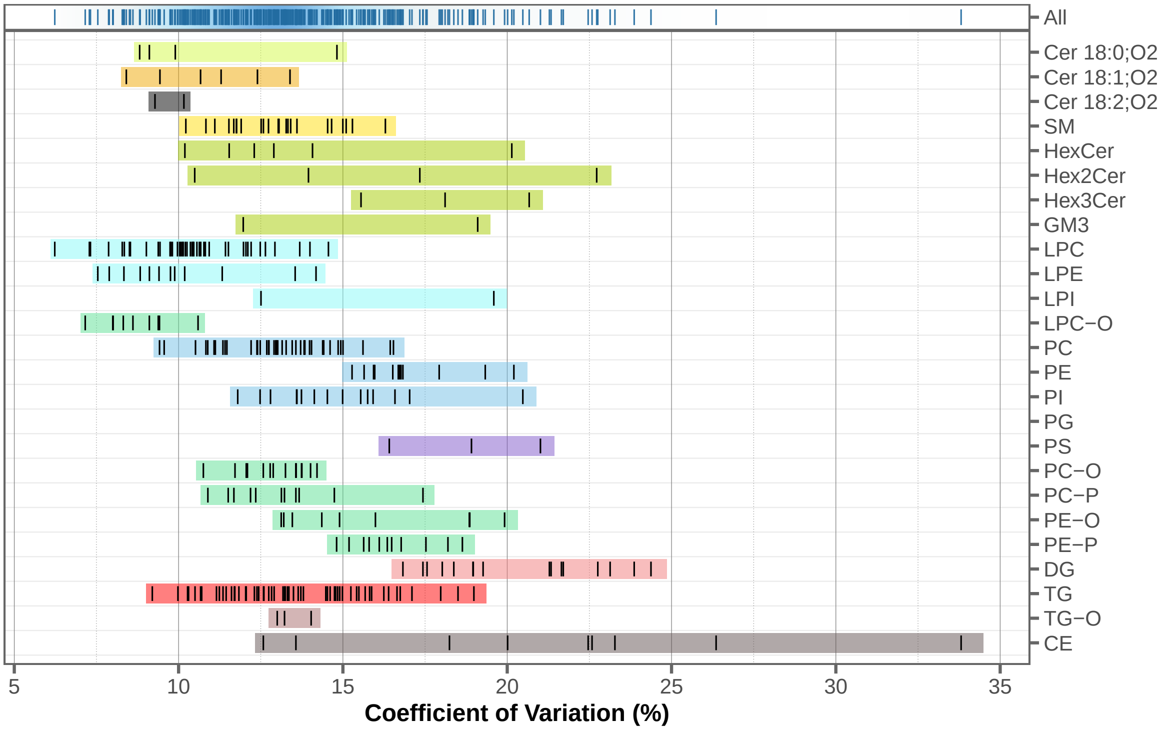

#### 3.21 24. Feature filter Venn

```
fig5ex <- plot_qc_summary_overall(mexp_final)
```

```
fig5g <- fig5ex[[2]] +
  theme(
    strip.text = ggplot2::element_text(size = 5),
    #aspect.ratio = 0.9,
    legend.position = "inside",
    axis.text = element_text(size = 4),
    axis.title = element_text(size = 6),
    legend.direction = "vertical",
    legend.text = element_text(size = 6*0.7),
    legend.title = element_text(size = 6*0.7),
    legend.key.size = unit(6 *0.7, "pt"))

fig5g
```

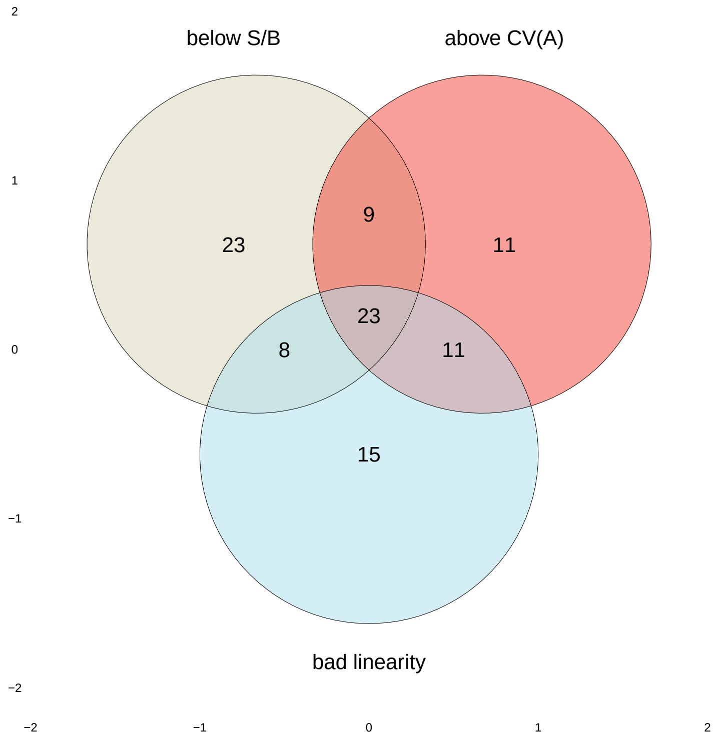

##### 3.22 25. Lipidome Profile

As a final overview, the feature concentration profile of the filtered dataset is shown below. Validating the concentrations, for example, those of the most abundant species, the summed concentrations per lipid class, or the ratios between lipid classes, against in-house reference values or literature data helps ensure that the quantification is within expected ranges and that no major quantification errors are present.

```
fig5i <- plot_abundanceprofile(
  data = mexp_final,
  log_scale = TRUE,
  variable = "conc",
  filter_data = TRUE,
  qc_types = "SPL",
  #x_lim = c(-6, 2),
  x_label = NA,font_base_size = 6,
  feature_map = "lipidomics")

fig5i
```

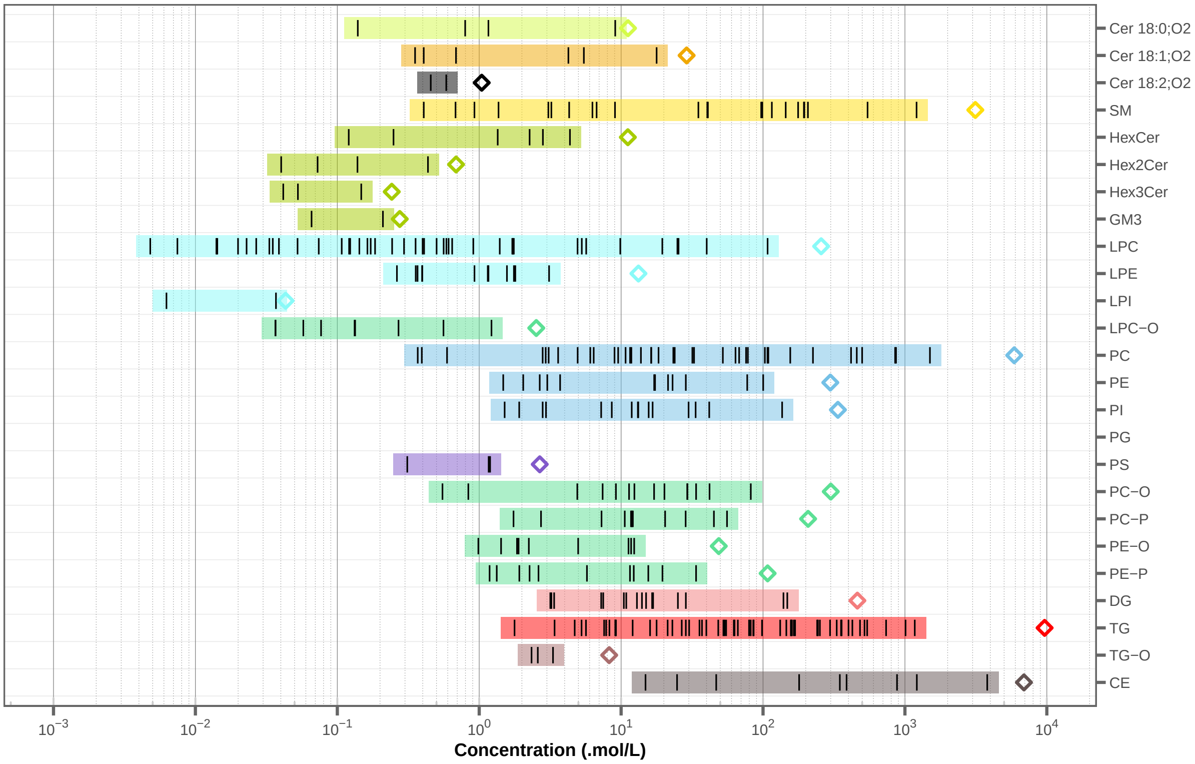

#### 3.23 26. Exporting final dataset

```
save_dataset_csv(
  data = mexp,
```

```
  path = "./output/sperfect_UNFILTERED_RAW-feature_conc_uM_20250518a.csv",
  variable = "conc_raw",
  qc_types = "SPL",
  include_qualifier = FALSE,
  filter_data = FALSE)
#> v Conc_raw values for 678 analyses and 412 features have been exported to './output/sperf
```

### 4 Figures for the manuscript in preparation

**Note**: the code and corresponding plots below are used for a manuscript currently in preparation**.

#### 4.1 1. Assembling Figure 4

```
fig4 <- (
  (fig4a | fig4b) /
  ((fig4c | fig4d | fig4e) + plot_layout(widths = c(0.9, 0.9, 1))) /
  ((fig4f | fig4g) + plot_layout(widths = c(1.07, 0.93)))
) +
  plot_layout(heights = c(1, 0.9, 1)) +
  plot_annotation(tag_levels = 'a') +
  theme(
    plot.tag = element_text(size = 12, face = "bold"),
    strip.placement = 'inside'
  )

ggsave("output/fig4.png", fig4, width = 180, height = 200, units = "mm", dpi = 300)
ggsave("output/fig4.pdf", fig4, width = 180, height = 200, units = "mm", dpi = 300)
```

#### 4.2 2. Assembling Figure 5

```
layout <- "
AABB
CCDD
EEFF
"
```

```r
fig5 <- (fig5a |fig5b) /
  #(p_fig2c | p_fig2d) /
  (fig5c | fig5d) /
  ((fig5e | fig5f | (fig5g + theme(axis.text = element_blank(), axis.title = element_blank()
    plot_layout(widths = c(1.0,1.1, 0.9))) /
  (fig5h | fig5i) +
plot_layout(heights = c(0.8,1,0.9,1.5)) +
plot_annotation(tag_levels = 'a') +
  theme(plot.tag = element_text(size = 10, face = "bold"),
        strip.placement = 'inside')
fig5
#> `geom_smooth()` using formula = 'y ~ x'
```

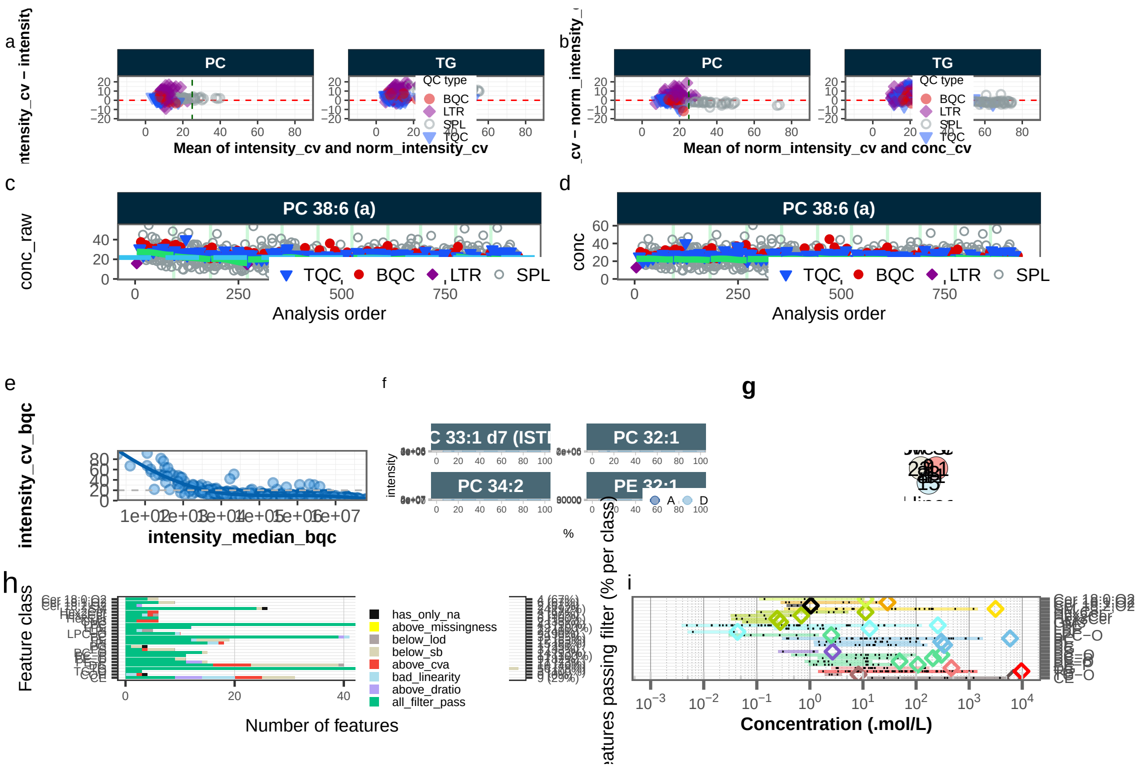

```r
ggsave("output/fig5.png", fig5, width = 180, height = 240, units = "mm", dpi = 300)
#> `geom_smooth()` using formula = 'y ~ x'
ggsave("output/fig5.pdf", fig5, width = 180, height = 240, units = "mm", dpi = 300)
#> `geom_smooth()` using formula = 'y ~ x'
#> Warning in grid.Call.graphics(C_text, as.graphicsAnnot(x$label), x$x, x$y, :
#> conversion failure on 'Concentration ( mol/L)' in 'mbcsToSbcs': for   (U+03BC)
```

#### 4.3 3. Assembling Extended Figure 4

```
# include fig4b again as extfig2a for better comparison of RLAs before and after normalizati
extfig2a <- fig4b

extfig2 <- (extfig2a / extfig2b / extfig2c / extfig2d) +  plot_layout(ncol = 1, heights = c(
  plot_annotation(tag_levels = 'a') +
  theme(plot.tag = element_text(size = 19, face = "bold"),
        strip.placement = 'inside')

ggsave(extfig2, filename = "output/ExtendedFig-4_RLAs.pdf",
  width = 180, height = 250, units = "mm", dpi = 300)
extfig2
```

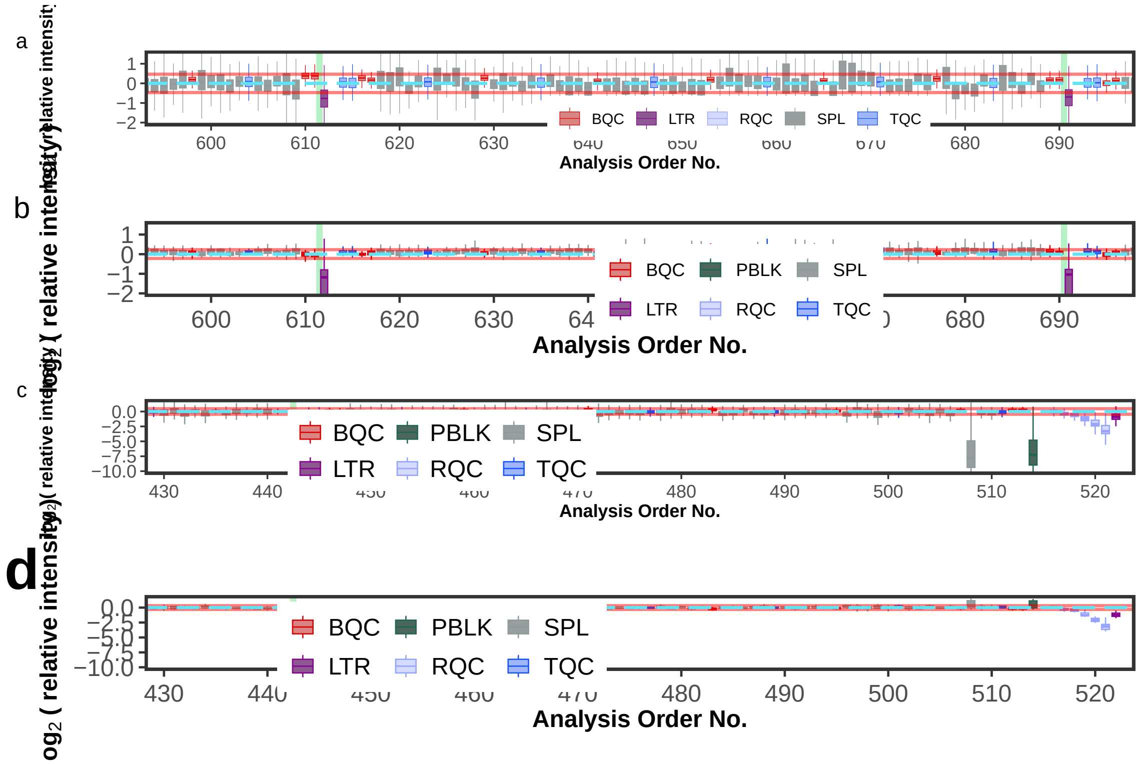

#### 4.4 3. Assembling Extended Figure 5

```
extfig3 <- extfig3a / extfig3b +
  plot_annotation(tag_levels = 'a')+
```

```
  theme(plot.tag = element_text(size = 12, face = "bold"),
        strip.placement = 'inside')

ggsave(extfig3, filename = "output/ExtendedFig-5.png",
  width = 180, height = 250, units = "mm", dpi = 300)
extfig3
```

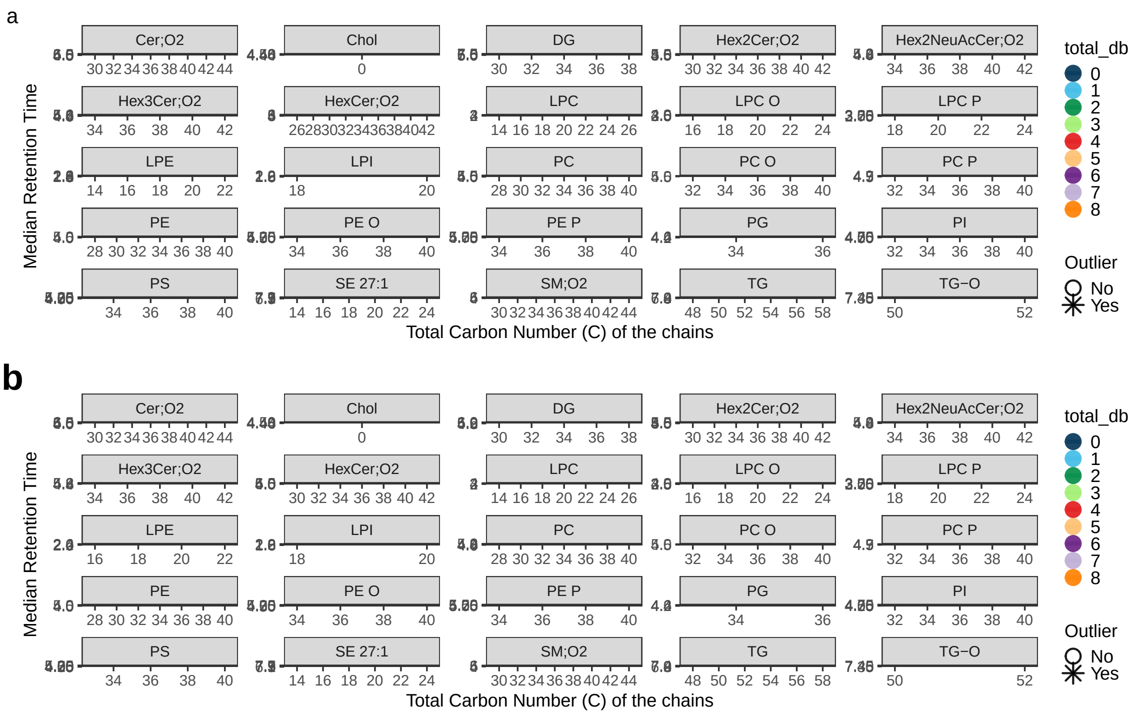

#### 4.5 4. Assembling Extended Figure 6

```
extfig4 <- extfig4a / extfig4b + plot_layout(ncol = 1, heights = c(1,1.5)) +
  plot_annotation(tag_levels = 'a') +
  theme(plot.tag = element_text(size = 12, face = "bold"),
        strip.placement = 'inside')

ggsave(
  extfig4, filename = "output/ExtendedFig-6_PCAloading.png",
  width = 180, height = 250, units = "mm", dpi = 300)
ggsave(
```

```
  extfig4, filename = "output/ExtendedFig-6_PCAloading.pdf",
  width = 180, height = 250, units = "mm", dpi = 300)
extfig4
```

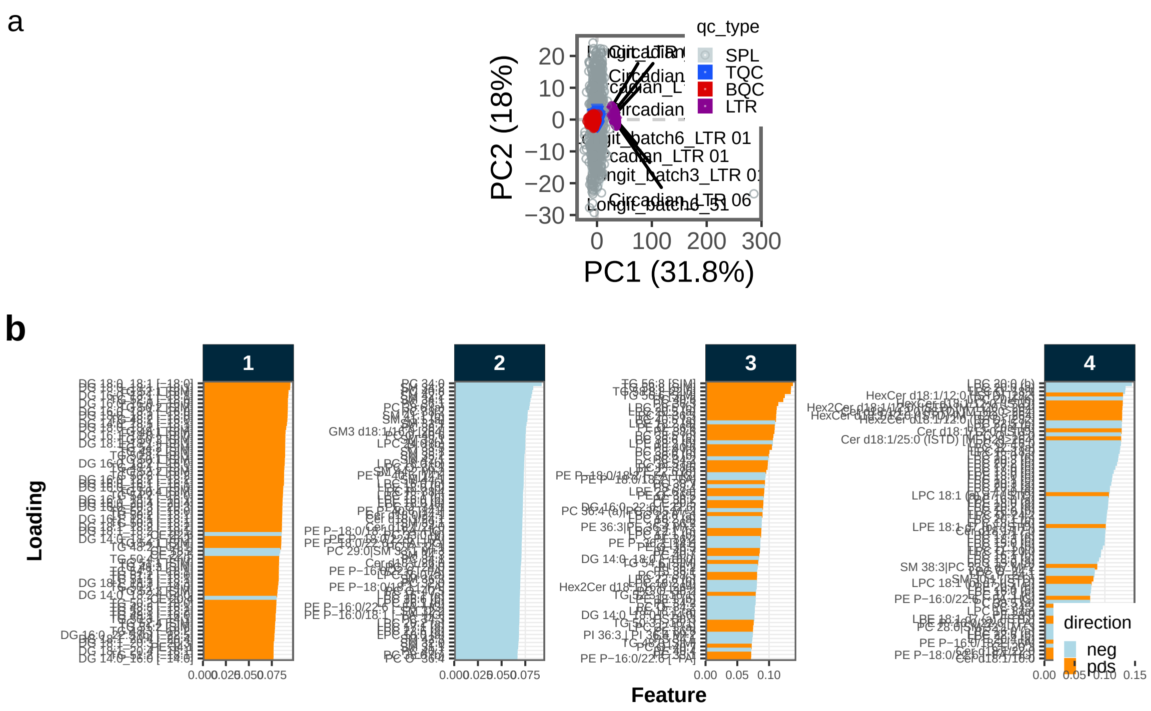

##### 4.6 5. Saving Extended Figure 7

```
ggsave(extfig5[[1]], filename = "output/ExtendedData_Fig7_ISTDtrends.png",
  width = 180, height = 180, units = "mm", dpi = 300)
```

##### 4.7 5. Assembling Extended Figure 8

```
extfig6 <- extfig6a / extfig6b / extfig6c + plot_layout(ncol = 1, heights = c(1,1,1)) +
  plot_annotation(tag_levels = 'a') +
  theme(plot.tag = element_text(size = 12, face = "bold"),
        strip.placement = 'inside')
ggsave(extfig6, filename = "output/ExtendedData_Fig8_CVnormbeforeafter.pdf",
```

```
  width = 160, height = 270, units = "mm", dpi = 600)
extfig6
```

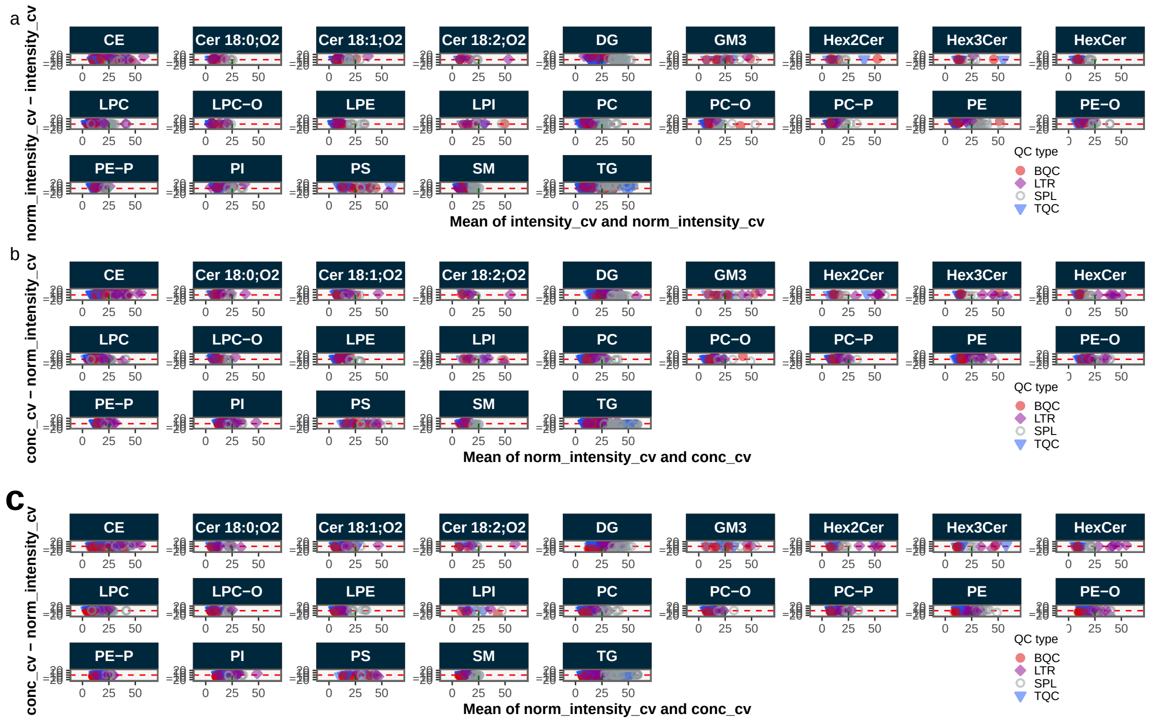

### 5 Supplementary Figures of Manuscript

#### 5.1 Runscatter plots

```
plot_runscatter(mexp,
  variable = "intensity",
  qc_types = c("SPL", "BQC", "TQC", "LTR"),
  show_reference_lines = TRUE,
  ref_qc_types = "SPL",
  reference_fill_color = "#111111",
  reference_k_sd = 3,
  show_trend = FALSE,
  point_size = 1.2,
  base_font_size = 6,
```

```r
  cols_page = 3,
  rows_page = 3,
  cap_outliers = TRUE,
  reference_sd_shade = FALSE,
  #batch_zebra_stripe = TRUE,
  show_progress = FALSE,
  output_pdf = TRUE,
  path = "output/suppl-fig1_runscatter_area.pdf", return_plot = FALSE)

plot_runscatter(mexp,
  variable = "conc_raw",
  qc_types = c("SPL", "BQC", "TQC", "LTR"),
  show_reference_lines = TRUE,
  ref_qc_types = "SPL",
  reference_fill_color = "#111111",
  reference_k_sd = 3,
  show_trend = TRUE,
  point_size = 1.2,
  base_font_size = 6,
  cols_page = 3,
  rows_page = 3,
  cap_outliers = TRUE,
  reference_sd_shade = FALSE,
  #batch_zebra_stripe = TRUE,
  show_progress = FALSE,
  output_pdf = TRUE,
  path = "output/suppl-fig2_runscatter_conc-raw.pdf", return_plot = FALSE)

plot_runscatter(mexp,
  variable = "conc",
  qc_types = c("SPL", "BQC", "TQC", "LTR"),
  show_reference_lines = TRUE,
  ref_qc_types = "SPL",
  reference_fill_color = "#111111",
  reference_k_sd = 3,
  show_trend = TRUE,
  point_size = 1.2,
  base_font_size = 6,
  cols_page = 3,
  rows_page = 3,
  cap_outliers = TRUE,
  reference_sd_shade = FALSE,
```

```
  #batch_zebra_stripe = TRUE,
  show_progress = FALSE,
  output_pdf = TRUE,
  path = "output/suppl-fig3_runscatter_conc-final.pdf", return_plot = FALSE)
```
