## Supplementary Code 2 for "MRMhub: one-stop solution for automated processing of large-scale targeted metabolomics data"

### MRMhub Data Processing Workflow for Dataset 3

**Supplementary Code 2**

Bo Burla Guo Shou Teo Hyungwon Choi

2025-12-09

**References** **50**

### 1 Overview Dataset

This example workflow uses a targeted lipidomics dataset measuring 829 features across 4,591 samples (Chen *et al.* (Chen et al. 2025)) and demonstrates the complete MRMhub data processing pipeline, including peak integration, quantitation, quality control, and reporting. This document covers many elements of the MRMhub workflow for demonstration purposes, some of which may not be applicable or necessary when processing other datasets.

> **!** Important
>
> All datasets used and generated within this workflow have been deposited in the **Zenodo** repository https://zenodo.org/records/15370293. Within this repository, the raw mass spectrometry files and the MRMhub-INTEGRATOR input files required for peak integration are available under the record "**Dataset 3**". The data and metadata files used and produced by the code in this Quarto notebook can be found under the"**mrmhub-workflows**" record. Please note that due to size limitations, the corresponding GitHub repository does not include any of these datasets.

### 2 Raw Data Processing: Peak Picking and Integration

This section describes key aspects of raw data processing workflow used for this dataset. For more a detailed description on the workflow, see Workflow for Dataset 1 and the INTEGRATOR Manual.

#### 2.1 Conversion of vendor raw files to mzML

The original raw data files (Agilent .d) were converted to mzML using msconvert from the ProteoWizard software (https://proteowizard.sourceforge.io) with the following settings: output format = mzML; binary encoding precision = 32 bit; write index = true.

#### 2.2 Preparation of the INTEGRATOR input files

The final input files used for this dataset are available from the Zenodo repository (see above). The following subsections summarize the key settings used for this example.

##### 2.2.1 Global Settings File (param.txt)

The parameter mz_tol, defining the maximum tolerance in m/z values for identifying transitions, was set to 0.06. The RT_tol parameter, defining the window around the expected RT (see Feature/Transition table below) in which features are searched, was set to 0.04 min. This relatively narrow RT window was chosen to reduce selection of incorrect peaks in complex chromatograms, as observed in this dataset. The global peak_width parameter was set to [0.15, 0.08, 0.08, 0.18], which matched the majority of the peaks. For specific features with broader or convoluted peaks, feature-specific peak widths were applied, overwriting the global settings (see Feature Table below). The maximum allowed RT shift correction was set to (-0.1, 0.4), and the maximum allowed sample-to-sample RT shift was 0.05 min. The full set of settings in param.txt is shown in the figure below.

```
# location of .mzML files
mzML_files= 'mzML*/MS*Batch[3456]*/*.mzML'

### table indicating file name, batch, sample type and reference sample
batch_info ='run_order_20251010.csv'

### Table with transition information:
transition_list=  'transition_list_20251010.csv'

# If "peak_width = [w, x, y, z]", the left integration bound will be between
peak_width = [0.15, 0.08, 0.08, 0.18]

# Number of threads to use for parallel processing
num_threads = 16

# m/z tolerance, maximum difference indicated m/z and m/z values in mzML fil
mz_tol = 0.06

### RT tolerance, maximum difference indicated RT and detected RT
RT_tol = 0.04

# if "RT_shift = [x, y]", the estimation will assume that RT shift is no mor
RT_shift = [-0.1, 0.4]

# this parameter restricts RT shift between adjacent samples
RT_shift_bound = 0.05
```

Figure 1: **Global settings file (param.txt).** Screenshot of the param.txt file used for this dataset, showing all configured parameters and their values.

##### 2.2.2 Sample Table

In the sample table (run_order_20251009.csv) the samples to be processed were listed by their .mzML file names. The first two BQC (batch quality control) samples were selected as retention time reference anchors. The sample type for each sample was defined in column C (sample_type) using the MRMhub QC type nomenclature. This is optional and INTEGRATOR distinguishes only between blanks (identified by the presence of the text `BLK`), and non-blanks.

| | A | B | C | D |
|---|---|---|---|---|
| 1 | sample | batch | type | reference |
| 4 | PBlank-03_MS-3.mzML | P-08 | PBLK | |
| 5 | TQC_056_MS-3.mzML | P-08 | TQC | |
| 6 | BQC-08_0_MS-3.mzML | P-08 | BQC | x |
| 7 | LTR-08_MS-3.mzML | P-08 | LTR | |
| 8 | NIST-08_MS-3.mzML | P-08 | NIST | |
| 9 | LV1017832528_MS-3.mzML | P-08 | SPL | |
| 10 | LV1017832529_MS-3.mzML | P-08 | SPL | |
| 11 | LV1017832530_MS-3.mzML | P-08 | SPL | |

Figure 2: Sample Table. Screenshot of a portion of the CSV file used for this dataset (run__order__20251010.csv).

##### 2.2.3 Feature/Transition (Feature) Table

The feature/transition table (transition__list__20251010.csv) defined the transitions to be extracted from the .mzML files by precursor and product m/z values. Each feature (peak or peak group) was specified by its expected retention time (RT, in minutes). For some transitions, multiple features were defined. The corresponding internal standard feature for each entry was included in the table, however, these internal standards were not used by INTEGRATOR but were exported in the integration results.

| | A | B | C | D | E | F | G | H | I | J |
|---|---|---|---|---|---|---|---|---|---|---|
| 1 | Compound Name | Transition Name | ISTD | Precursor Ion | Product Ion | RT (w.r.t reference samples) | uniform_width (y/n) | left integration bound (w.r.t reference samples) | right integration bound (w.r.t reference samples) | peak width |
| 516 | PE 38:3 (a) | PE 38:3 (a\b) | PE 34:0 (ISTD) | 770.6 | 629.6 | 8.47 | y | | 8.64 | |
| 517 | PE 38:4 | PE 38:4 | PE 34:0 (ISTD) | 768.6 | 627.5 | 8.03 | n | | | |
| 518 | PE 38:5 (a) | PE 38:5 (a\b\c) | PE 34:0 (ISTD) | 766.5 | 625.5 | 7.03 | y | | 7.14 | 0.2, 0.15, 0.1, 0.15 |
| 519 | PE 38:5 (b) | PE 38:5 (a\b\c) | PE 34:0 (ISTD) | 766.5 | 625.5 | 7.18 | n | 7.14 | 7.24 | |
| 520 | PE 38:5 (c) | PE 38:5 (a\b\c) | PE 34:0 (ISTD) | 766.5 | 625.5 | 7.32 | n | 7.24 | | 0.12, 0.15, 0.1, 0.25 |
| 521 | PE 38:6 | PE 38:6 | PE 34:0 (ISTD) | 764.5 | 623.5 | 6.7 | n | | | |

Figure 3: **Feature/Transition Table**. Screenshot of a portion of the CSV file (transition__list__20251010__Final.csv) used for this dataset. Feature-specific peak widths and fixed integration borders were set for three adjacent features to ensure correct integration.

Feature-specific 'peak__width' settings were applied for multiple features in this dataset when peaks were broader, exhibited increased tailing, or consisted of convoluted peaks that had to be co-integrated (see Figure 4 a-d). Fixed borders were assigned for features for which automatic integration failed or produced inconsistent results due to being poorly separated from adjacent peaks or having noisy, and consequently were applied when necessary (see Figure 4 e-h). The final version of the feature/transition table (transition__list__20251010.csv) is the result from several rounds of parameter optimization with INTEGRATOR.

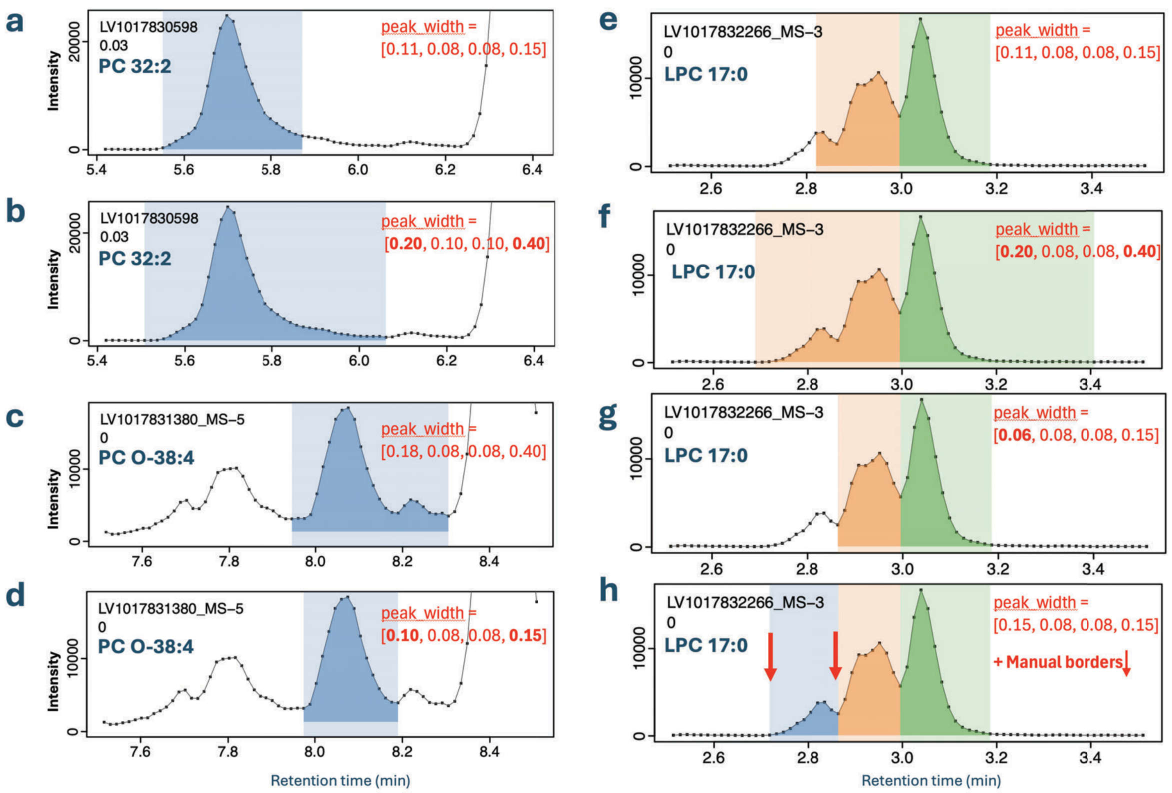

Figure 4: **Examples of manually set peak boundaries for MRMhub-INTEGRATOR.** (a, b) The effect of different peak_width parameter settings on the integration of a peak with clear tailing (PC 36:6). Panel (a) displays the result using the global setting, which is suitable for the majority of peaks. Panel (b) shows the outcome of applying a feature-specific peak width setting that overrides the global parameter for improved integration. (c, d) The influence of the peak_width parameter on the integration of a peak with nearby interferences (PC O-38:4). Panel (c) corresponds to the global setting, whereas panel (d) illustrates how an excessively wide peak border setting can cause incorrect integration, emphasizing the importance of establishing a suitable global value. (e, f) The effect of various peak integration settings on a complex chromatogram containing overlapping peaks. The global setting, shown in Panel (e), leads to partial integration where the first peak is not correctly recognized. While adjusting the peak width to be too relaxed resulted in the inclusion of the entire interfering signal and a more narrow setting led to its exclusion, neither was optimal. The final configuration, shown in Panel (f), used fixed boundaries defined for the first peak, which allowed for the correct picking and integration of all three isomer peaks of LPC 17:0.

#### 2.3 Running MRMhub-INTEGRATOR and Review of Results

The most recent release OF INTEGRATOR was obtained from https://github.com/SLINGhub/MRMhub/releases. After preparing all input files, the mrmhub application is started. Steps 1 to 4 then run consecutively without further input to perform peak detection, peak picking, integration, and export of data and PDF results. The integration results are reported in the generated file 'long.csv', a long-format table that includes peak areas, actual retention time, peak width, and other metadata. This data file will be used for subsequent data postprocessing (see the following section). A wide-format table with peak areas is available as ‘quant_raw.csv’. PDFs of the integrated transitions are available in the by_* folders. See the INTEGRATOR Manual for more details.

The peak integration results were inspected for all features across samples using the generated PDFs (see above). In this dataset, RT shifts across the entire run of 4,591 samples were less than 0.04 min, except for ~ 20 samples at the start of a batch towards the end of this analysis, which had increased RT shifts.

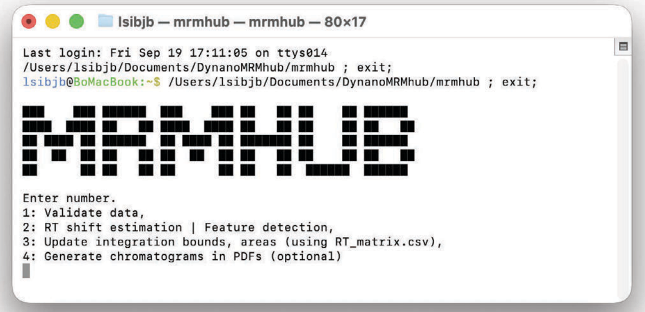

Figure 5: MRMhub-INTEGRATOR application.

### 3 Data Postprocessing and QC

#### 3.1 Setting up a Quarto Notebook and Installation of {mrmhub}

A Quarto notebook was created in RStudio the {mrmhub} R package was installed by running following code in the R console, which also installs two other R packages used in this workflow.

```
if (!require("remotes")) install.packages("remotes")
remotes::install_github("SLINGhub/MRMhub", subdir = "quant")
install.packages("here") # Used for parallel processing
```

The Quarto notebook code generated in this example workflow is available from https://github.com/SLINGhub/MRMhub-workflows (file Dataset3.qmd). In the subsequent section, the data postprocessing is encoded and documented step-by-step into the notebook as individual code chunks using {mrmhub} functions. Certain code chunks include additional R code to showcase specific examples for illustrative purposes. In real-world applications, this supplementary may not be necessary.

#### 3.2 Postprocessing workflow

##### 3.2.1 Load `mrmhub` and other required R packages

To improve performance multi-threading was used for some of the calculations and plottings, for which the R package {mirai} needs to be installed and loaded.

```
library(mrmhub)
library(mirai)  # Used for parallel processing
```

##### 3.2.2 Import the MRMhub-INTEGRATOR results

The results from the peak integration performed with the INTEGRATOR workflow, as described in the previous section, were imported into a `MRMhubExperiment` data object. This object represents the central data container used in this postprocessing workflow. Nogte that the original result file from INTEGRATOR (‘long.csv’) has been renamed to ‘Dataset3_MRMhub-Integrator_20251010.csv’

```
data_path <- "./data/dataset-3/data/Dataset3_MRMhub-Integrator_20251010.csv"
mexp <- MRMhubExperiment()
mexp <- import_data_mrmhub(mexp, data_path, import_metadata = TRUE)
#> v Imported 4591 analyses with 828 features
#> i `feature_area` selected as default feature intensity. Modify with `set_intensity_var()`
#> v Analysis metadata associated with 4591 analyses.
#> v Feature metadata associated with 828 features.
```

##### 3.2.3 Import the analysis metadata

Detailed analysis metadata, describing the analytes, samples, features, internal standards (ISTDs), and response curves required for subsequent steps, were imported from the Excel workbook ‘Dataset3_Metadata_20251010.xlsm’. This workbook is available within the previously mentioned Zenodo archive (record Dataset 3).

```
file_path <- "./data/dataset-3/data/Dataset3_Metadata_20251010.xlsm"
mexp <- import_metadata_msorganiser(mexp, file_path, ignore_warnings = TRUE)
#> ! Metadata has following warnings and notifications:
#> ----------------------------------------------------------------------------------------------
#>   Type Table    Column     Issue                            Count
#> 1 W*   Features feature_id Feature(s) not in analysis data    31
#> 2 W*   Features feature_id Feature(s) without metadata         1
#> ----------------------------------------------------------------------------------------------
```

```
#> E = Error, W = Warning, W* = Supressed Warning, N = Note
#> ------------------------------------------------------------------------------------------
#> v Analysis metadata associated with 4591 analyses.
#> v Feature metadata associated with 827 features.
#> v Internal Standard metadata associated with 26 ISTDs.
#> v Response curve metadata associated with 36 annotated analyses.
```

##### 3.2.4 Analytical design and timeline

An overview of the analysis structure, detailing the types and sequence of QC samples analyzed is provided by the plot below. It also shows information on the start and end dates, total duration, and median run time for each sample. Setting `show_timestamp = TRUE` in the code below will plot the absolute dates as the x-axis, showing three short interruptions, and the last 4 batches were measured approximately one year later.

```
plot_runsequence(
  mexp,
  show_batches = TRUE,
  qc_types = c("SPL", "BQC", "TQC", "PBLK", "UBLK", "RQC", "SBLK", "LTR", "NIST"),
  batch_zebra_stripe = TRUE, base_font_size = 6,
  batch_fill_color = "#fffbdb",
  segment_linewidth = 0.25,
  show_timestamp = FALSE) +
  theme(plot.title = element_text(size = 5))
```

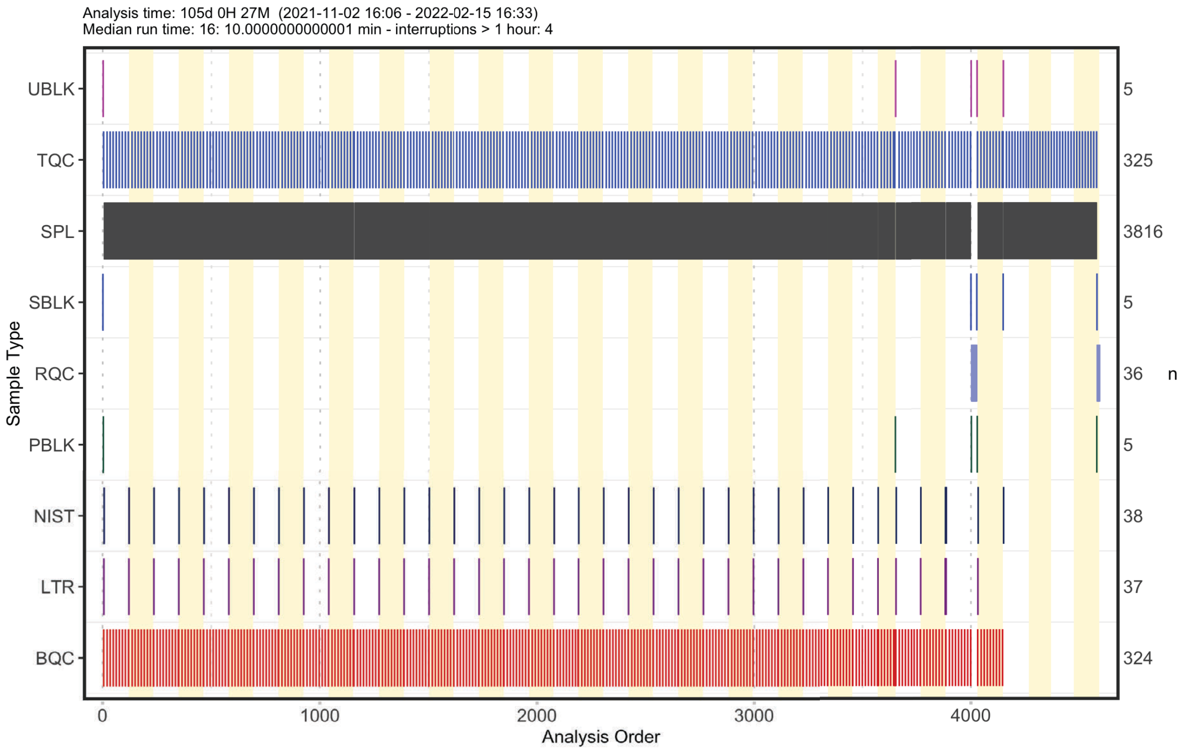

##### 3.2.5 Overview Chromatographic Separation

The following plot shows the retention time distribution of all detected lipid species. The ceramides (Cer) species show little chromatographic separation in contrast to other lipid classes, with the majority eluting between 10.0 to 10.5 min. The triglycerides (TG) also elute within a narrower retention time range at around 11 min.

```
plot_abundanceprofile(
  data = mexp,
  log_scale = FALSE,
  variable = "rt",
  density_strip = TRUE,
  qc_types = "SPL",
  analysis_range = c(1,4000),
  show_sum = FALSE,
  #x_lim = c(6.5, 7.5),
  x_label = NA,
  feature_map = "lipidomics")
```

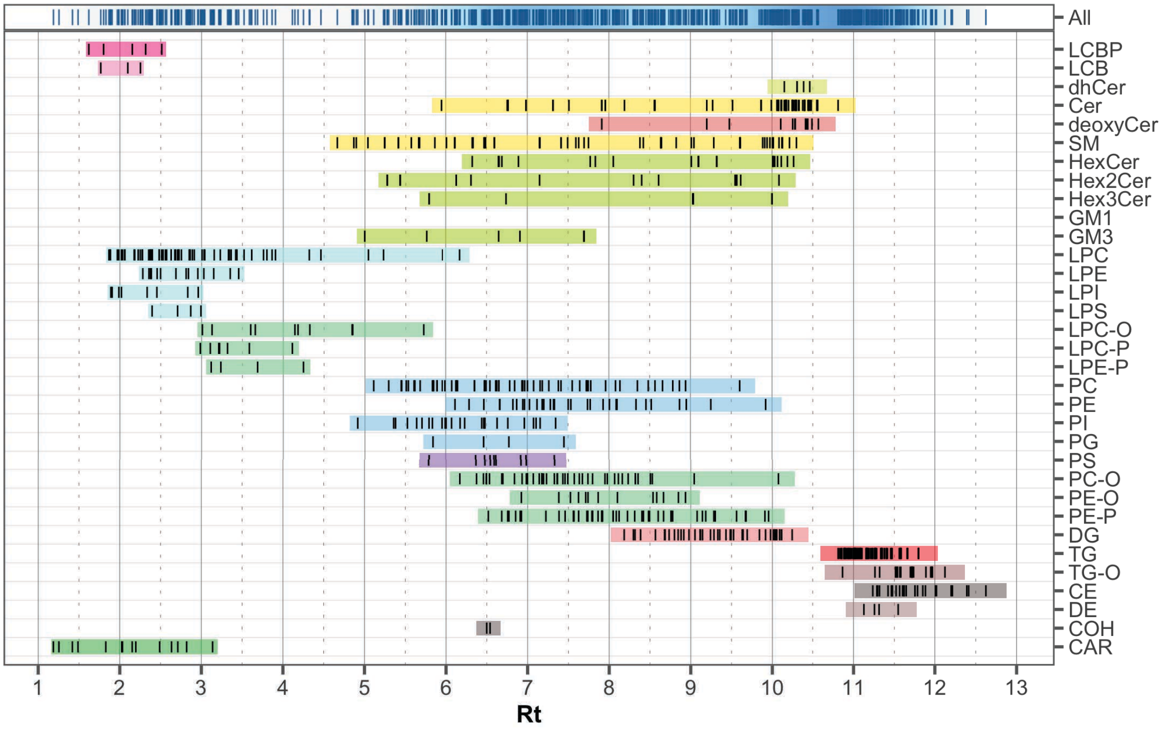

##### 3.2.6 Overall trends and check for possible outliers

To examine overall technical trends and issues affecting most analytes (features), the Relative Log Abundance (RLA) plot is useful (De Livera et al., 2015 (Livera et al. 2015)). In an RLA plot, each feature is normalized to the across-sample or within-batch median and the resulting values are shown as boxplots for each sample. This visualization helps identify technical problems such as pipetting errors, sample loss, or changes in injection volume or instrument sensitivity. First, an overview of all analyses (runs) is plotted.

First, an overview of all analyses (samples) or samples is plotted, whereby on the signals from the spiked-in internal standards signals (ISTDs) are shown. The red horizontal lines indicate 1.5X IQR outlier limits.

```
fig2a <- plot_rla_boxplot(
  data = mexp,
  rla_type_batch = c("within"),
  variable = "intensity",
  qc_types = c("BQC", "TQC", "SPL", "NIST", "LTR", "SBLK"),
  #plot_range = c(1, 4900),
  rla_limit_to_range = FALSE,
```

```
  filter_data = FALSE,
  min_feature_intensity = 1000,
  include_feature_filter = "ISTD",
  #y_lim = c(-4,4),
  show_timestamp = FALSE,
  outlier_method = "fold",
  outlier_k = c(-0.4, 0.3),
  outlier_exclude = FALSE,
  x_gridlines = FALSE,
  show_plot = TRUE,
  batch_zebra_stripe = TRUE,
  linewidth = 0.1
)
#> i Found 148 outliers in the 4545 shown analyses
```

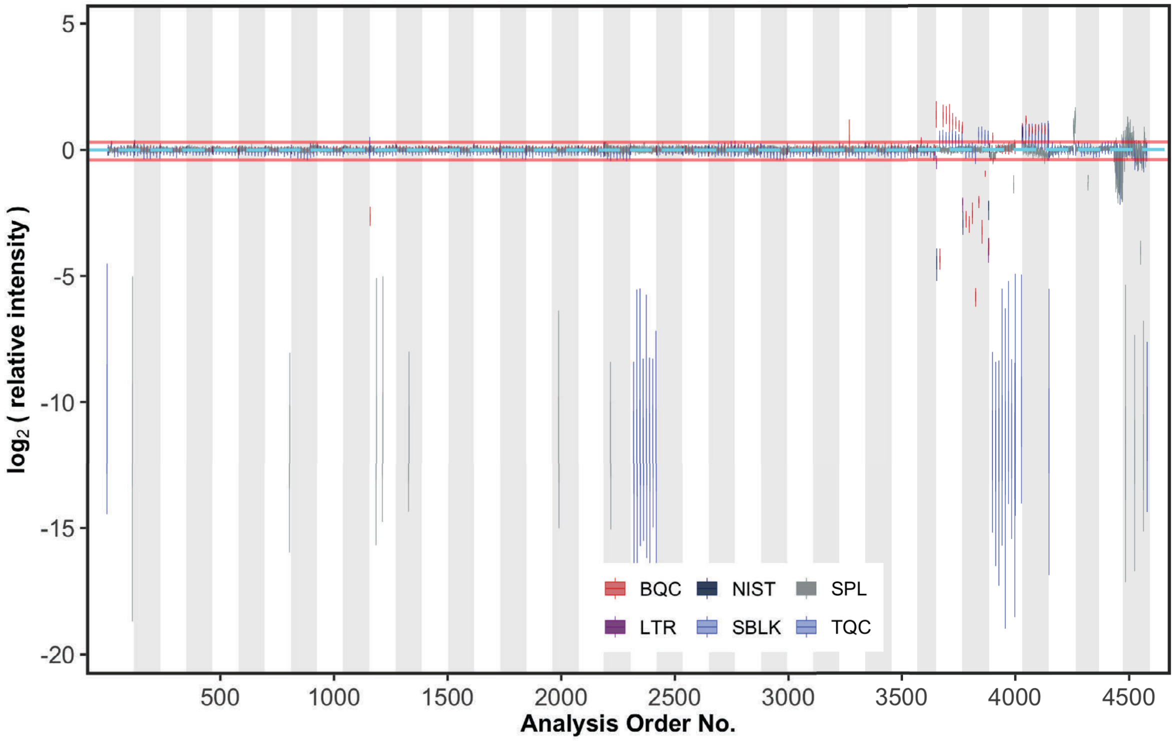

Several outliers among study and QC samples were detected using this RLA analysis (visible also in the plot above)

```
# Get outlier in ISTD total signal excluding 3 batches with overall lower ISTD
istd_outlier <- fig2a$outliers |>
```

```
  filter(!(batch_id %in% c("P-01", "P-02", "P-43") & val_res_median > -3))

print(istd_outlier)
#> # A tibble: 79 x 5
#>    analysis_id       analysis_order batch_id val_res_median qc_type
#>    <chr>                      <int> <chr>             <dbl> <chr>
#>  1 LV1017832619_MS-3            113 P-08             -11.8  SPL
#>  2 LV1017835020_MS-3            804 P-14             -11.8  SPL
#>  3 BQC-18_0_MS-4               1157 P-18              -2.69 BQC
#>  4 LV1017831110_MS-4           1185 P-18             -11.7  SPL
#>  5 LV1017832095_MS-4           1213 P-18             -12.1  SPL
#>  6 LV1017831999_MS-4           1328 P-19             -12.0  SPL
#>  7 LV1017834759_MS-4           1990 P-25             -11.9  SPL
#>  8 LV1017834663_MS-5           2218 P-27             -11.6  SPL
#>  9 TQC_217_MS-5                2320 P-28             -12.0  TQC
#> 10 TQC_218_MS-5                2334 P-28             -12.9  TQC
#> # i 69 more rows
```

To confirm these potential outliers, the peak areas of all ISTDs in different QC types is plotted against the run order (RunScatter plot). The outliers seen\ in these plots (with values close to zero) correspond to the outliers seen in the RLA plot above.

```
plot_runscatter(mexp, variable = "intensity",
                include_qualifier = FALSE,
                qc_types = c("SPL", "BQC", "TQC", "PBLK", "RQC", "SBLK"),
                include_feature_filter = "IS",
                #y_min = 0.00, y_max = 0.15,
                #plot_range = c(0, 910),
                point_size = .2,
                point_border_width = 0.1,
                point_transparency = .7,
                base_font_size = 4,
                cols_page = 3,
                rows_page = 11,
                show_progress = FALSE,
                cap_outliers = TRUE)
#> Generating plots (1 page)
```

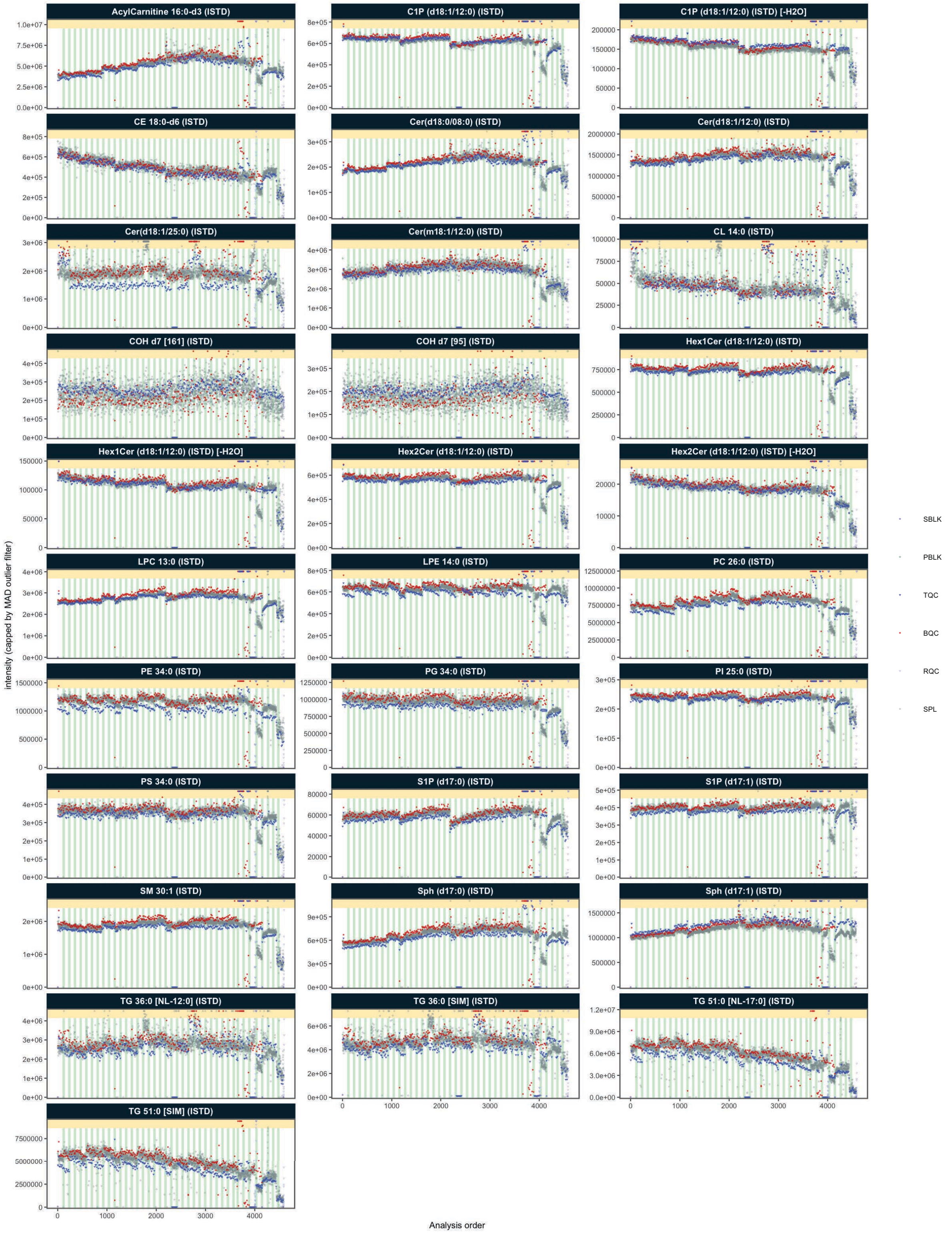
AcylCarnitine 16:0-d3 (ISTD)
C1P (d18:1/12:0) (ISTD)
C1P (d18:1/12:0) (ISTD) [-H2O]
CE 18:0-d6 (ISTD)
Cer(d18:0/08:0) (ISTD)
Cer(d18:1/12:0) (ISTD)
Cer(d18:1/25:0) (ISTD)
Cer(m18:1/12:0) (ISTD)
CL 14:0 (ISTD)
COH d7 [161] (ISTD)
COH d7 [95] (ISTD)
Hex1Cer (d18:1/12:0) (ISTD)
Hex1Cer (d18:1/12:0) (ISTD) [-H2O]
Hex2Cer (d18:1/12:0) (ISTD)
Hex2Cer (d18:1/12:0) (ISTD) [-H2O]
LPC 13:0 (ISTD)
LPE 14:0 (ISTD)
PC 26:0 (ISTD)
PE 34:0 (ISTD)
PG 34:0 (ISTD)
PI 25:0 (ISTD)
PS 34:0 (ISTD)
S1P (d17:0) (ISTD)
S1P (d17:1) (ISTD)
SM 30:1 (ISTD)
Sph (d17:0) (ISTD)
Sph (d17:1) (ISTD)
TG 36:0 [NL-12:0] (ISTD)
TG 36:0 [SIM] (ISTD)
TG 51:0 [NL-17:0] (ISTD)
TG 51:0 [SIM] (ISTD)
intensity (capped by MAD outlier filter)
Analysis order
SBLK
PBLK
TQC
BQC
RQC
SPL

Sepatedly, the full dataset is then checked for potential systematic outliers in endogenous analytes.

```
fig2b <- plot_rla_boxplot(
  data = mexp,
  rla_type_batch = c("within"),
  variable = "intensity",
  qc_types = c("BQC", "TQC", "SPL", "NIST", "LTR"),
  filter_data = FALSE,
  min_feature_intensity = 1000,
  exclude_feature_filter = "ISTD",
  plot_range = c(1, 4800),
  #y_lim = c(-4,4),
  show_timestamp = FALSE,
  outlier_method = "fold",
  outlier_k = c(-2,1) ,
  outlier_exclude = FALSE,
  x_gridlines = FALSE,
  batch_zebra_stripe = TRUE,
  show_plot = TRUE,
  linewidth = 0.1
)
#> i Found 81 outliers in the 4540 shown analyses
```

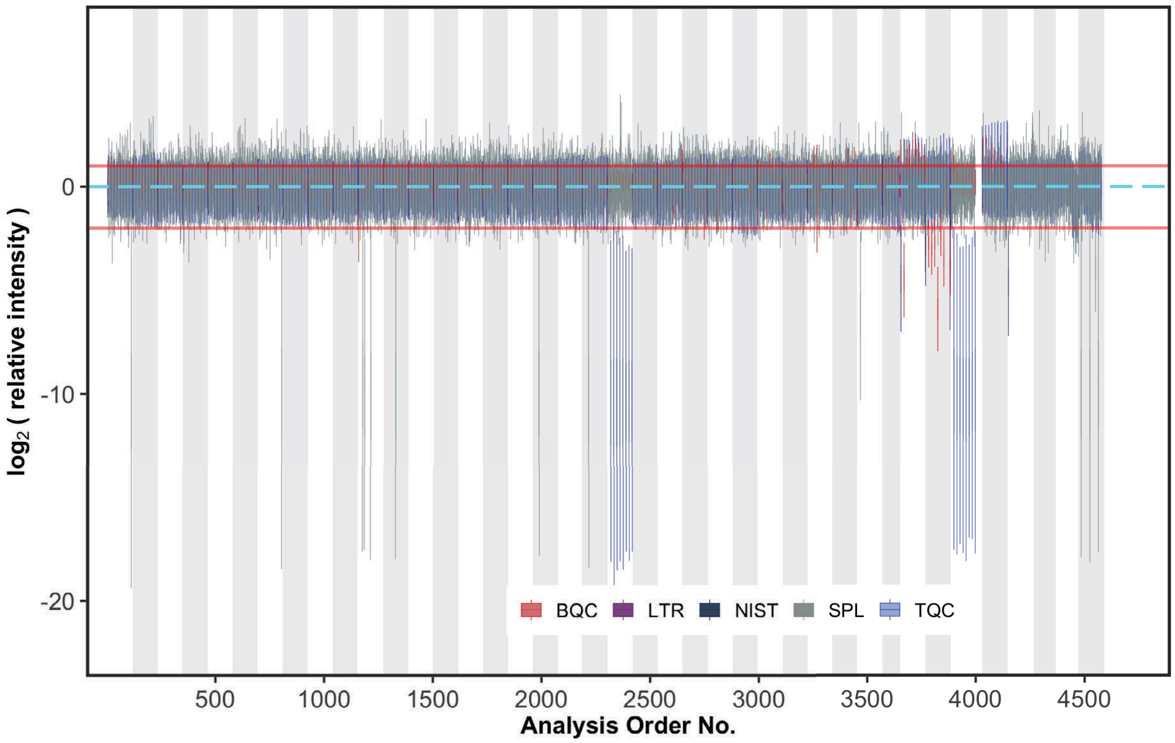

Also for the endogenous analytes, several outliers among study and QC samples were detected:

```
# Get outlier in ISTD total signal excluding 3 batches with conistently lower ISTD
analyte_outlier <- fig2b$outliers |>
  filter(!(batch_id %in% c("P-01", "P-02", "P-43") & val_res_median > -3))
print(analyte_outlier)
#> # A tibble: 72 x 5
#>    analysis_id       analysis_order batch_id val_res_median qc_type
#>    <chr>                      <int> <chr>             <dbl> <chr>
#>  1 LV1017832619_MS-3            113 P-08             -10.5  SPL
#>  2 LV1017835020_MS-3            804 P-14              -9.91 SPL
#>  3 BQC-18_0_MS-4               1157 P-18              -2.29 BQC
#>  4 LV1017832063_MS-4           1176 P-18              -9.48 SPL
#>  5 LV1017831110_MS-4           1185 P-18              -9.98 SPL
#>  6 LV1017832095_MS-4           1213 P-18             -10.0  SPL
#>  7 LV1017832134_MS-4           1260 P-18               1.21 SPL
#>  8 LV1017831999_MS-4           1328 P-19              -9.78 SPL
#>  9 LV1017834759_MS-4           1990 P-25              -9.82 SPL
#> 10 LV1017834663_MS-5           2218 P-27             -10.4  SPL
#> # i 62 more rows
```

Inspection of the generated plots and outlier tables identified several individual study samples, Technical QC (TQC), and Batch QC (BQC) samples that exhibited markedly lower signals for both internal standards (ISTDs) and endogenous analytes. TQC samples from two batches showed similarly reduced signals, while adjacent samples displayed normal signal levels, suggesting potential issues with sample preparation or errors in the LC-MS worklist for these specific samples and batches. Accordingly, these samples were classified as technical outliers and excluded from subsequent analysis.

Furthermore, in three batches analyzed toward the end of the sequence, all BQC samples exhibited significant differences in ISTD and analyte signals, whereas study samples in the same batch presented normal signals. This pattern is indicative of probable sample preparation errors affecting these BQCs. Consequently, they were also classified as technical outliers and excluded from the dataset..

Finally, an additional 18 samples were identified as outliers based on their endogenous analyte signals, although their ISTD signals remained comparable to other samples. These discrepancies were attributed to errors that occurred during the pipetting of sample aliquots into the extraction tubes.

```r
# Outliers in analyte but not ISTD signals
setdiff(
  analyte_outlier$analysis_id,
  istd_outlier$analysis_id
)
#>  [1] "LV1017832063_MS-4" "LV1017832134_MS-4" "LV1017834496_MS-5"
#>  [4] "LV1017834500_MS-5" "LV1017831025_MS-5" "BQC-30_8_MS-5"
#>  [7] "BQC-31_0_MS-5"     "LV1017831362_MS-5" "BQC-34_4_MS-5"
#> [10] "BQC-36_2_MS-5"     "BQC-37_5_MS-5"     "BQC-37_7_MS-5"
#> [13] "LV1017834075_MS-5" "LV1017832625_MS-5" "LV1017832682_MS-5"
#> [16] "NIST-04"           "LV1017830587"      "LV1017830728"
```

In total, 94 analyses/samples were classified as technical outliers and will be excluded from further analysis.

```r
# Combined ISTD and analyte outlier
outlier_combined <- istd_outlier |>
  bind_rows(analyte_outlier) |>
  bind_rows(tibble(analysis_id = "NIST-04", qc_type = "NIST")) |>
  select(-val_res_median) |>
  distinct()

outlier_combined |> dplyr::count(qc_type)
#> # A tibble: 5 x 2
```

```
#>   qc_type     n
#>   <chr>   <int>
#> 1 BQC        27
#> 2 LTR         3
#> 3 NIST        5
#> 4 SPL        35
#> 5 TQC        28

mexp <- exclude_analyses(mexp, analyses = outlier_combined$analysis_id, clear_existing = TRU
#> i 97 analyses were excluded for downstream processing. Please reprocess data.
```

To check if these identified outliers were removed the ISTDs are plotted again. The ISTD signals fairly stable in the first 35 batches, then varied in the subsequent last 4 batches.

```
plot_runscatter(mexp, variable = "intensity",
                include_qualifier = FALSE,
                qc_types = c("SPL", "BQC", "TQC", "PBLK", "RQC", "SBLK"),
                include_feature_filter = "IS",
                #y_min = 0.00, y_max = 0.15,
                #plot_range = c(0, 910),
                point_size = .2,
                point_border_width = 0.1,
                point_transparency = .7,
                base_font_size = 4,
                cols_page = 3,
                rows_page = 11,
                show_progress = FALSE,
                cap_outliers = TRUE)
#> Generating plots (1 page)
```

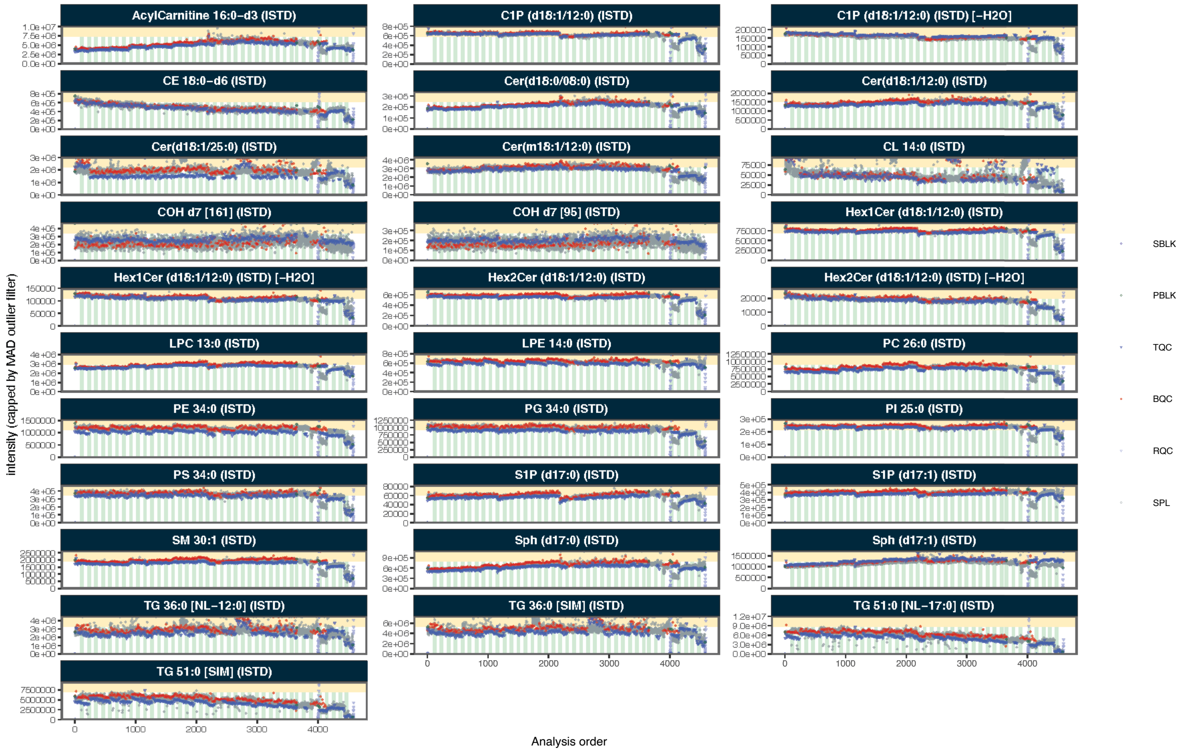

#### 3.3 Peak picking QC

To check for potential peak picking errors, the retention time (RT) of lipid features was plotted against the total carbon number and the number of double bonds in the acyl chains. This dataset was the result of an iterative process involving data review with QC plots and curation, which removed detectable annotation errors. Lipid species that remained flagged as potential misannotations were then manually inspected in the chromatograms and compared with an online resource providing peak annotation information for the utilized LC-MS method (https://metabolomics.baker.edu.au/method/lipids). Following this verification, these features were deemed likely to be correct.

```
plot_rt_vs_chain(
  mexp,
  qc_types = "SPL",
  x_var = "total_c", outlier_residual_min = 0.3,
  base_font_size = 6,
  cols_page = 4,
  point_size = 1) +
  theme(
    legend.position = "right",
```

```
    legend.direction = "vertical",
    legend.text = element_text(size = base_font_size*0.8),
    legend.title = element_text(size = base_font_size*0.8),
    legend.key.size = unit(base_font_size *0.8, "pt"),
    legend.position.inside = c(0.1, 0.7)
 )
#> ! The names of 8 of 827 feature_id(s) could not be parsed: PDMS-10 [+NH4], PDMS-11 [+NH4]
#> i The following features were flagged as potential annotation outliers: Cer(d18:0/08:0) (
```

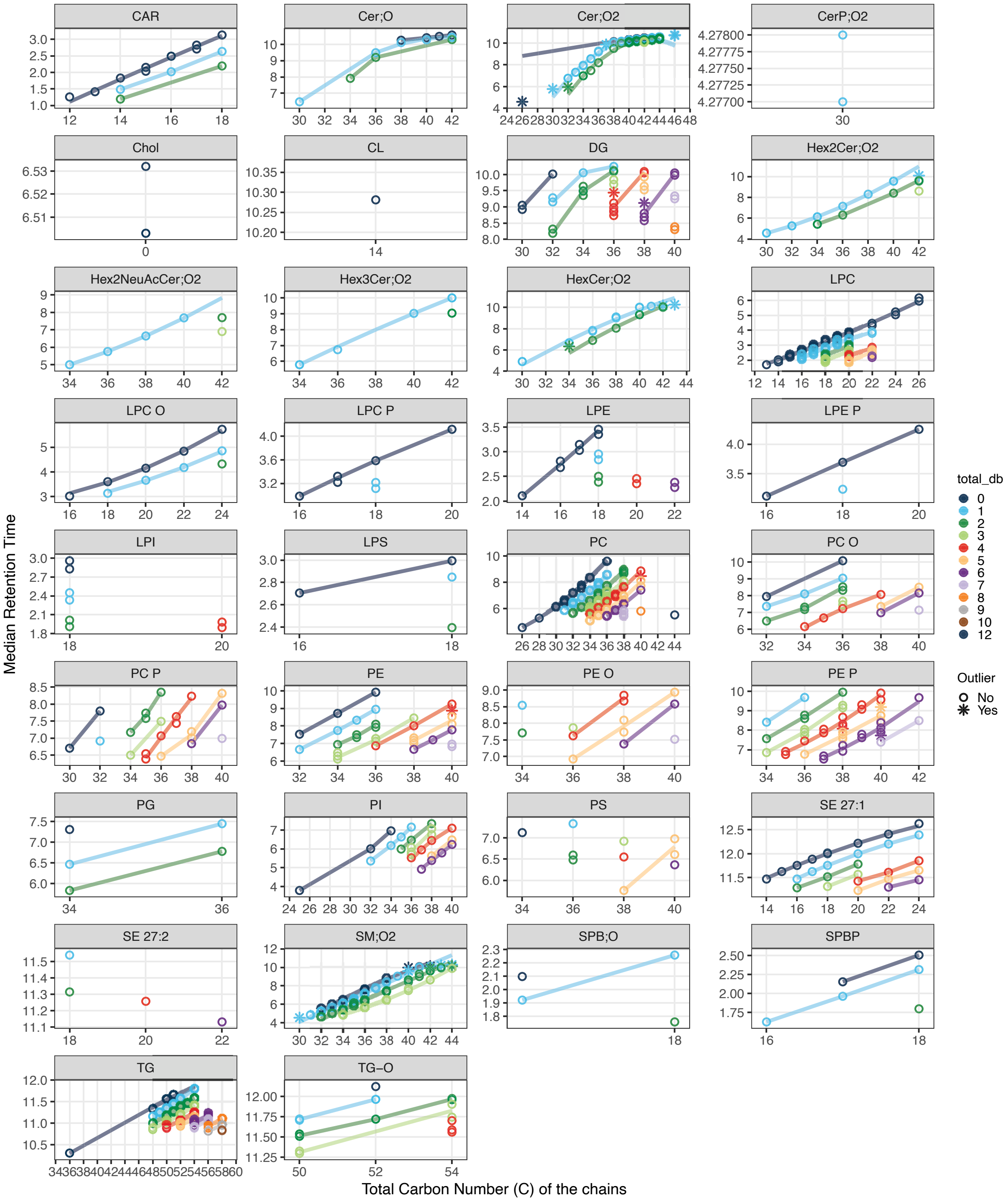
CAR
Cer;O
Cer;O2
CerP;O2
Chol
CL
DG
Hex2Cer;O2
Hex2NeuAcCer;O2
Hex3Cer;O2
HexCer;O2
LPC
LPC O
LPC P
LPE
LPE P
LPI
LPS
PC
PC O
PC P
PE
PE O
PE P
PG
PI
PS
SE 27:1
SE 27:2
SM;O2
SPB;O2
SPBP
TG
TG−O
Median Retention Time
Total Carbon Number (C) of the chains
total_db
0
1
2
3
4
5
6
7
8
9
10
12
Outlier
No
Yes

##### 3.4 Feature correlation analysis

Feature correlation analysis was performed as an additional check for potential peak annotation errors. In this analysis, feature pairs exhibiting high correlation (Pearson's r $>$ 0.98), based on their raw peak areas, were visualized using scatter plots.

```
# this below is to exclude a sample that has a very low intensity for all features, see next
# details
plot_feature_correlations(
  mexp,
  variable = "intensity" ,
  qc_types = c("SPL", "BQC", "TQC"),
  point_size = 0.5,
  cor_min = 0.98,
  point_stroke = 0.1,
  sort_by_corr = TRUE,
  return_plot = TRUE,
  show_progress = FALSE,
  log_scale = TRUE,
  cols_page = 5,
  rows_page = 6,
  font_base_size = 5)
#> Generating plots (1 page)
#> [[1]]
```

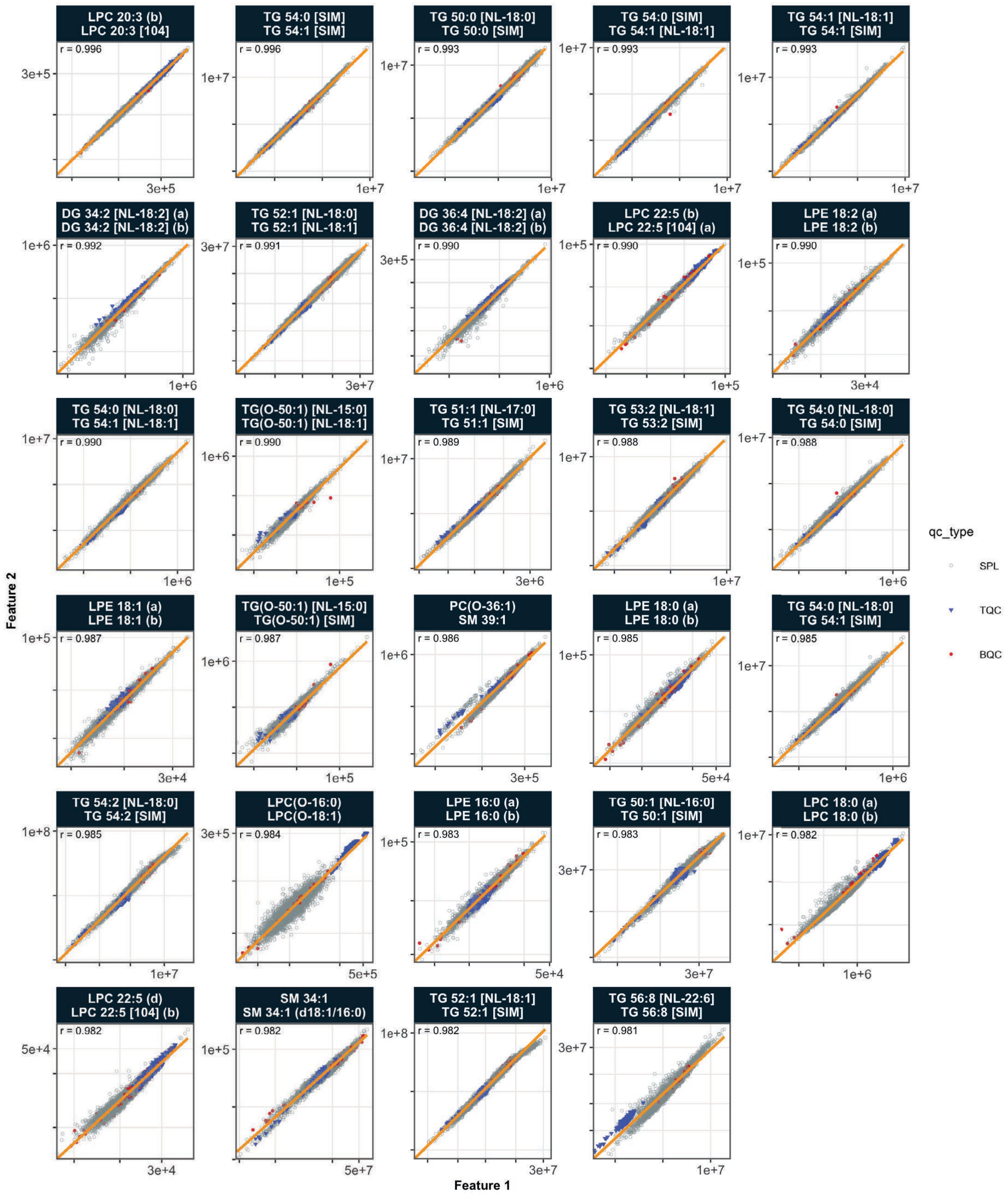

A high correlation (r = 0.986) was observed between features from unrelated classes, namely SM 39:1 and PC O-36:1. As such a strong correlation between unrelated species is highly improbable, this was presumed to be a likely technical or data annotation artefact. Investigation revealed that these features had overlapping retention times, and the M+1 isotope of SM 39:1 shared the same transition as PC O-36:1. A retention time analysis, confirmed with the previously mentioned online resource, indicated that PC O-36:1 should elute after SM 39:1. However, the peak corresponding to PC O-36:1 was found at the end of the MRM window and was partially truncated, which likely contributed to a peak picking error. Due to this truncation of the PC O-36:1 peak, this feature was excluded from subsequent analysis.

```
mexp <- exclude_features(mexp, features = "PC(O-36:1)", clear_existing = TRUE)
#> i 1 features were excluded for downstream processing. Please reprocess data.
```

##### 3.5 Summing up LysoPL and DG isomers

Highly correlating feature pairs from above analysis were found to consist of numerous lysophospholipid (lyso-PL) sn-1/sn-2 isomers and diacylglycerol (DG) isomer peaks. This is an expected observation, as spontaneous positional isomerization, or acyl migration, is known to occur in vitro during sample preparation and storage (Okudaira et al., J Lipid Research, 2014 (Okudaira et al. 2014)). Therefore, lyso-PL and DG isomer pairs were summed into a single feature before subsequent processing.

```
mexp <- data_sum_features(mexp, feature_classes = c("LPC", "LPE", "LPG", "LPI", "LPS", "DG"))
```

##### 3.6 PCA to check for potential technical outliers

To obtain an overview of the data and to perform an additional check for potential outlier samples, a Principal Component Analysis (PCA) was conducted based on the raw peak areas of all detected features. The results show that the BQC samples cluster within the study samples, whereas the TQC samples form a distinct group. Closer inspection reveals that TQCs from the last four batches are separated from the other TQCs. Many samples from these same batches are also found in the lower-left area of the plot, outside the bulk of the sample points. Furthermore, BQCs located outside of the main BQC cluster were observed, which are primarily from batches P-30 to P-38.

```
plot_pca(
  data = mexp,
  variable = "intensity",
  filter_data = FALSE,
  pca_dims = c(1,2),
  labels_threshold_mad = 5,
  labels_column = "analysis_order",
  qc_types = c("BQC", "TQC", "LTR", "NIST", "SPL"),
```

```
  ellipse_variable = "qc_type",
  log_transform = TRUE,shared_labeltext_hide = "_MS-5",
  point_size = 0.7, point_alpha = 0.7, font_base_size = 8, ellipse_alpha = 0.3,
  include_istd = FALSE,show_labels = TRUE,label_font_size = 1.5)   + theme(
    plot.title = element_blank(),
    aspect.ratio = 1,
        legend.position = "inside",
      legend.direction = "vertical",
      legend.text = element_text(size = 9*0.7),
      legend.title = element_text(size = 9*0.7),
      legend.key.size = unit(7 *0.7, "pt"),
      legend.position.inside = c(0.83, 0.83))
#> ! 66 features contained missing or non-numeric values and were exluded.
#> i The PCA was calculated based on `feature_intensity` values of 741 features.
```

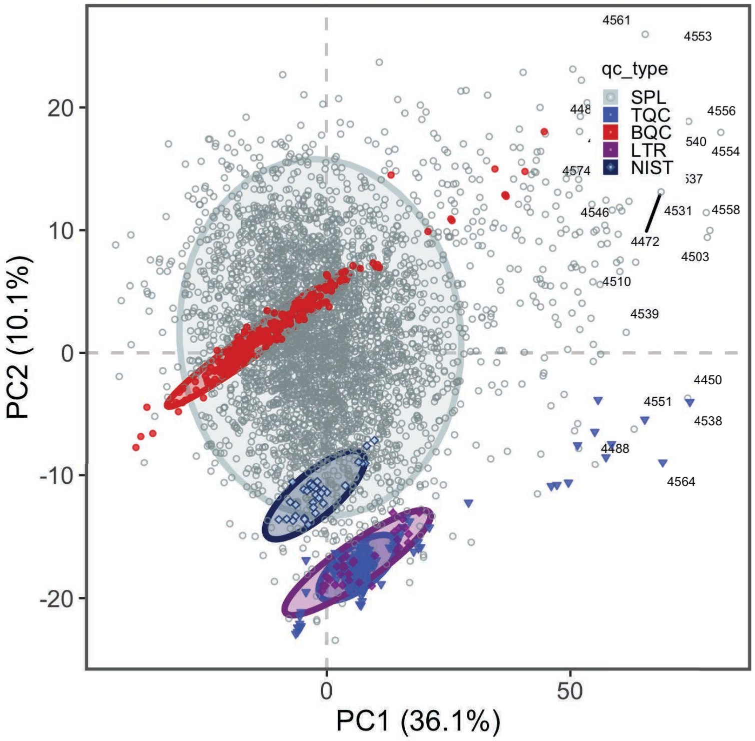

The loadings of the principal components (PC) revealed which features contribute to the variability in the data. While PC1 was contributed to by many features at similar levels, PC2 showed high loadings for numerous lyso-PC species. This may indicate systematic effects during sample preparation and/or analysis that specifically affected this class of lipids.

```
plot_pca_loading(
  data = mexp,
  variable = "feature_intensity",
  include_istd = FALSE,
  pca_dims = c(1,2,3,4),
  top_n = 70,
  font_base_size = 7,
  #qc_types = c("SPL", "BQC", "TQC", "LTR"),
  log_transform = TRUE,
  #point_size = 1, point_alpha = 0.7, font_base_size = 8, ellipse_alpha = 0.3,
  #include_istd = FALSE,
  #show_labels = TRUE,label_font_size = 2,
  #shared_labeltext_hide = NA
  )   + theme(
      plot.title = element_blank(),
      legend.position = "inside",
      legend.direction = "vertical",
      legend.text = element_text(size = 8*0.7),
      legend.title = element_text(size = 8*0.7),
      legend.key.size = unit(6 *0.7, "pt"),
      legend.position.inside = c(0.96, 0.06))
```

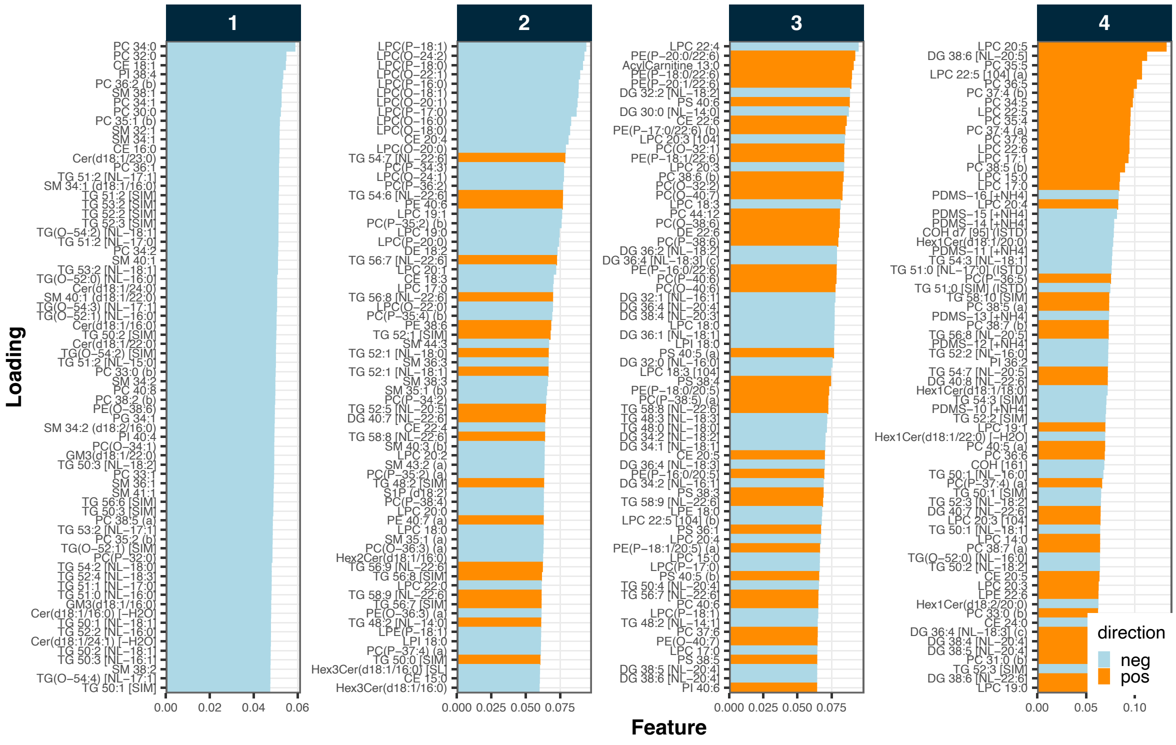

##### 3.7 Matrix effects

To assess potential matrix effects arising from differences between individual samples, the signal distributions of the internal standards (ISTDs) were plotted for each sample type. The signals correspond to the median of batch-wise normalized signals. Cholesterol and the triglyceride (TG) ISTDs exhibited considerably higher matrix effects relative to other ISTDs, as indicated by the wider distributions of their normalized intensities. On average, the Technical QC (TQC) samples displayed lower ISTD signals compared to the Batch QC (BQC) and study samples.

```
plot_qc_matrixeffects(
  mexp,
  variable = "intensity",
  batchwise_normalization = TRUE,
  only_istd = FALSE,include_qualifier = FALSE,
  include_feature_filter = "ISTD",
  exclude_feature_filter = "95|CL|25|d17\\:0|C1P|H2O",
  y_lim = c(50, 150),
  point_alpha = 0.05,
  box_alpha = 0.3,
  point_size = 0.2,
```

```
  box_linewidth = 0.2,
  font_base_size = 7,
  min_median_value = 1000)
```

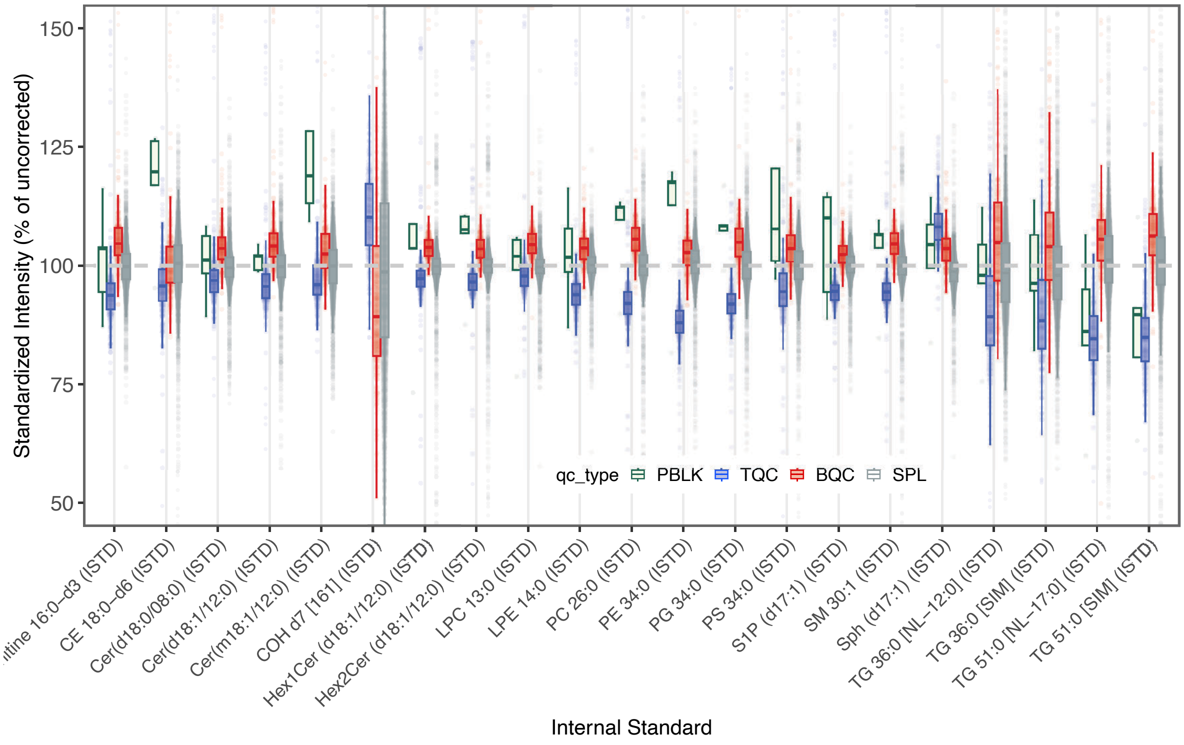

#### 3.8 Isotope correction

The measured, fully saturated phosphatidylcholine (PC) species, i.e., PC 26:0 (ISTD), PC 28:0, PC 30:0, PC 32:0, PC 34:0, and PC 36:0, coelute with the sphingomyelin (SM) species SM 30:1 (ISTD), SM 32:1, SM 34:1, SM 36:1, SM 38:1, and SM 40:1, respectively. Since the M+3 isotopes of these SM species share the same transition as the corresponding PC species, this can lead to an overestimation of PC concentrations. Therefore, an isotope correction was applied to these PC species based on the relative abundance of the SM M+3 isotopes. The correction factors, provided via the feature metadata, were derived using the method described by Gao et al., Analytical Chemistry, 2021 ((Gao et al. 2021))

The corresponding QC plot shows that relevant interference is present for PC 26:0 (ISTD), PC 28:0, and PC 30:0, and that PC 36:0 in particular had interference of up to 50%. In the case of PC 36:0, the interference contributed an average of 20% to the signal, ranging from 50% to negative values after correction. Instances where the correction was greater than the PC 36:0 peak area may be a result of analytical variability in both features or the presence of other interferences not accounted for in the SM 40:1 species. Any downstream analysis results for PC 36:0 must be interpreted with consideration of this substantial correction.

```
mexp <- correct_interferences(mexp)
#> ! Interference correction led to negative or zero values in 2 feature(s) in samples/QCs. I
#> v Interference-correction has been applied to 6 of the 766 features.
plot_qc_interferences(mexp, y_lim = c(-10, 110), qc_types = c("LTR", "NIST", "SPL", "TQC", "I
```

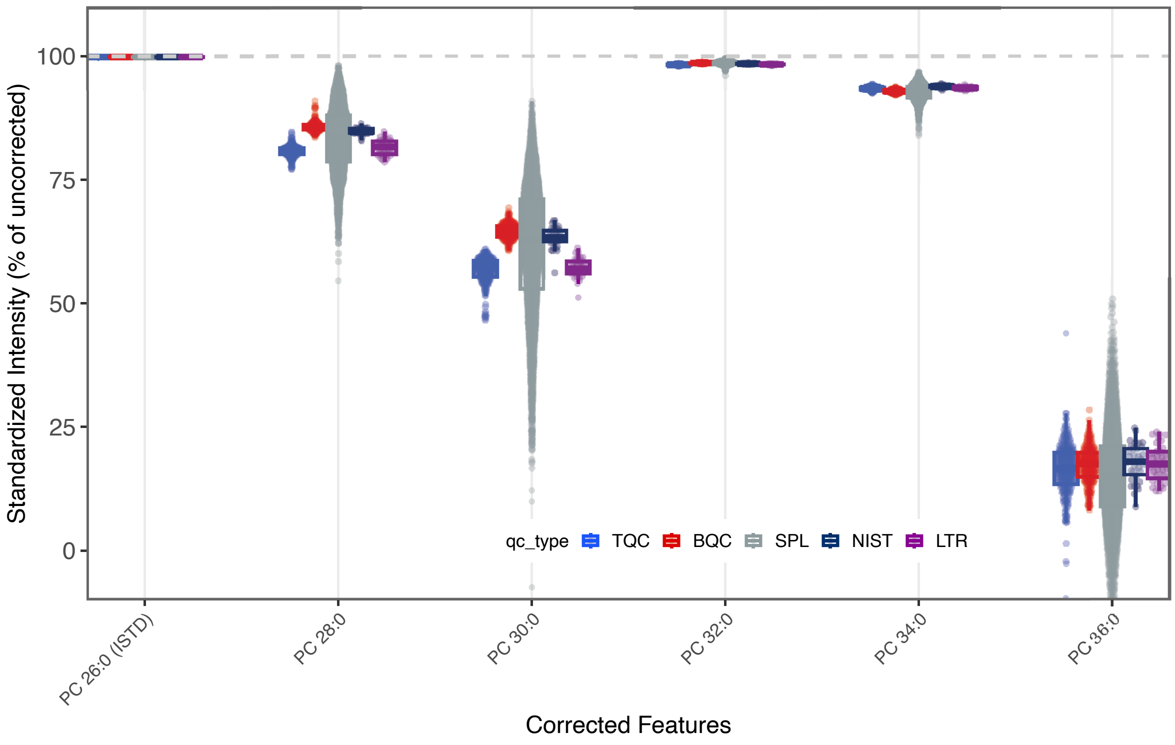

#### 3.9 Normalization and Quantification

The raw intensities were first normalized by their corresponding ISTD, as defined in the feature metadata. Concentrations were then calculated based on the spiked-in ISTD amounts and the sample amounts, which were also defined in the corresponding metadata.

```
mexp <- normalize_by_istd(mexp, ignore_missing_annotation = FALSE)
#> v 740 features normalized with 25 ISTDs in 4494 analyses.
mexp <- quantify_by_istd(mexp)
#> v 741 feature concentrations calculated based on 26 ISTDs and sample amounts of 4494 analy
#> i Concentrations are given in  mol/L.
```

The PCA plot of the normalized and quantified data revealed that most TQC samples clustered together. However, the TQCs from batch P-43 formed a distinct cluster in the lower right corner, and TQCs from the last four batches (P-01 to P-04) were also separated from the main TQC cluster. A separate cluster of BQCs, distinct from the main group in the top right corner, was composed primarily of samples from batches P-30 to P-38. These batches also contained BQCs that were identified as outliers in the PCA on raw intensities. This observation suggests that the normalization procedure was insufficient to correct for the observed differences in these BQC samples.

```r
plot_pca(
  data = mexp,
  variable = "conc",
  filter_data = FALSE,
  pca_dims = c(1,2),labels_column = "analysis_order",
  labels_threshold_mad = 4,
  qc_types = c("BQC", "TQC", "LTR", "NIST", "SPL"),
  ellipse_variable = "qc_type",
  log_transform = TRUE,
  point_size = 0.7, point_alpha = 0.7, font_base_size = 8, ellipse_alpha = 0.3,
  include_istd = FALSE,show_labels = TRUE,label_font_size = 1.5)
#> ! 93 features contained missing or non-numeric values and were exluded.
#> i The PCA was calculated based on `feature_conc` values of 741 features.
```

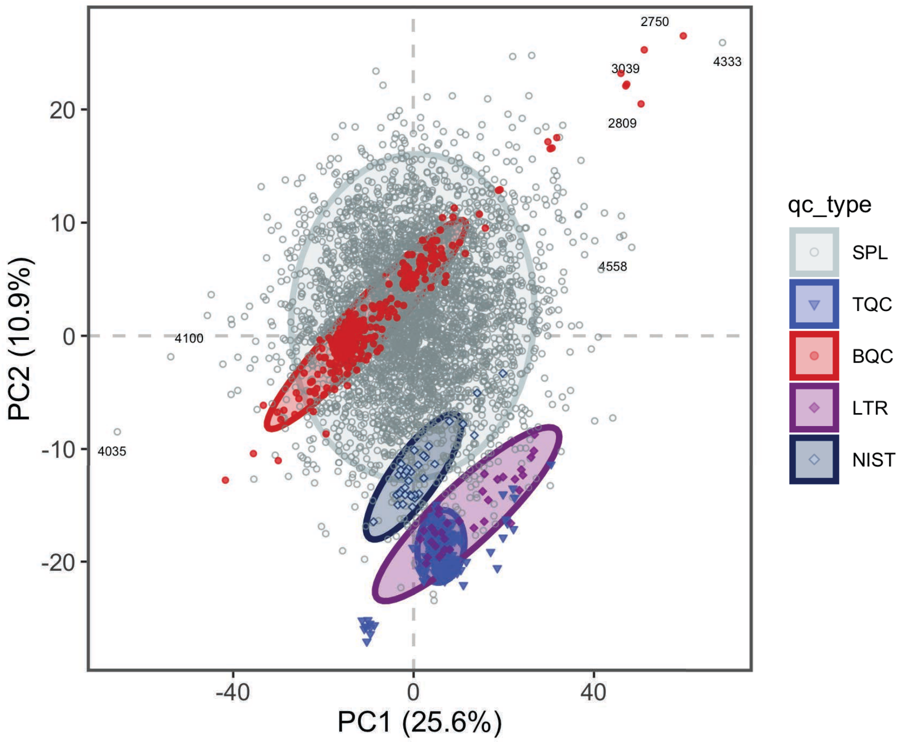

#### 3.10 Removal of batches with identified technical issues

Based on the runscatter plots, and the PCA of the concentrations, it became apparent that batches P-42 and P-43 exhibited technical issues. These batches, which consisted of re-run samples from previous batches, had problems that were not fully resolved by ISTD normalization. Therefore, these batches were excluded from further analysis.

```
ids_reruns <- mexp@annot_analyses |> filter(batch_id %in% c("P-43", "P-42")) |> pull(analysis

# ToDo: Currently exclude_analyses usage requires re-processing of the data. Needs to be upda
#mexp_final <- exclude_analyses(mexp, analyses = ids_reruns, clear_existing = FALSE)

# Workaround is to filter the dataset directly
mexp@dataset <- mexp@dataset |> filter(!analysis_id %in% ids_reruns)
mexp@annot_analyses <- mexp@annot_analyses |> filter(!analysis_id %in% ids_reruns)
mexp@annot_responsecurves <- mexp@annot_responsecurves |> filter(!analysis_id %in% ids_rerun
```

#### 3.11 Inspection of individual features concentrations across the run order

The calculated concentrations of individual features were plotted against the analysis order using RunScatter plots. In the examples shown below, some features did not show any significant drift or batch effects (e.g., SM 34:1). In contrast, clear drift effects within batches were visible for lyso-PC features, while other features exhibited batch effects (e.g., PC 36:4) or drifts that spanned across multiple batches (e.g., PI 38:4). The presence of these drift and batch effects indicates that ISTD normalization was insufficient to correct for such analytical artifacts. Therefore, further data correction is required to minimize these effects, as described in the next section..

```
plot_runscatter(mexp,
  variable = "conc", filter_data = FALSE,
  qc_types = c("SPL", "BQC", "TQC", "LTR"),
  include_feature_filter = "PC 34\\:2|PC 36\\:4|PE 38\\:4|PI 38\\:4|
    PC 38\\:6|CE 181\\:1|Cer d18\\:1\\/16\\:0|TG 52\\:3|SM 34\\:1|
    TG 56\\:3|LPC 18\\:1",
    #y_min = 0.00,
    #y_max = 0.15,
    #plot_range = c(0, 910),
    show_reference_lines = TRUE,ref_qc_types = "SPL",
    reference_fill_color = "#111111",
    reference_k_sd = 3,
    point_size = 0.5,
    point_border_width = 0.1,
    base_font_size = 6,
    cols_page = 2,
    rows_page = 6,
    cap_outliers = TRUE,
    reference_sd_shade = FALSE,
    #batch_zebra_stripe = TRUE,
    output_pdf = FALSE,
    path = "rt_all.pdf")
#> Generating plots (1 page)...
```

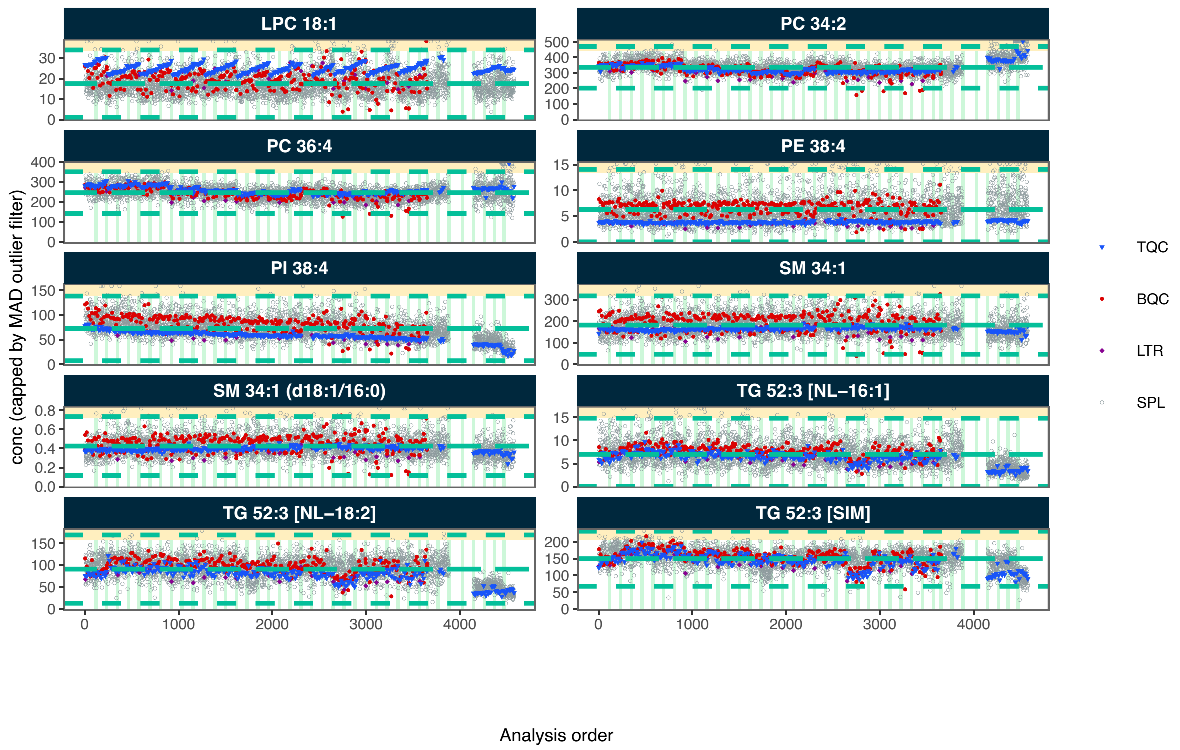

#### 3.12 Drift and Batch Correction

While QC-based drift correction is typically use for such metabolomics and lipidomics datasets, we concluded that the QC samples in this dataset did not fully and adequately represent the drifts observed in the study samples. This discrepancy may be due to the fact that the preparation of QC samples in such a large-scale analysis may not have been identical to that of the study samples due to logistical limitations, including differences in timing during sample extraction. We therefore opted for a sample-based Gaussian Kernel smoothing approach, assuming that the samples had been fully stratified and randomized. After the within-batch smoothing, the batches were re-aligned using sample-based median centering.

```
mexp <- correct_drift_gaussiankernel(
  mexp,
  variable = "conc",
  ref_qc_types = "SPL",
  batch_wise = TRUE,
  kernel_size = 10,
  outlier_filter = TRUE,
  outlier_ksd = 5,
  recalc_trend_after = TRUE,
```

```
  show_progress = FALSE)
#> i Applying `conc` drift correction...
#> i 29 feature(s) contain one or more zero or negative `conc` values. Verify your data or us
#> ! 54 features showed no variation in the study sample's original values across analyses.
#> ! 54 features have invalid values after smoothing. NA will be be returned for all values o
#> v Drift correction was applied to 687 of 741 features (batch-wise).
#> i The median CV change of all features in study samples was -1.38% (range: -15.44% to 4.47

mexp <- correct_batch_centering(
  mexp,
  ref_qc_types = "SPL",
  variable = "conc")
#> i Adding batch correction on top of `conc` drift-correction.
#> v Batch median-centering of 38 batches was applied to drift-corrected concentrations of al
#> i The median CV change of all features in study samples was -1.10% (range: -37.50% to 49.9
```

After applying the drift and batch correction, the RunScatter plots for the same features showed that the drift and batch effects observed in the study samples were minimized. However, because these corrections were based on the study samples, drift and batch effects appear to have been introduced into the QC samples. This is a further indicates that the QC samples did not adequately represent the analytical variations affecting the study samples.

```
plot_runscatter(mexp, variable = "conc",
  filter_data = FALSE,
  #include_feature_filter = "ISTD",
  qc_types = c("SPL", "BQC", "TQC", "LTR"),
  include_feature_filter = "PC 34\\:2|PC 36\\:4|PE 38\\:4|PI 38\\:4|
    PC 38\\:6|CE 181\\:1|Cer d18\\:1\\/16\\:0|TG 52\\:3|SM 34\\:1|
    TG 56\\:3|LPC 18\\:1",
    #y_min = 0.00,
    #y_max = 0.15,
    #plot_range = c(0, 910),
    show_reference_lines = TRUE,ref_qc_types = "SPL",
    reference_fill_color = "#111111",
    reference_k_sd = 3,
    point_size = 0.5,
    point_border_width = 0.1,
    base_font_size = 6,
    cols_page = 2,
    rows_page = 6,
    cap_outliers = TRUE,
    reference_sd_shade = FALSE,
```

```
    #batch_zebra_stripe = TRUE,
    output_pdf = FALSE,
    path = "rt_all.pdf")
#> Generating plots (1 page)...
```

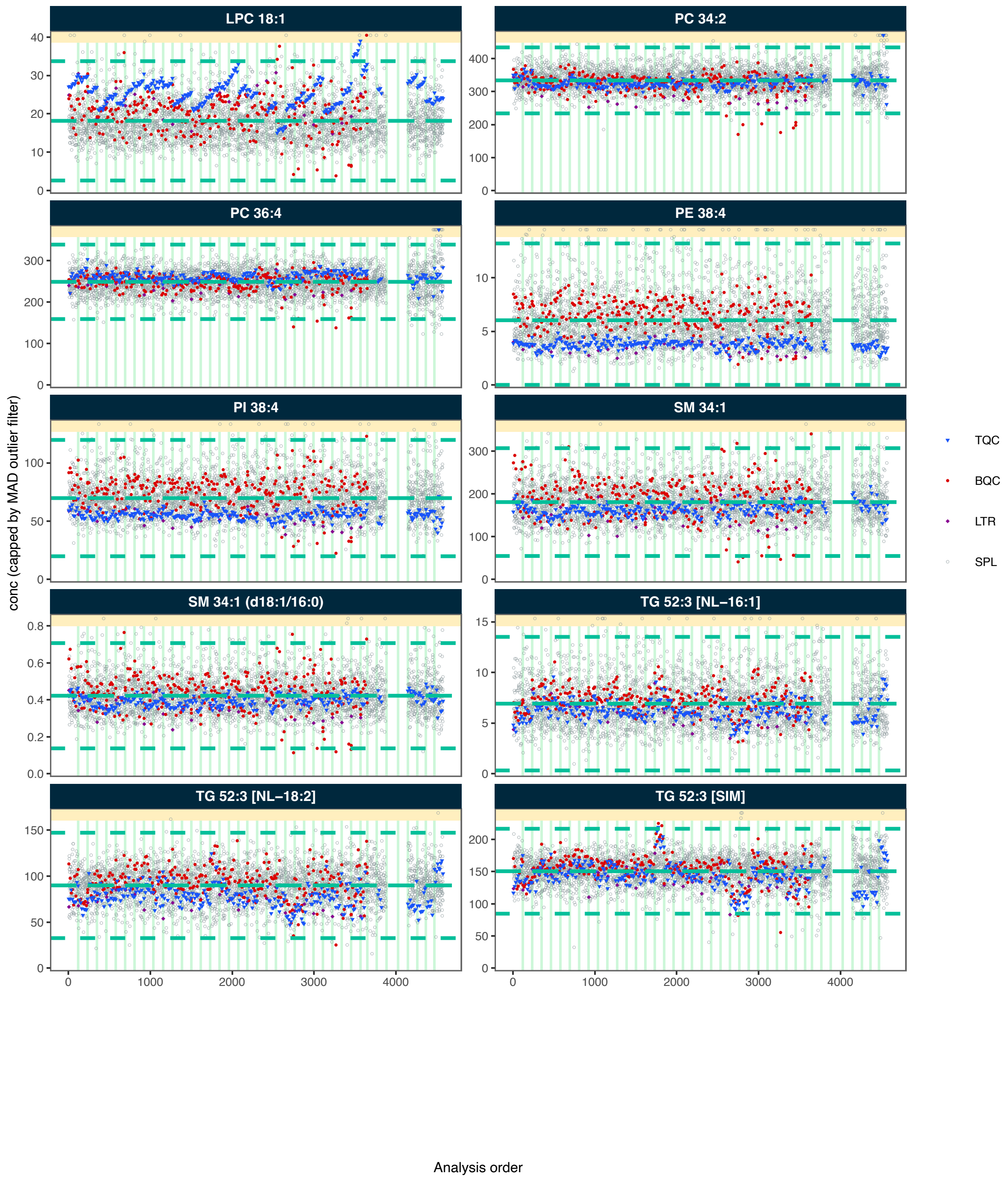

LPC 18:1
PC 34:2
PC 36:4
PE 38:4
PI 38:4
SM 34:1
SM 34:1 (d18:1/16:0)
TG 52:3 [NL−16:1]
TG 52:3 [NL−18:2]
TG 52:3 [SIM]
conc (capped by MAD outlier filter)
Analysis order
TQC
BQC
LTR
SPL

##### 3.13 RunScatter plots of final concentrations of all features

The code below generates PDF files with RunScatter plots for all features, showing the final concentrations after drift and batch correction. Run this code manually if you wish to generate these plots.

```
plot_runscatter(mexp, variable = "conc_raw",
                #include_feature_filter = "ISTD",
                qc_types = c("SPL", "BQC", "TQC", "LTR"),
                #include_feature_filter = example_species,
                #y_min = 0.00,
                #y_max = 0.9,
                #plot_range = c(0, 910),
                show_reference_lines = TRUE,
                ref_qc_types = "SPL",
                reference_fill_color = "#111111",
                reference_k_sd = 3,
                show_trend = TRUE,
                point_size = 1,
                base_font_size = 6,
                cols_page = 2,
                rows_page = 3,
                cap_outliers = FALSE,
                reference_sd_shade = FALSE,
                show_progress = FALSE,
                #batch_zebra_stripe = TRUE,
                output_pdf = TRUE,
                path = "./output/Dataset3_runscatter_rawConc_all.pdf")

plot_runscatter(mexp, variable = "conc",
                #include_feature_filter = "ISTD",
                qc_types = c("SPL", "BQC", "TQC", "LTR"),
                #include_feature_filter = example_species,
                #y_min = 0.00,
                #y_max = 0.9,
                #plot_range = c(0, 910),
                show_reference_lines = TRUE,
                ref_qc_types = "SPL",
                reference_fill_color = "#111111",
                reference_k_sd = 3,
                show_trend = TRUE,
                point_size = 1,
                base_font_size = 6,
```

```r
                 cols_page = 2,
                 rows_page = 3,
                 cap_outliers = FALSE,
                 show_progress = FALSE,
                 reference_sd_shade = FALSE,
                 #batch_zebra_stripe = TRUE,
                 output_pdf = TRUE,
                 path = "./output/Dataset3_runscatter_FinalConc_all.pdf")
```

##### 3.14 QC of normalization and drift/batch correction

Normalization with ISTDs, particularly the non-authentic, class-wide ISTDs used in this analysis, can introduce artifacts that can lead to an increase in sample variability rather than the expected reduction. Similarly, batch and drift correction can introduce artifacts and increase, rather than decrease, analytical variability. The following plot compares the variability of QC and study samples before (raw areas) and after normalization and drift/batch correction (final concentrations). The plot shows that for most QC types and study samples, variability was reduced after normalization and correction, indicating an overall effect of these processing steps. However, while the study samples exhibited decreased CVs after processing, the BQC and TQC samples showed increases. This may be another illustration that the QC samples did not fully represent the analytical variations affecting the study samples.

```r
mexp <- calc_qc_metrics(mexp, use_robust_cv = FALSE, use_batch_medians = TRUE)
```

```r
plot_normalization_qc(
  plot_type = "diff",
  data = mexp,
  before_norm_var = "intensity",
  after_norm_var = "conc",
  y_lim = c(-15,15),
  x_lim = c(0,75),
  qc_types = c("TQC", "BQC", "SPL", "NIST"),
  cols_page = 5,
  font_base_size = 5,
  point_size = 0.5,
  facet_by_class = TRUE,
  include_qualifier = FALSE,
)
```

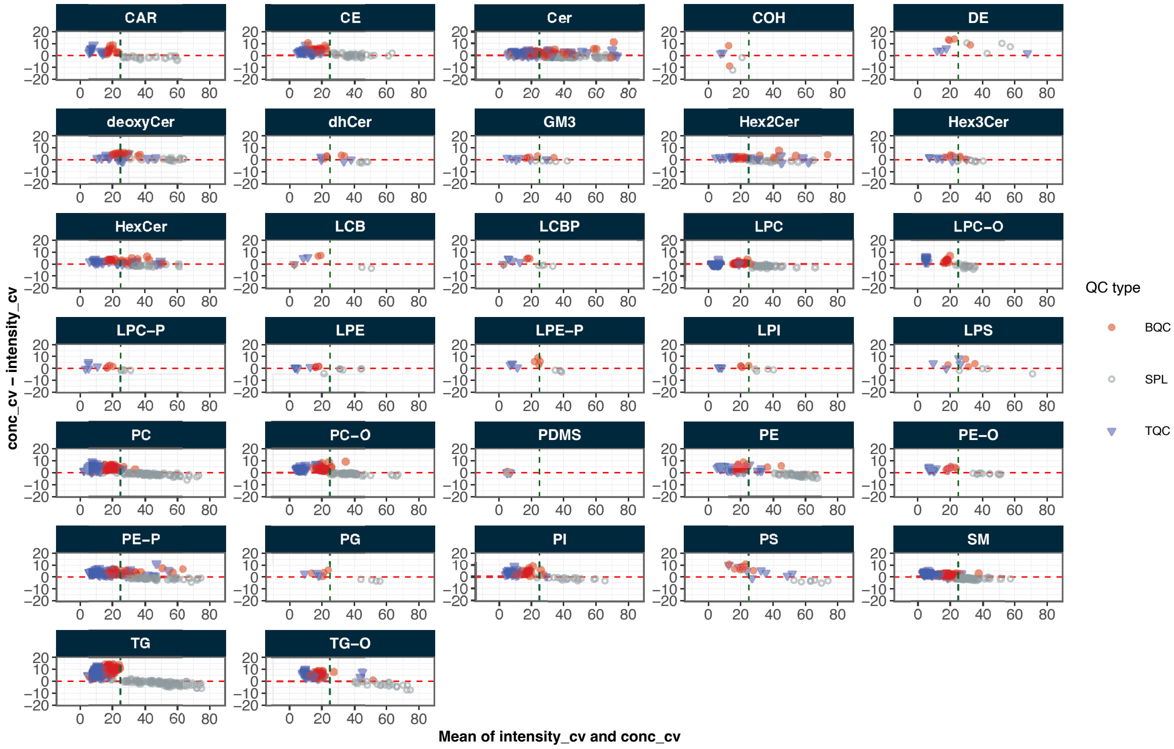

#### 3.15 Process vs instrument variability

To understand how much of the total technical variability comes from the overall process (sample preparation and the instrument variability) or just from the instrument, the CVs of a feature in batch(process) QC samples (a pooled sample repeatdly extracted and measured along with the study samples ) and the technical QC samples (a pooled extract measured at regula intervals) are compared. The plot below shows that for most features, the variability in the BQCs is higher than in the TQCs, indicating that sample preparation contributes considerable to the overall technical variability.

```
mrmhub::plot_qcmetrics_comparison(
  mexp,
  plot_type = "diff",
  y_shared = TRUE,
  "conc_cv_tqc",
  "conc_cv_bqc",
  log_scale = FALSE,
  equality_line = TRUE,
  facet_by_class = TRUE,
  point_size = 2,
```

```
  font_base_size = 5,
  x_lim = c(0, 25),
  y_lim = c(-15, 15))
```

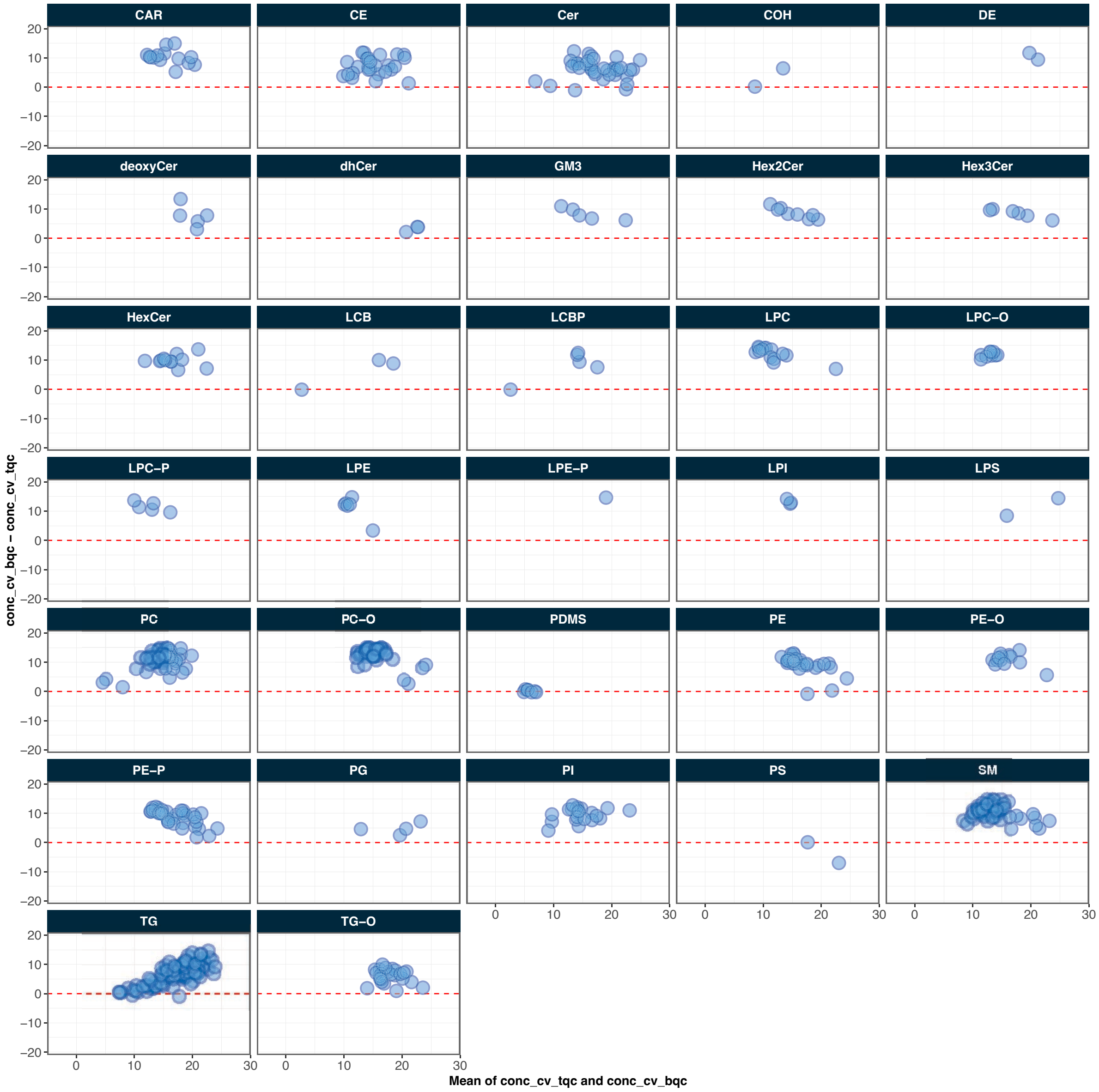

##### 3.16 Effect of feature intensity on technical variability

An analysis of the relationship between feature abundance and technical variability can be used to determine the intensity threshold at which feature signals exhibit increased noise and variance. This information can then be applied during peak picking to set appropriate intensity thresholds for feature detection and to reduce the processing time spent on integrating low-abundance features

```
mrmhub::plot_qcmetrics_comparison(
  mexp,
  plot_type = "scatter",
  y_shared = FALS,
  "intensity_median_bqc",
  "intensity_cv_bqc",
  log_scale = FALSE,
  equality_line = FALSE,
  facet_by_class = FALSE,
  point_size = 0.5,
  font_base_size = 5,
  x_lim = c(10, Inf),
  y_lim = c(0, 100))+
ggplot2::geom_smooth(method = "loess", se = FALSE, span = 0.75)+
geom_hline(yintercept = 20, linetype = "dashed", color = "grey70") +
scale_x_log10(expand = ggplot2::expansion(mult = c(0, 0.00)),
             breaks = c(1E2, 1E3, 1E4, 1E5, 1E6, 1E7, 1E8)) +
scale_y_continuous(expand = ggplot2::expansion(mult = c(0.03, 0.00)),breaks = c(0, 20, 40, 6
#> Scale for x is already present.
#> Adding another scale for x, which will replace the existing scale.
#> Scale for y is already present.
#> Adding another scale for y, which will replace the existing scale.
#> `geom_smooth()` using formula = 'y ~ x'
```

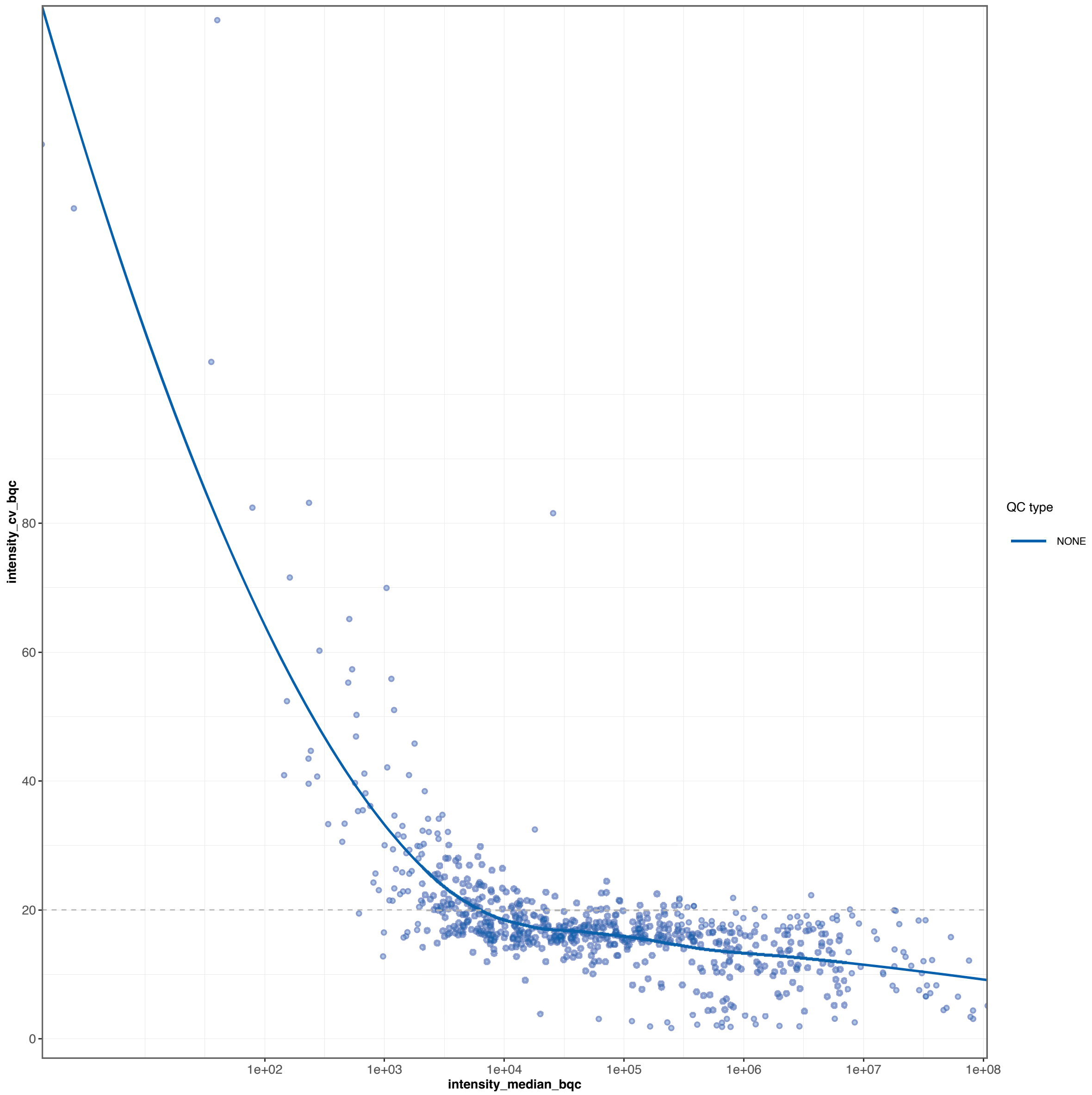

#### 3.17 Response curves

```
#| message: false

sel_species <- c("PC 26:0 (ISTD)", "SM 34:1", "PC 34:2", "CE 18:2", "TG 50:1 [NL-16:0]", "CE
plot_responsecurves(
```

```
  data = mexp,
  variable = "intensity",
  max_regression_value = 100,
  filter_data = FALSE,
  font_base_size = 6,
  line_width = 0.5, point_size = 1.2,
  include_feature_filter = sel_species,
  output_pdf = FALSE, path = "response-curves-dataset3.pdf",
  show_progress = FALSE,
  cols_page = 3,
  rows_page = 3,
  return_plots = TRUE)[[1]]+
  theme(panel.grid.minor = element_blank(),
        legend.position = "inside",
        legend.direction = "horizontal",
        legend.text = element_text(size = 6*0.6),
        legend.title = element_blank(),
        legend.key.size = unit(6 *0.6, "pt"),
        strip.text = element_text(size = 6),
        legend.position.inside = c(0.9, 0.01))
#> Registered S3 methods overwritten by 'ggpp':
#>   method                  from
#>   heightDetails.titleGrob ggplot2
#>   widthDetails.titleGrob  ggplot2
#> Generating plots (1 page)...  done!
```

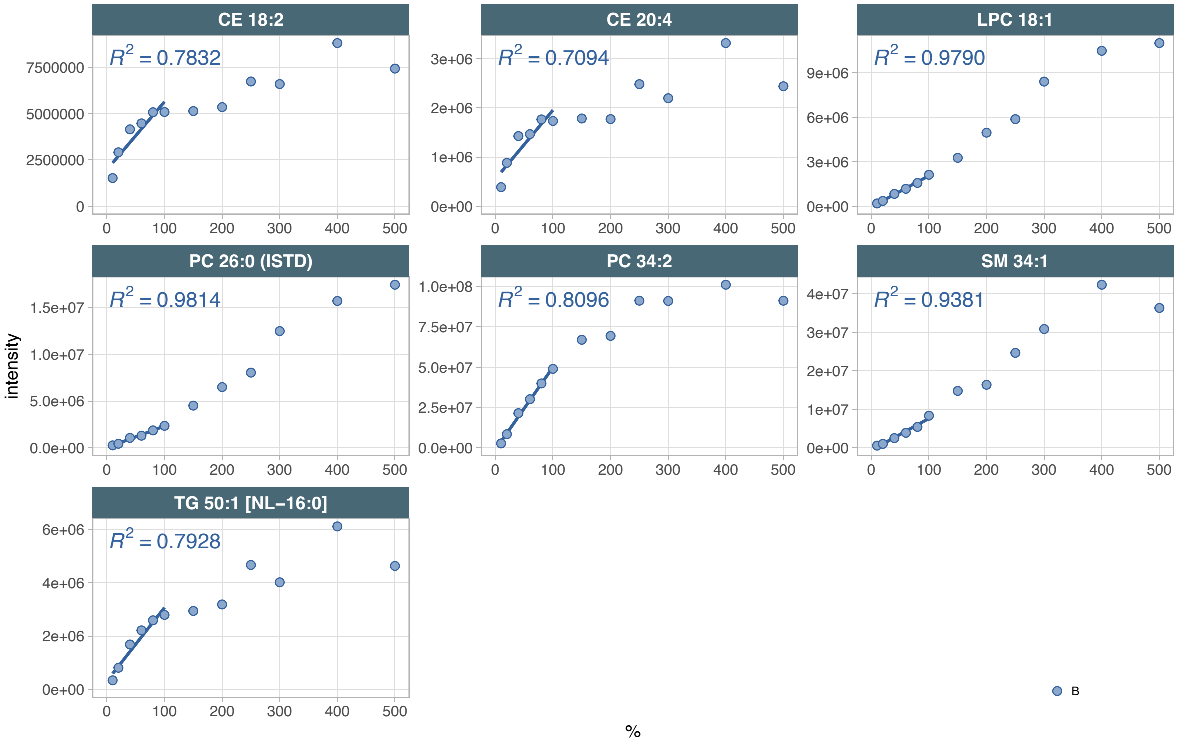

The code below generates PDF files with response curve plots for all features. Run this code manually if you wish to generate these plots.

```
plot_responsecurves(
  data = mexp,
  variable = "intensity", max_regression_value = 100,
  filter_data = FALSE,font_base_size = 6, line_width = 0.5, point_size = 1.2,
  output_pdf = TRUE, path = "./output/dataset3-response-curves.pdf",show_progress = FALSE,
  cols_page = 6, rows_page = 5,
  return_plots = FALSE)
```

#### 3.18 Feature filter

```
mexp <- calc_qc_metrics(mexp, use_robust_cv = FALSE, use_batch_medians = TRUE, include_respo
```

```
mexp  <- filter_features_qc(
  data =  mexp,
  clear_existing = TRUE,
  use_batch_medians = TRUE,
```

```r
  include_qualifier = FALSE,
  include_istd = FALSE,
  response.curves.selection = 1,
  response.curves.summary = "mean",
  min.rsquare.response = 0.8,
  min.slope.response = 0.5,
  max.yintercept.response = 0.5,
  min.signalblank.median.spl.pblk = 10,
  min.intensity.median.spl = 100,
  max.cv.conc.bqc = 25,
  #max.dratio.sd.conc.bqc = 0.75,
  max.prop.missing.conc.spl = 100,
  features.to.keep = c("CE 20:4", "CE 22:5", "CE 22:6", "CE 16:0", "CE 18:0")
)
#> ! The following features were forced to be retained despite not meeting filtering criteri
#> v
New feature QC filters were defined: 504 of 741 quantifier features meet QC criteria (not in
```

#### 3.19 Feature filter Results

```r
plot_qc_summary_byclass(mexp) +
    theme(
      strip.text = ggplot2::element_text(size = 5),
      #aspect.ratio = 0.9,
      legend.position = "inside",
      axis.text = element_text(size = 7),
      axis.title = element_text(size = 8, face = "plain"),
      axis.text.y.right =  element_text(size = 7, face = "plain"),
      legend.direction = "vertical",
      legend.text = element_text(size = 8*0.7),
      legend.title = element_blank(),
      legend.key.size = unit(8 *0.7, "pt"),
      legend.position.inside = c(0.77, 0.27)
)
```

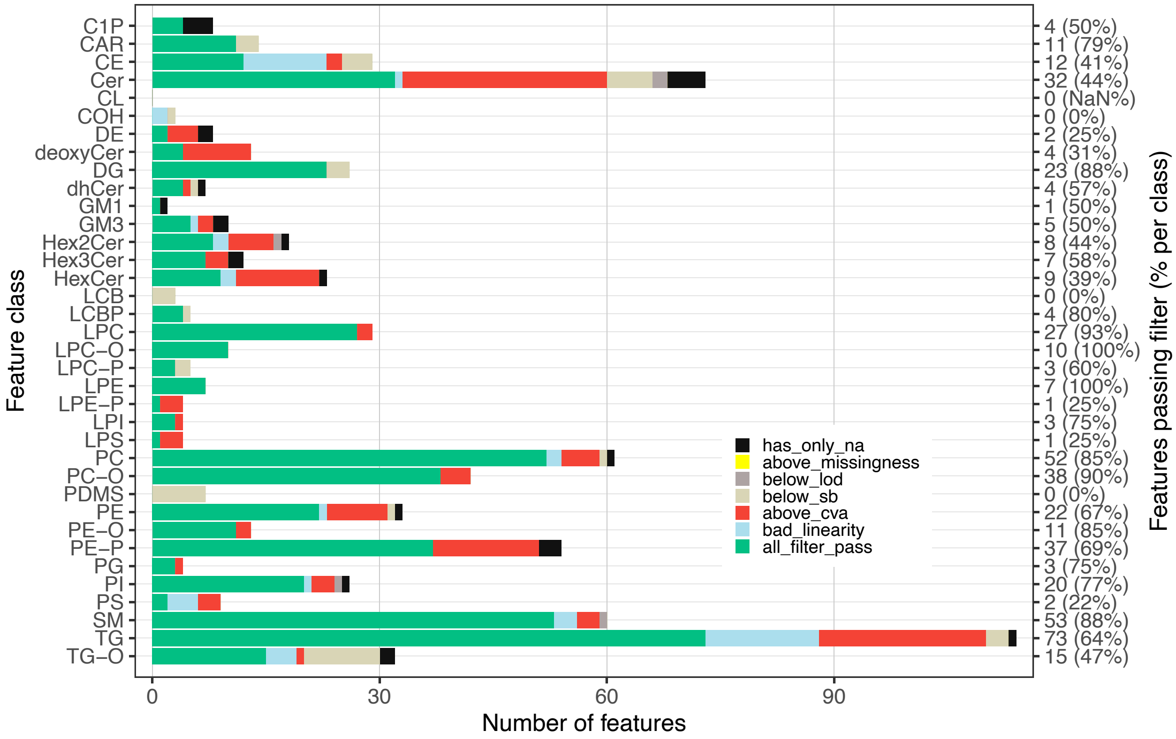

##### 3.20 Feature filter Venn

```
plot_qc_summary_overall(mexp)
```

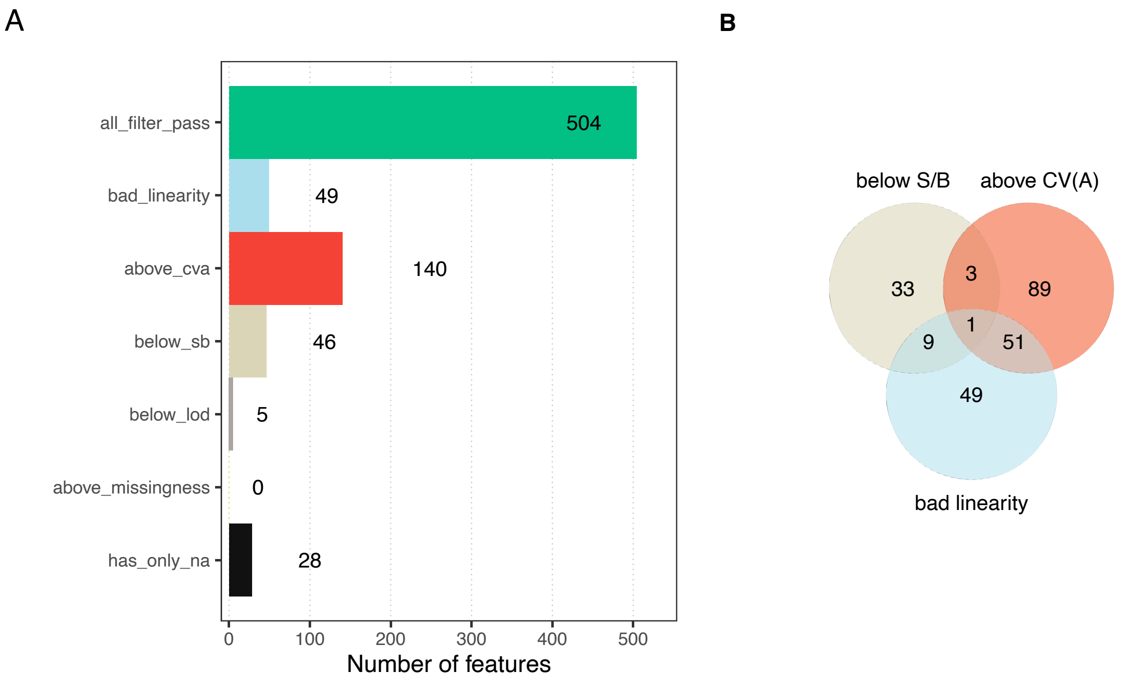

#### 3.21 PCA plots of the final dataset

The PCA plot shows that the overall structure remains similar to that observed before correction. The TQCs are more dispersed, which could reflect the fact that they capture only instrument variability, whereas the sample-based drift correction may have artificially increased the apparent TQC variability. Nevertheless, the TQCs still form a relatively compact cluster. In contrast, the BQCs, LTR, and NIST exhibit long elliptical clusters spanning a larger area. This pattern suggests that sample processing may have introduced systematic variability that was not fully corrected by ISTD normalization and drift/batch correction.

```
plot_pca(
  data = mexp,
  variable = "conc",
  filter_data = FALSE,
  pca_dims = c(1,2),labels_column = "analysis_order",
  labels_threshold_mad = 5,
  qc_types = c("BQC", "TQC", "LTR", "NIST", "SPL"),
  ellipse_variable = "qc_type",
  log_transform = TRUE,
  point_size = 0.7, point_alpha = 0.7, font_base_size = 8, ellipse_alpha = 0.3,
```

```
  include_istd = FALSE,show_labels = TRUE,label_font_size = 1.5)
#> ! 91 features contained missing or non-numeric values and were exluded.
#> i The PCA was calculated based on `feature_conc` values of 741 features.
```

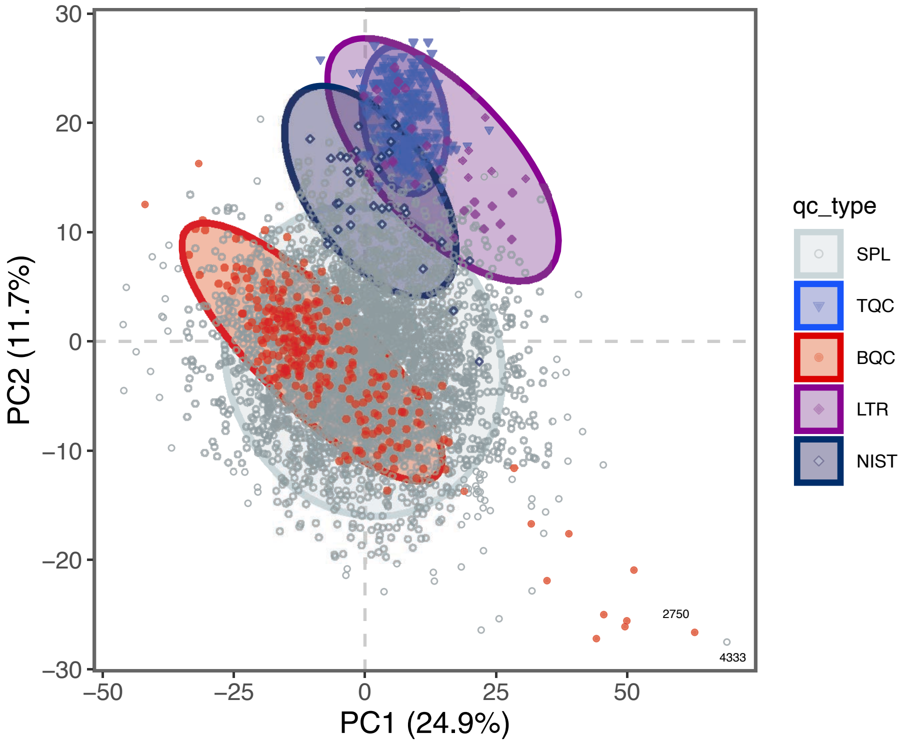

##### 3.22 Lipidome Profile

As a final overview, the feature concentration profile of the filtered dataset is shown below. Validating the concentrations, for example, those of the most abundant species, the summed concentrations per lipid class, or the ratios between lipid classes, against in-house reference values or literature data helps ensure that the quantification is within expected ranges and that no major quantification errors are present.

```
plot_abundanceprofile(
  data = mexp,
  log_scale = TRUE,
  filter_data = TRUE,
  variable = "conc",
  qc_types = "SPL",
  #x_lim = c(-6, 2),
```

```
  x_label = NA,
  feature_map = "lipidomics")
```

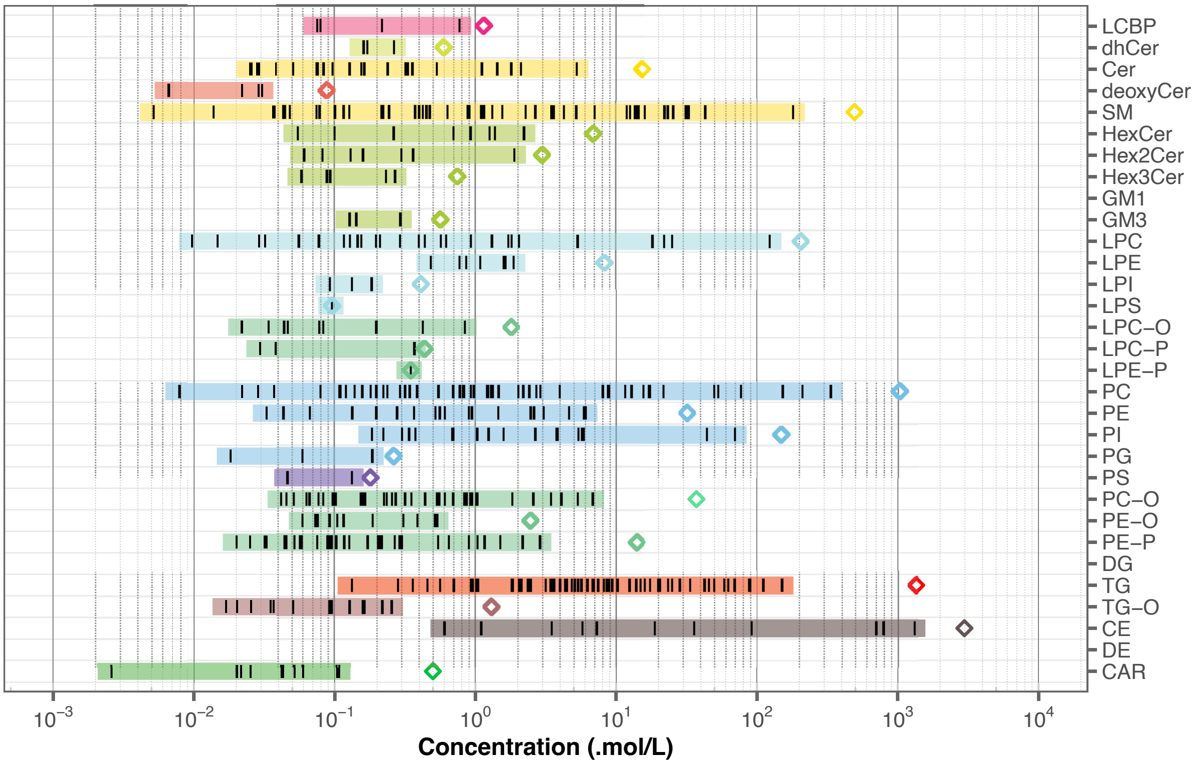

#### 3.23 Export final dataset

The final dataset is exported as a CSV file, containing the final concentrations of the filtered dataset. The user can specify whether to include qualifier features, and whether to filter the data for specific QC types. Here, only the study samples (SPL) samples are included.

```
save_dataset_csv(
  data = mexp,
  path = "./output/sperfect_UNFILTERED_RAW-feature_conc_uM_20250518a.csv",
  variable = "conc",
  qc_types = "SPL",
  include_qualifier = FALSE,
  filter_data = FALSE)
```
